# Genome-Enabled Discovery of Anthraquinone Biosynthesis in *Senna tora*

**DOI:** 10.1101/2020.04.27.063495

**Authors:** Sang-Ho Kang, Ramesh Prasad Pandey, Chang-Muk Lee, Joon-Soo Sim, Jin-Tae Jeong, Beom-Soon Choi, Myunghee Jung, So Youn Won, Tae-Jin Oh, Yeisoo Yu, Nam-Hoon Kim, Ok Ran Lee, Tae-Ho Lee, Puspalata Bashyal, Tae-Su Kim, Chang-Kug Kim, Jung Sun Kim, Byoung Ohg Ahn, Seung Yon Rhee, Jae Kyung Sohng

## Abstract

*Senna tora* is a widely used medicinal plant. Its health benefits have been attributed to the large quantity of anthraquinones, but how they are made in plants remains a mystery. To identify the genes responsible for plant anthraquinone biosynthesis, we sequenced and annotated the genome of *S. tora* at the chromosome level with contig N50 and super-scaffold N50 of 4.03 Mb and 41.7 Mb. Comparison among related plant species showed that a chalcone synthase-like (CHS-L) gene family has lineage-specifically and rapidly expanded in *S. tora*. Combining genomics, transcriptomics, metabolomics, and biochemistry, we identified a CHS-L responsible for biosynthesis of anthraquinones, the first example in plants. The *S. tora* reference genome will accelerate the discovery of biologically active anthraquinone biosynthesis pathways in medicinal plants.

**One Sentence Summary:** The chromosome-scale reference genome of a medicinal plant *Senna tora*, transcriptomics, metabolomics, and biochemical analysis provide new insights into anthraquinone biosynthesis in plants.

## Main Text

*Senna tora* (L.) Roxb., also known as *Cassia tora*, is a favorite of ancient Chinese and Ayurvedic herbal medicine that is now widely used around the world today (*1*). Recent studies point to *S. tora*’s beneficial activities against microbial (*2-5*) and parasitic (*6*) infections, prevention or delay of the onset of neurodegenerative diseases (*7, 8*), and diabetes (*9*). *S. tora*’s positive health impact is attributed to the significant amount of anthraquinones in mature seeds and other parts of the plant (*10-12*). Anthraquinones are aromatic polyketides made by bacteria, fungi, insects, and plants (*13, 14*). Besides their medicinal benefits, natural anthraquinones are garnering attention as alternatives to synthetic dyes that damage aquatic ecosystems (*15-17*). Bacteria, fungi, and insects make anthraquinones via a polyketide pathway using type I polyketide synthases (*13, 18*).

For plants, how anthraquinones are made remains unknown. Two biosynthesis pathways have been proposed for anthraquinones in plants: 1) a polyketide pathway (*19*) and 2) a combination of shikimate and mevalonate/methyl-D-erythritol 4-phosphate pathways (*20, 21*). More than three decades ago, radiolabeled feeding experiments indicated that the A and B rings of anthraquinones were derived from shikimate and *α*-ketoglutarate via *O*-succinylbenzoate (*22-24*) and C-ring from mevalonate pathway via isopentenyl pyrophosphate (IPP) and dimethylallyl pyrophosphate (DMAPP) (*20, 21, 25*) or 2-C-methyl-D-erythritol 4-phosphate (MEP) pathway (*26, 27*). Contrarily, recent studies speculated biosynthesis of anthraquinones in plants to occur via a polyketide pathway (*28-30*). Type III polyketide synthase (PKS) enzymes could actively catalyze seven successive decarboxylative condensations of malonyl-CoA to produce octaketide chain (*29, 30*). The linear polyketide chain undergoes cyclization and decarboxylation reactions to produce the core unit of polyketides such as atrochrysone carboxylic acid followed by decarboxylation to atrochrysone and dehydration to emodin anthrone (**Fig. S1**) (*28-32*). However, to date, no study in type III PKS enzymes has provided conclusive evidence on the biosynthesis of anthraquinones or the intermediate metabolites of the pathways. Beerhues and colleagues (*28*) showed promising outcomes on the biosynthesis of an anthranoid scaffold via the polyketide pathway. The *in vitro* reaction using acetyl-CoA, stable carbon isotope labeled malonyl-CoA, and cell-free extracts of *Cassia bicapsularis* cell cultures produced emodin anthrone and *O*-methylated torochrysone (*28*). However, this study could not discern whether a PKS was involved in the biosynthesis of the anthranoid scaffolds.

Despite the extensive applications of *S. tora* in medicine and industry (*1*), there has been little report of molecular and genomic studies of this remarkable plant. Elucidating the genes responsible for biosynthesis of anthraquinones in *S. tora* will aid molecular breeding and development of tools for probing its biochemistry. As a first step, we present here a high-quality reference genome sequence of *S. tora* cultivar Myeongyun, which allowed us to examine the evolution of candidate gene families involved in anthraquinone biosynthesis and identify the first enzyme known to catalyze a plant anthraquinone. By combining genomic, transcriptomic, metabolomic, and biochemical approaches, we systematically screened, identified, and confirmed the key gene responsible for biosynthesis of an anthraquinone scaffold in *S. tora*.

We generated one of the highest quality genomes for medicinal plants using a combination of approaches. The *S. tora* cultivar Myeongyun genome was assembled with Illumina paired-end and mate-pair as well as Pacific Biosciences long-read sequencing (**Tables S1 and S2**). *S. tora* has an estimated genome size of ∼547 Mb based on k-mer analysis (**Fig. S2**). Through chromosome conformation capture (Hi-C) mapping, we generated 13 chromosome-scale scaffolds (hereafter called chromosomes; Chr1-Chr13) totaling 502.6 Mb, 95.5% of the ∼526.4 Mb of the assembled genome (**Table 1 and Figs. S3 and S4**). We evaluated the quality of assembly using Benchmarking Universal Single Copy Orthologs (BUSCO) (*33*), sequencing of 10 bacterial artificial chromosome (BAC) clones, and comparing to a linkage map (**Fig. S5**). BUSCO estimates 94.3% completeness (**Table S3**) suggesting that the assembly includes most of the *S. tora* gene space. BAC sequence alignments showed high mapping rates (99.8%) with the assemblies (**Fig. S6 and Table S4**). We also built a genetic map of diploid *S. tora*, to which 401.1 Mb of the assembled scafoolds were mapped (**Fig. S7**). The 13 linkage groups matched well to the 13 chromosomes, indicating high quality of *S. tora* genome assembly (**Fig. S7**).

**Table 1.** Summary of genome assembly and protein-coding genes in *S. tora*.

|  | Contigs | Superscaffolds |
| --- | --- | --- |
| **Assembly features** |  |  |
| Numbers | 732 | 13 |
| Total length | ~526.4 Mb | ~502.6 Mb |
| N50 | ~4.03 Mb | ~41.7 Mb |
| Longest | ~14.9 Mb | ~52.7 Mb |
| Coverage | ~96.23% | ~95.5% |
| GC content | ~35.45% |  |
| **Protein-coding genes** |  |  |
| No. of genes | 45,268 |  |
| Mean gene length | 3,162 bp |  |
| Gene coverage | ~27.15% |  |
| Mean exon length | ~217 bp |  |
| Exon coverage | ~8.11% |  |
| Mean intron length | ~656 bp |  |
| Intron coverage | ~18.92% |  |

*S. tora*’s genomic content is consistent with other sequenced plant genomes. A total of 45,268 genes were annotated with the average gene length (3,157 bp), exon sequence length (217 bp with 4.37 exons per gene), and intron length (655 bp) that were similar to those of other legume species (**Table 1 and Fig. S8**). Among the protein-coding genes, 31,010 (68.50%) showed homology to characterized genes based on BLAST searches and 25,453 (56.23%) and 17,450 (38.55%) were assigned to Gene Ontology (GO) terms and KEGG pathways, respectively (**Table S5**). As expected, the genes were unevenly distributed with an increase in density towards the ends of the pseudomolecules (**Fig. 1A**). We also identified genes encoding for 839 tRNA, 752 rRNA, 3,278 long non-coding RNA (lncRNA) (**Fig. 1A and Tables S6 and S7**), and 1,644 transcription factors (TFs) from 36 families that accounted for 3.63% of the protein-coding genes (**Fig. S9 and Table S8**).

**Fig. 1.**
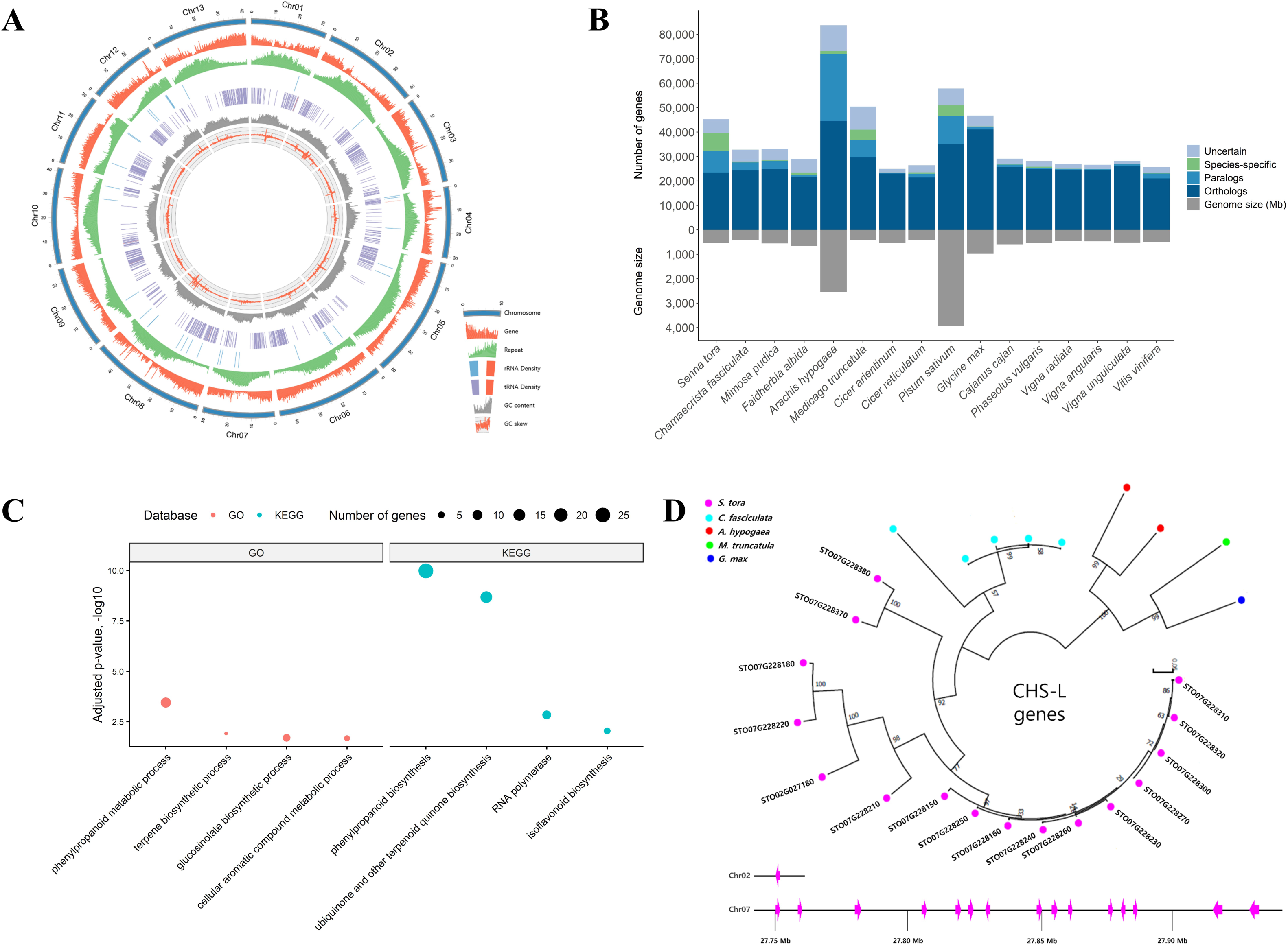
The *S. tora* genome and comparative genomic analysis. **(A)** The landscape of genome assembly (∼502 Mb) and annotation of *S. tora*. Tracks (from outside) correspond to chromosomes (Chr01 to Chr13 on a Mb scale), gene density, repeat density, rRNA density, tRNA density, GC content and GC skew. Tracks are drawn in non-overlapping 100 kb sliding windows. The red bars in the rRNA and tRNA tracks represent the maximum density of copies on the scale. **(B)** An overview of orthologous and paralogous genes among *S. tora*, related legumes, and *V. vinifera*. “Uncertain” indicates homologous genes obtained from BLAST but not found using OrthoMCL. “Species-specific” genes do not have any similarity to genes in the other species based on BLAST and OrthoMCL. **(C)** Significantly enriched biological process GO categories in secondary metabolism and all enriched KEGG categories of gene families that expanded specifically in *S. tora* compared to the other 15 species. **(D)** Lineage-specific expansion of the CHS-L gene family in *S. tora* and four other legumes. The 15 tandemly duplicated gene clusters are ordered and shown on chromosome 7, as well as one gene on chromosome 2.

To assess the evolution of candidate gene families involved in anthraquinone biosynthesis, we compared the *S. tora* genome with those from 15 related plant species. Reciprocal pair-wise comparisons (*34*) of the 16 species (15 legumes and grape vine) revealed that *S. tora* has the most number species-specific genes of all the 16 plants compared, with 7,231 (15.9%) genes that are specific to *S. tora* (**Figs. 1B, S10, and Table S9**). We compared gene family expansion and contraction across the species to identify gene families that were expanded or contracted specifically in *S. tora*. Of the 36,746 gene families found among the sixteen species, 3,076 and 3,659 gene families were expanded and contracted specifically in *S. tora*, respectively (**Fig. S11**). The gene families that were specifically expanded in *S. tora* were enriched for several Gene Ontology (GO) and KEGG terms, including those involved in specialized metabolism including ‘phenylpropanoid biosynthesis’, ‘isoflavoniod biosynthesis’, and ‘terpene biosynthesis’, likely reflecting the importance of genes for the biosynthesis of phenolics, isoflavonoids, and terpenoids in *S. tora* (**Fig. 1C and Table S10**).

We next probed which of the lineage-specifically expanded families might be involved in anthraquinone biosynthesis. In plants, type III polyketide synthases such as chalcone synthases (CHSs) are involved in the biosynthesis of plant secondary metabolites, particularly acetate-pathway derived flavonoids, stilbenes, and aromatic polyphenols (*28, 35, 36*). The *S. tora* CHS family contains twelve CHS and sixteen CHS-L genes (**Fig. S12**). Interestingly, the CHS-L gene family specifically and rapidly expanded only in the *S. tora* genome (16 genes in *S. tora*, 5 in *C. fasciculata*, 2 in *A. hypogaea*, 1 in *M. truncatula*, 1 in *G. max*, and none in the other 11 species) (**Fig. 1D and Table S11**). Twelve of the CHS-L genes are specific to the *S. tora* lineage and the majority of *S. tora* CHS-L genes (15 of 16) are distributed only in chromosome 7 and arranged in tandem (**Fig. 1D**). Interestingly, CHS genes have been contracted in the *S. tora* genome (**Table S11**). Even though carminic acid (C-glucosylated anthraquinone) was produced by combining an octaketide synthase gene from *Aloe arborescens*, two cyclases from *Streptomyces*, and a glycosyltransferase from an insect in *Nicotiana* plants (*37*), a direct evidence of anthraquinone biosynthesis using plant CHS enzymes has not been established so far. However, several studies speculated involvement of CHS-Ls in synthesizing anthraquinones (*28, 29, 37*). At the genomic level, we found that the CHS-L gene family has been expanded most notably in *S. tora*, which may explain in part why *S. tora* is rich in anthraquinones.

To test the hypothesis that CHS-Ls might be involved in anthraquinone biosynthesis in *S. tora*, we turned to the issue that is enriched in anthraquinones, the seed. We profiled anthranoids from seven developmental stages of the seed (**Fig. 2A**), using ten standard anthraquinones (**Table S12**) as references for quantification. Anthraquinone accumulation varied in each stage (**Fig. 2B**). Importantly, the profile shifted towards modified derivatives such as glucoaurantio-obtusin, aurantio-obtusin, obtusifolin, and chryso-obtusin during late stages of seed development (**Fig. 2B and Table S13**) essentially becoming major storage metabolites in dry seeds.

**Fig. 2.**
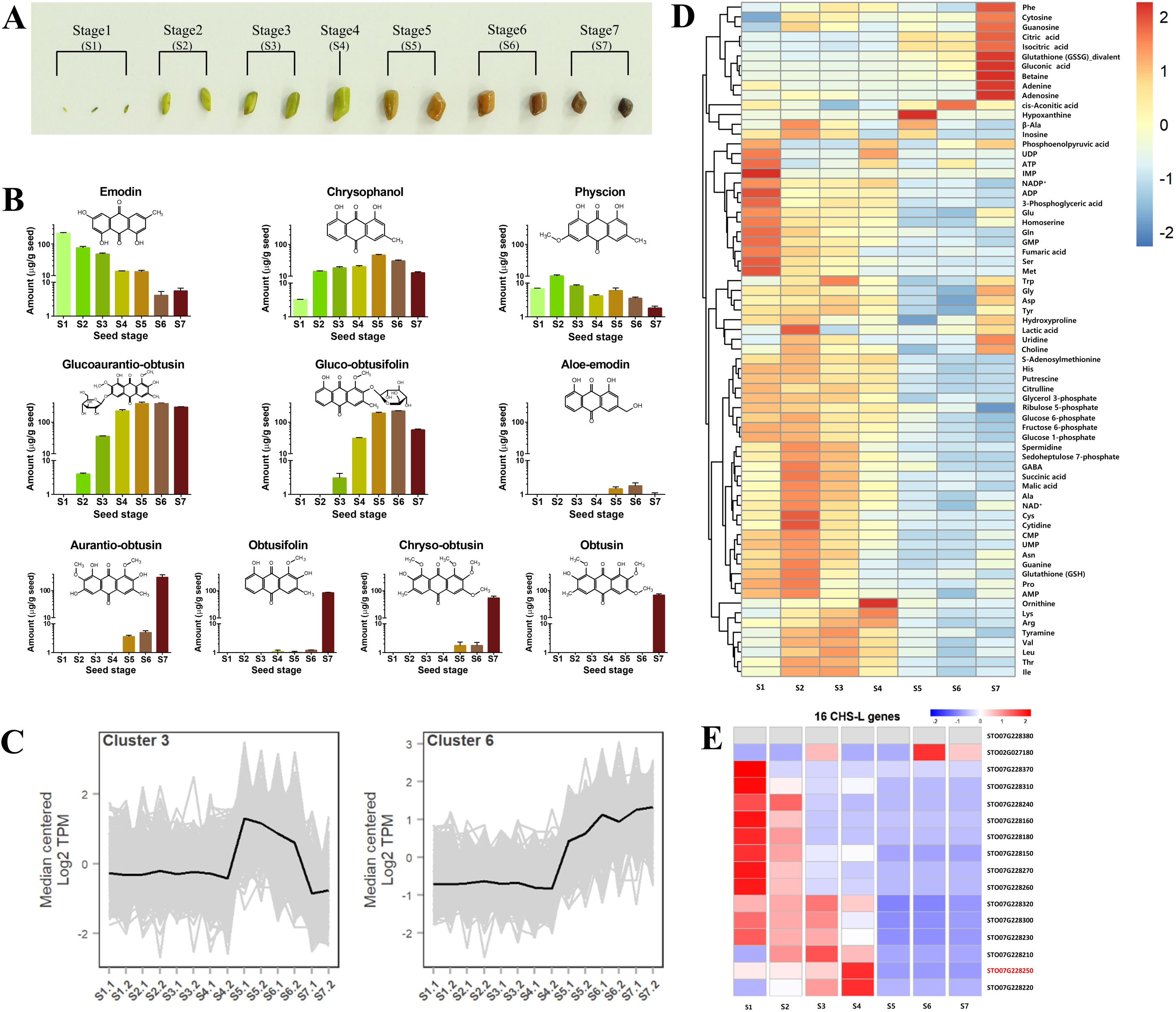
Analysis of anthraquinone contents, primary metabolites and CHS-L gene expression during *S. tora* seed development. **(A)** Developmental progression of *S. tora* seeds (Stage1-Stage7). **(B)** Concentrations of 10 anthraquinones during the 7 developmental stages of *S. tora* seeds (mean ± SD, n = 3). **(C)** Scaled transcript expression profiles (in transcripts per million, TPM) of cluster 3 and cluster 6 during seed development. **(D)** Quantitative estimation of 69 primary metabolites during seed development. Heatmap indicates normalized concentration from three biological replicates. **(E)** Expression analysis of CHS-L genes during seed development. Heatmap represents normalized transcripts per million (TPM) from two biological replicates. S1-S7 represents seed development stages of *S. tora*.

To identify genes involved in the biosynthesis of anthraquinones during seed development, we performed transcriptome and metabolome analysis from developing seeds. A total of 13,488 genes were differentially expressed relative to stage 1 during seed development. Co-expression analysis of differentially expressed genes during seed development detected nine co-expression clusters (**Fig. S13**). Among them, cluster 3 and cluster 6 showed similar patterns to anthraquinone accumulation in which genes were highly induced starting stage 5 (**Fig 2C**). Cluster 6 was statistically overrepresented with genes annotated as transferases, UDP-glycosyltransferases, and oxidoreductases, which may reflect enzymes involved in the tailoring of anthraquinones to produce gluco-obtusifolin, glucoaurantio-obtusin, and other derivatives including aurantio-obtusin (**Fig. 2B and Table S14**). The transcriptome and anthraquinone quantification suggested that there may be a metabolic switch from primary to secondary metabolism between stages 4 and 5. For example, Zhang and colleagues showed that *Streptomyces* underwent a metabolic switch from primary metabolism to polyketide biosynthesis during the transition from exponential to stationary phase (*38*). To test whether *S. tora* seeds underwent a similar metabolic switch, we profiled 178 selected primary metabolites of primary metabolic pathways including central carbon, urea cycle, lipid, amino acid, and nucleotide metabolism during the seven seed developmental stages (**Fig. S14 and Table S15**). We observed a clear metabolic switch between stages 4 and 5 where the majority of the primary metabolites were reduced after stage 4 (**Fig. 2D and Table S16**). With these data in hand, we searched specifically for CHS-L genes that were induced in stage 4 when the primary metabolite levels become reduced and anthraquinones start to accumulate. Among the 16 CHS-L genes, two genes (STO07G228250 and STO07G228220) showed high expression levels at stage 4 (**Fig. 2E**) where anthraquinone contents started to accumulate in seeds (**Table S13**). Both of these genes share high amino acid sequence similarities with each other and with STO02G027180, the gene that was highly expressed at stage 6 of seed development (**Fig. 2E**). Further comparison of sequence alignment and phylogenetic tree analysis with previously characterized octaketide synthases, HpPKS and ArOKS, which were presumed to be involved in hypericin and barbaloin biosynthesis based on the production of the octaketide shunt products (*30, 32*), showed that STO07G228250 was more similar to them than the other two CHS-Ls (**Fig. S15**). Therefore, we hypothesized that STO07G228250 could be engaged in anthraquinones biosynthesis in *S. tora*. As a control, we also selected a member of the CHS family, STO03G058250 (**Fig. S15**), hypothesized to be involved in flavonoid biosynthesis, for biochemistry experiments.

To perform enzymatic assays, we expressed STO07G228250 (CHS-L) and STO03G058250 (CHS) heterologously in *E. coli* and purified them to homogeneity (**Fig. S16**). Enzyme assays were conducted in a phosphate buffer saline containing malonyl-CoA for successive condensation reactions to produce polyketides. STO07G228250 (CHS-L)-catalyzed reaction mixture revealed the existence of two molecules with a molecular mass of 319.08 Da and 301.07 Da (**Fig. 3**). Neither of these metabolites was detected in reactions containing STO03G058250 (CHS) or heat-denatured STO07G228250 (CHS-L) (control), indicating that these two masses are most likely the products of the PKS-catalyzed reaction (**Fig. 3**). The ESI-MS spectrum showed a compound with a distinct peak at *m/z*^+^ 319.0827 (retention time (*t*_R_) 4.45 min), which corresponds exactly to the mass of atrochrysome carboxylic acid (C_16_H_14_O_7_ with 319.0818 Da in the proton adduct mode). Furthermore, the theoretical isotope model for the same chemical formula corroborated perfectly to the observed isotope mass (**Fig. S17**). Likewise, the ESI-MS spectrum of the latter metabolite (*t*_R_ 4.62 min) *m/z*^+^ 301.0715 matched to the mass of endocrocin anthrone (C_16_H_12_O_6_ with calculated exact mass of 301.0712 Da) for which the theoretical isotope mass model and observed mass isotope were perfectly aligned (**Fig. S17**). To further verify these metabolites as PKS-derived products, a set of reactions were conducted with STO07G228250 (CHS-L) and heat-denatured STO07G228250 (dead CHS-L) containing ^13^C_3_-malonyl-CoA as substrate. The EIC for all carbon labeled atrochrysome carboxylic acid (^13^C_16_H_14_O_7_, exact mass: 335.1355 Da) (**Fig. 3 and Fig. S17**) and endocrocin anthrone (^13^C_16_H_12_O_6_, exact mass: 317.1249 Da) (**Fig. 3 and Fig. S17**) were confirmed to be present in only the reaction with CHS-L. The observed ESI-MS spectra aligned to the theoretical isotope mass of corresponding metabolites. Except for these two metabolites, none of the other octaketide metabolites such as emodin anthrone, endocrocin, emodin, chrysophanol, or islandicin was produced even when NADPH was added in the reaction mixtures (data not shown).

**Fig. 3.**
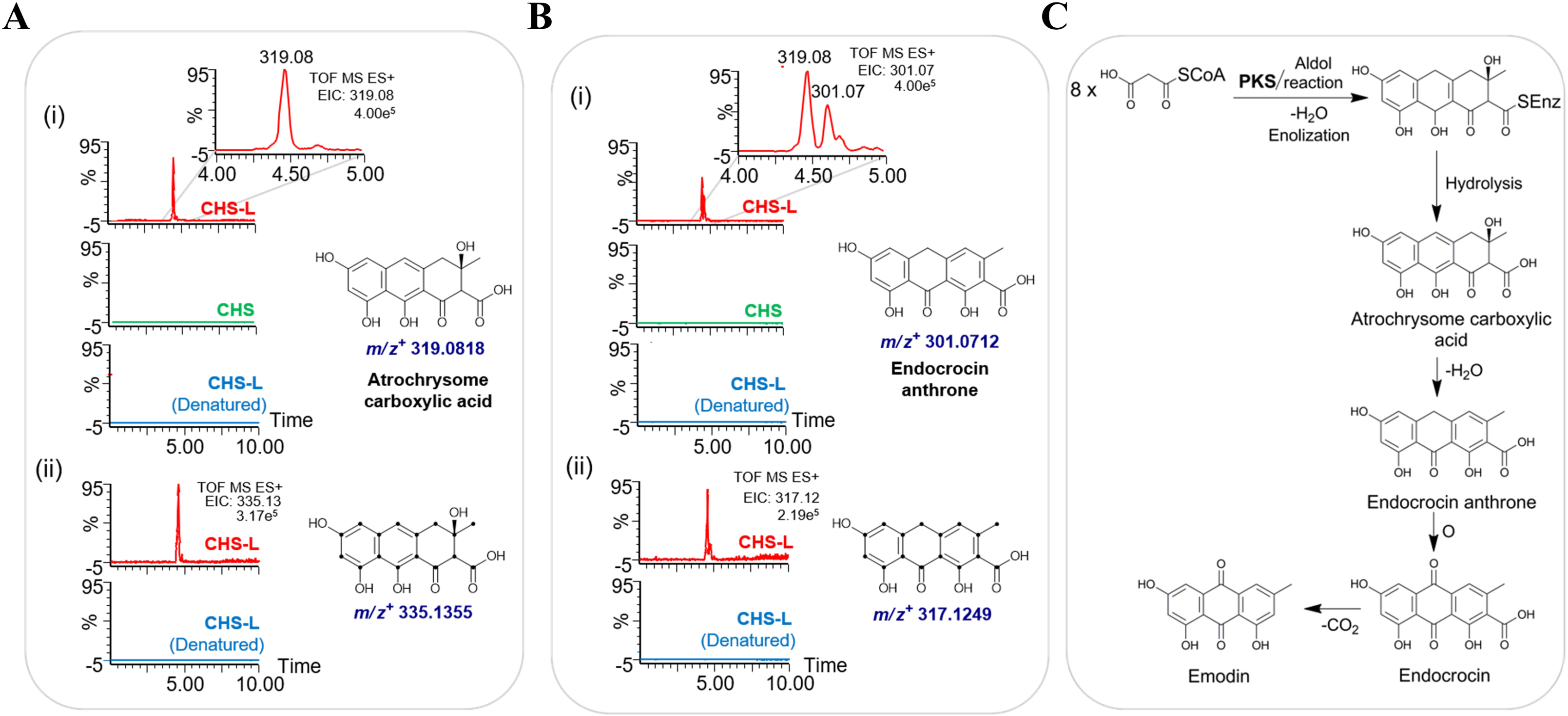
Enzyme assays and MS analysis of anthraquinones. **(A)** (i) Extracted ion chromatograms (EIC) for the compound with the mass 319.08 Da in reaction mixtures containing malonyl-CoA as substrate and chalcone synthase-like (CHS-L) (STO07G228250), chalcone synthase (CHS) (STO03G058250), or heat denatured CHS-L. (ii) EIC for the mass 335.13 Da in reaction mixtures containing ^13^C_3_-malonyl-CoA containing CHS-L or heat denatured CHS-L enzyme. Dots in the structure represent ^13^C-labelled carbons. **(B)** (i) EIC for the mass 301.07 Da in reaction mixtures containing malonyl-CoA as substrate containing CHS-L, CHS, or heat denatured CHS-L. Inset shows a zoomed region of the EIC chromatogram. (ii) EIC for the mass 317.12 Da in reaction mixtures containing ^13^C_3_-malonyl-CoA containing CHS-L or heat denatured CHS-L enzyme. **(C)** PKS-mediated biosynthetic pathway of anthraquinones. Two pathway intermediates, atrochrysome carboxylic acid and endocrocin anthrone, were produced in the CHS-L catalyzed reaction mixture.

Previous studies reported production of derailment products (SEK4 and SEK4b) by plant octaketide synthases such as HpPKS2 and ArOKS (**Fig. S1**) (*29, 30*). We performed TOF-ESI-MS_2_ analysis for both of the precursor ions 319 and 301 and compared them to mass fragments of emodin and aloe-emodin (**Figs. S18 and S19**), which share similar anthranoid scaffold and are known to be derived from atrochrysome carboxylic acid and endocrocin anthrone (**Fig. S1**). The ESI-MS_2_ fragments of the precursor ions 319 (**Figs. S20 and S21**) and 301 (**Figs. S22 and S23**) shared most of the fragments with the emodin and aloe-emodin. However, TOF-ESI-MS_2_ sister fragments of SEK4 and SEK4b reported previously (*29*) did not align to any of the fragments of standard anthraquinones, nor to atrochrysome carboxylic acid and endocrocin anthrone produced in the reactions. These evidences indicate that the metabolites produced in the reaction mixture are anthranoid scaffolds, not the octaketide shunt products. Altogether, these results provided conclusive evidence that STO07G228250 (CHS-L), a type III PKS, carries out the first committed step of anthraquinone biosynthesis via polyketide pathway in plants for the first time.

In summary, the reference genome of *S. tora* revealed a rapid evolution of putative polyketide synthase genes. By combining metabolomics, transcriptomics, and biochemical characterization of a candidate polyketide synthase, we discovered the first anthranoid forming enzyme in plants. With these tools in hand, elucidation of genes involved in the rest of the anthraquinone biosynthesis pathway in *S. tora* and other species will be accelerated. These resources can also be used as a platform to develop a medicinally useful cultivar of *S. tora* with a high content of bioactive molecules.

## Acknowledgments

We thank the National Institute of Agricultural Sciences (NAS) Genome Sequencing Core facility for their services. We thank DNA Link (Seoul, Korea) for their assistance to generate the PacBio data and contig assembly. We thank Phase Genomics (Seattle, USA) for their assistance in generation of the Hi-C data and superscaffold assembly. We thank Human Metabolome Technologies Inc. (Yamagata, Japan) for their assistance in analyzing the primary metabolites. We thank GnC Bio Co. for their assistance in generation of BAC libraries.

## Author contributions

S.H.K. conceived the project. S.H.K., C.M.L., J.S.S., J.T.J., S.Y.W., O.R.L., and T.J.O. contributed to sample preparation and sequencing. S.H.K., B.S.C., Y.Y, N.H.K., M.J., and T.H.L. performed the data analysis. R.P.P., P.B., T.S.K., and J.K.S. performed biochemistry experiments. C.K.K., J.S.K., and B.O.A. participated in methodology. S.Y.R. provided guidance on project directions and manuscript organization. S.H.K., R.P.P., and S.Y.R. wrote the manuscript. S.H.K., S.Y.R., and J.K.S. revised the manuscript. All authors read and approved the manuscript.

## Data and materials availability

Genome sequence reads, transcriptome sequence reads, Hi-C sequence reads, GBS sequence reads, and BAC library sequence data are deposited in GenBank under project number PRJNA605066. The *S. tora* genome is also available at http://nabic.rda.go.kr/Species/Senna_tora2.

## Supplementary Materials

Materials and Methods

Figures S1-S26

Tables S1-S22

Large Tables S7, S15, and S16

References (1-69)

## References and Notes

1. M. o. A. National Medicinal Plants Board, Government of India, Medicinal List. (2020, March 28).

2. R. K. Manoharan, J.-H. Lee, Y.-G. Kim, J. Lee, Alizarin and chrysazin inhibit biofilm and hyphal formation by *Candida albicans*. Front Cell Infect Microbiol 7, 447–447 (2017).

3. M. K. Parvez, M. S. Al-Dosari, P. Alam, M. Rehman, M. F. Alajmi, A. S. Alqahtani, The anti-hepatitis B virus therapeutic potential of anthraquinones derived from *Aloe vera*. Phytotherapy Research 33, 2960–2970 (2019).

4. V. C. Roa-Linares, Y. Miranda-Brand, V. Tangarife-Castaño, R. Ochoa, P. A. García, M. Á. Castro, L. Betancur-Galvis, A. San feliciano, anti-herpetic, anti-dengue and antineoplastic activities of simple and heterocycle-fused derivatives of terpenyl-1,4-naphthoquinone and 1,4-anthraquinone. Molecules 24, 1279 (2019).

5. Q.-W. Wang, Y. Su, J.-T. Sheng, L.-M. Gu, Y. Zhao, X.-X. Chen, C. Chen, W.-Z. Li, K.-S. Li, J.-P. Dai, Anti-influenza A virus activity of rhein through regulating oxidative stress, TLR4, Akt, MAPK, and NF-κB signal pathways. PLoS One 13, e0191793–e0191793 (2018).

6. M. R. Dhananjeyan, Y. P. Milev, M. A. Kron, M. G. Nair, Synthesis and activity of substituted anthraquinones against a human filarial parasite, *Brugia malayi*. Journal of Medicinal Chemistry 48, 2822–2830 (2005).

7. Y. Li, J.-G. Jiang, Health functions and structure-activity relationships of natural anthraquinones from plants. Food Funct 9, 6063–6080 (2018).

8. S. K. Ravi, R. B. Narasingappa, M. Prasad, M. R. Javagal, B. Vincent, *Cassia tora* prevents Aβ1-42 aggregation, inhibits acetylcholinesterase activity and protects against Aβ1-42-induced cell death and oxidative stress in human neuroblastoma cells. Pharmacological Reports 71, 1151–1159 (2019).

9. F.-R. Cheng, H.-X. Cui, J.-L. Fang, K. Yuan, Y. Guo, Ameliorative effect and mechanism of the purified anthraquinone-glycoside preparation from *Rheum palmatum* L. on type 2 diabetes mellitus. Molecules 24, 1454 (2019).

10. M. Chaubey, V. P. Kapoor, Structure of a galactomannan from the seeds of *Cassia angustifolia* Vahl. Carbohydr Res 332, 439–444 (2001).

11. V. E. Fernand, D. T. Dinh, S. J. Washington, S. O. Fakayode, J. N. Losso, R. O. van Ravenswaay, I. M. Warner, Determination of pharmacologically active compounds in root extracts of *Cassia alata* L. by use of high performance liquid chromatography. Talanta 74, 896–902 (2008).

12. V. K. Mahesh, R. Sharma, R. S. Singh, S. K. Upadhya, Anthraquinones and kaempferol from *Cassia* species section *fistula*. Journal of Natural Products 47, 733–733 (1984).

13. Y.-M. Chiang, E. Szewczyk, A. D. Davidson, R. Entwistle, N. P. Keller, C. C. C. Wang, B. R. Oakley, Characterization of the *Aspergillus nidulans* monodictyphenone gene cluster. Appl Environ Microbiol 76, 2067–2074 (2010).

14. G. Shamim, K. S. Ranjan, M. D. Pandey, R. Ramani, Biochemistry and biosynthesis of insect pigments. EJE 111, 149–164 (2014).

15. J. Duval, V. Pecher, M. Poujol, E. Lesellier, Research advances for the extraction, analysis and uses of anthraquinones: A review. Industrial Crops and Products 94, 812–833 (2016).

16. R. Javaid, U. Y. Qazi, Catalytic oxidation process for the degradation of synthetic dyes: An overview. Int J Environ Res Public Health 16, 2066 (2019).

17. A. Tkaczyk, K. Mitrowska, A. Posyniak, Synthetic organic dyes as contaminants of the aquatic environment and their implications for ecosystems: A review. Science of The Total Environment 717, 137222 (2020).

18. T. Awakawa, K. Yokota, N. Funa, F. Doi, N. Mori, H. Watanabe, S. Horinouchi, Physically discrete β-lactamase-type thioesterase catalyzes product release in atrochrysone synthesis by iterative type I polyketide synthase. Chemistry & Biology 16, 613–623 (2009).

19. A. J. J. Van Den Berg, R. P. Labadie, in Methods in Plant Biochemistry, J. B. Harborne, Ed. (Academic Press, 1989).

20. E. Leistner. (Springer Berlin Heidelberg, Berlin, Heidelberg, 1985), pp. 215–224.

21. E. Leistner, M. H. Zenk, Mevalonic acid a precursor of the substituted benzenoid ring of rubiaceae-anthraquinones. Tetrahedron Letters 9, 1395–1396 (1968).

22. H. J. Bauch, E. Leistner, Aromatic metabolites in cell suspension cultures of *Galium mollugo* L. Planta Med 33, 105–123 (1978).

23. E. Leistner, Isolation, identification and biosynthesis of anthraquinones in cell suspension cultures of *Morinda citrifolia* (author’s transl). Planta medica Suppl, 214–224 (1975).

24. S. Patai, Z. Rappoport, The Chemistry of the Quinonoid Compounds. (2010), pp. 1–878.

25. A. R. Burnett, R. H. Thomson, Naturally occurring quinones. Part XV. Biogenesis of the anthraquinones in *Rubia tinctorum* L. (madder). Journal of the Chemical Society C: Organic, 2437–2441 (1968).

26. T. Furumoto, A. Hoshikuma, Biosynthetic origin of 2-geranyl-1,4-naphthoquinone and its related anthraquinone in a *Sesamum indicum* hairy root culture. Phytochemistry 72, 871–874 (2011).

27. Y.-S. Han, R. v. d. Heijden, A. W. M. Lefeber, C. Erkelens, R. Verpoorte, Biosynthesis of anthraquinones in cell cultures of *Cinchona* ‘Robusta’ proceeds via the methylerythritol 4-phosphate pathway. Phytochemistry 59, 45–55 (2002).

28. I. A. M. Abdel-Rahman, T. Beuerle, L. Ernst, A. M. Abdel-Baky, E. E.-D. K. Desoky, A. S. Ahmed, L. Beerhues, In vitro formation of the anthranoid scaffold by cell-free extracts from yeast-extract-treated *Cassia bicapsularis* cell cultures. Phytochemistry 88, 15–24 (2013).

29. K. Karppinen, J. Hokkanen, S. Mattila, P. Neubauer, A. Hohtola, Octaketide-producing type III polyketide synthase from *Hypericum perforatum* is expressed in dark glands accumulating hypericins. The FEBS journal 275, 4329–4342 (2008).

30. Y. Mizuuchi, S.-P. Shi, K. Wanibuchi, A. Kojima, H. Morita, H. Noguchi, I. Abe, Novel type III polyketide synthases from *Aloe arborescens*. The FEBS Journal 276, 2391–2401 (2009).

31. I. Abe, S. Oguro, Y. Utsumi, Y. Sano, H. Noguchi, Engineered biosynthesis of plant polyketides: Chain length control in an octaketide-producing plant type III polyketide synthase. Journal of the American Chemical Society 127, 12709–12716 (2005).

32. P. P. Pillai, A. R. Nair, Hypericin biosynthesis in *Hypericum hookerianum* Wight and Arn: investigation on biochemical pathways using metabolite inhibitors and suppression subtractive hybridization. C R Biol 337, 571–580 (2014).

33. F. A. Simão, R. M. Waterhouse, P. Ioannidis, E. V. Kriventseva, E. M. Zdobnov, BUSCO: assessing genome assembly and annotation completeness with single-copy orthologs. Bioinformatics 31, 3210–3212 (2015).

34. H. Li, R. Durbin, Fast and accurate long-read alignment with Burrows-Wheeler transform. Bioinformatics 26, 589–595 (2010).

35. A. K. Anguraj Vadivel, K. Krysiak, G. Tian, S. Dhaubhadel, Genome-wide identification and localization of chalcone synthase family in soybean (*Glycine max* [L]Merr). BMC Plant Biol 18, 325–325 (2018).

36. S. A. Pandith, N. Dhar, S. Rana, W. W. Bhat, M. Kushwaha, A. P. Gupta, M. A. Shah, R. Vishwakarma, S. K. Lattoo, Functional promiscuity of two divergent paralogs of type III plant polyketide synthases. Plant Physiol 171, 2599–2619 (2016).

37. J. Andersen-Ranberg, K. T. Kongstad, M. Nafisi, D. Staerk, F. T. Okkels, U. H. Mortensen, B. Lindberg Møller, R. J. N. Frandsen, R. Kannangara, Synthesis of C-glucosylated octaketide anthraquinones in *Nicotiana benthamiana* by using a multispecies-based biosynthetic pathway. Chembiochem 18, 1893–1897 (2017).

38. W. Wang, S. Li, Z. Li, J. Zhang, K. Fan, G. Tan, G. Ai, S. M. Lam, G. Shui, Z. Yang, H. Lu, P. Jin, Y. Li, X. Chen, X. Xia, X. Liu, H. K. Dannelly, C. Yang, Y. Yang, S. Zhang, G. Alterovitz, W. Xiang, L. Zhang, Harnessing the intracellular triacylglycerols for titer improvement of polyketides in *Streptomyces*. Nat. Biotechnol. 38, 76–83 (2020).

