## Supplementary Material for "Genome-Enabled Discovery of Anthraquinone Biosynthesis in *Senna tora*"

### Supplementary Materials for
#### Genome-Enabled Discovery of Anthraquinone Biosynthesis in *Senna tora*

Sang-Ho Kang*†, Ramesh Prasad Pandey†, Chang-Muk Lee, Joon-Soo Sim, Jin-Tae Jeong, Beom-Soon Choi, Myunghee Jung, So Youn Won, Tae-Jin Oh, Yeisoo Yu, Nam-Hoon Kim, Ok Ran Lee, Tae-Ho Lee, Puspalata Bashyal, Tae-Su Kim, Chang-Kug Kim, Jung Sun Kim, Byoung Ohg Ahn, Seung Y. Rhee*, Jae Kyung Sohng*

†These authors contributed equally to this work.

#### Contents: *Senna tora* Genome Supplementary Online Materials

Section Page
Authorship Information……………………………………………………………….…1
Supplementary Text
1: DNA Sequencing, *de novo* Assembly, and validation……………………………………1
2: Linkage Map Analysis…………………………………………………………...…3
3: Pseudochromosome Construction and Genome Annotation………………………….…4
4: Comparative Analysis and Genome Evolution……………………………………..6
5: Metabolome and Transcriptome Analyses...………………………………………..6
6: Biochemistry of Anthraquinones and Flavonoids……………………………………..8
Supplementary Figures………………………………………………………….……10
Supplementary Tables………………………………………………….…………….…38
Supplementary References……………………………………………………...……55

##### 1. DNA Sequencing, *de novo* Assembly, and Validation

**DNA sequencing** – We sequenced a cultivated diploid *Senna tora* cv. Myeongyun (voucher number: IT89788) grown in Jeonju, Korea (N: 35° 49'; E: 127° 09'). Total DNA was extracted from young fresh leaves of *S. tora* cv. Myeongyun using the modified cetyltrimethylammonium bromide (CTAB) method (*1*). DNA purity and concentration were checked by electrophoresis analysis on 1.2% agarose gel and by DropSense96 Spectrophotometer (Trinean, Belgium). A total of 34 single molecule real-time (SMRT) cells were run on the PacBio RS II system and 5 cells on the Sequel system using P6/C4 chemistry. We generated a total of 80.01 Gb of clean reads (**Table S1**).

Illumina sequencing libraries were prepared according to the Illumina protocols. Briefly, 1 μg of genomic DNA was fragmented by Covaris. The fragmented DNA was repaired, and the base adenine was ligated to the 3' end. Illumina adapters were then ligated to the fragments, and the proper samples were selected. The size selected product was PCR amplified, and the final product was validated using the Agilent Bioanalyzer. Then we sequenced 200 bp paired-end (PE) and 3 to 20 kb mate-pair (MP) libraries and 500 bp PE using the HiSeq™ 2500 and MiSeq platforms (Illumina, San Diego, USA), respectively. Finally, we generated a total of 577.93 Gb of clean reads for the 200 and 500 bp PE and 3, 5, 10 and 20 kb MP libraries (**Table S1**).

**Genome size estimation** – Total Illumina DNA sequences were subjected to pre-processing steps, which included adapter trimming, quality trimming (Q20), and contamination removal. Adapter trimming and quality trimming were conducted using Trimmomatic v0.36 (*2*), and *S.*

*tora* organellar genome contamination of each sample was removed by CLCMapper v4.2.0 (https://www.qiagenbioinformatics.com/products/clc-assembly-cell/) using the chloroplast genome (Genbank ID: NC_030193) and mitochondria genome (Genbank ID: NC_038053 (*3*) sequences. All pre-processed sequences were subjected to genome size estimation using the Kmer-based method (*4*). The Kmer frequencies (Kmer size = 21) obtained using the Jellyfish v2.0 method (*5*), and the genome size was calculated by using the following formulas: 1) Genome Coverage Depth = (Kmer Coverage Depth X Average Read Length) / (Average Read Length – Kmer size +1); and 2) Genome size = Total Base Number / Genome Coverage Depth. A total of 27.5 Gb of clean Illumina reads from the 200 bp PE library were used to determine the genome size of *S. tora*. In this study, the distribution of 21 *k*-mer showed a major peak at 50x. According to the total number of *k*-mers and the corresponding *k*-mer depth, the *S. tora* genome size was estimated to be ~ 547.02 Mb (**Fig. S2**).

**Genome assembly** – High-quality PE and MP sequences (Phred score > 20) were obtained by removing low-quality sequences and duplicated reads from whole genome NGS data. Three *de novo* assemblers, SOAPdenovo v2.04 (*6*), Allpaths-LG v48777 (*7, 8*) and Platanus v1.2.1 (*9*), were performed using default parameters. For scaffolding of contig sequences, mate-pair (MP) reads were mapped to contig sequences and scaffold sequences were generated using SSPACE v3.0 with default parameters. To validate scaffold sequences, MP reads were re-mapped to the scaffold sequences and mis-scaffold sequences were disassembled into initial contig sequences using in-house script.

The average coverage of SMRT sequences was about 146x by using RS II and Sequel systems. An average subread length was about 9 kb and the maximum length was 104.5 kb. We removed the sequences of *S. tora* organellar genomes. Then, the filtered subread sequences were assembled *de novo* using the diploid assembly FALCON assembler (*10*). To increase the assembly accuracy, the length cut-off option was specified based on the subreads' N50 value of 14 kb and contigs were further corrected by Arrow (https://github.com/PacificBiosciences/GenomicConsensus, v2.1.0). To improve the quality of genome assembly results, we also performed error correction using BWA and GATK (*11*) with haplotig-merged primary contigs and Illumina reads.

To obtain the best possible draft sequence, we compared the results obtained by SOAPdenovo2, Allpaths-LG, Platanus, and FALCON algorithms. *De novo* assembly by Platanus and FALCON outperformed the results produced by SOAPdenovo2 and Allpaths-LG (**Table S2**). The number of contigs was lower, N50 length was longer, the assembled size was close to the estimated genome size, and their contiguity statistics were higher. With the assembly obtained by Platanus and FALCON, we also assessed the quality of genome assembly using Benchmarking Universal Single Copy Orthologs (BUSCO) (*12*) The percentage of complete proteins was 90.6% for the Platanus assembly and 94.3% for the FALCON assembly (**Table S3**). Based on these criteria, the assembly developed using FALCON assembler was chosen for the genome annotation. We evaluated the quality of the assembly by mapping the Illumina reads back to the scaffolds (99.7%) and expressed sequence tag (EST) sequences mapping to the scaffolds (97.2% of Iso-Seq and 89.9% of RNA-Seq) (**Table S17**), supporting the high quality of the *S. tora* genome assembly.

**Physical map validation with BAC libraries** – To validate the assembled genome against a physical map, we generated bacterial artificial chromosome (BAC) libraries. First, 15 g of young fresh leaves was harvested from growth-room-grown *S. tora* cv. Myeongyoun plants that have been placed in the dark for 48h to reduce carbohydrate concentration, which may cause carryover contamination and be detrimental to subsequent enzyme reactions. Fresh leaf tissues were ground to a fine powder in liquid nitrogen using a pestle and mortar. Leaf tissues were transferred immediately to an ice-cold lysis buffer and gently stirred to extract nuclei. The nuclei were embedded in agarose plugs and transferred to proteinase K buffer to obtain high molecular weight (HMW) DNA. The HMW DNA was partially digested using *Hin*dIII and *Bam*HI restriction enzymes and underwent a size selection three times in order to obtain consistently large inserts. Size-selected DNA was ligated with pSMART BAC vector and transformed in DH10B competent cell. The *Hin*dIII BAC library has an average insert size of 95 kb and titer of $1.6x10^6$. Certified BAC clones were colonized on agar medium and cultured in liquid medium supplemented with chloramphenicol.

The 10 BAC clones were completely sequenced using 454 Life Sciences GS FLX System (GS FLX) and ABI 3730xl DNA Analyzer. Analyzed sequencing data was assembled using Newbler v2.8 (https://www.ncbi.nlm.nih.gov/assembly/GCA_000507345.1/) and used to create contigs or scaffolds. To fill the gap of sequences, we used primer walking (*13*). The primer walking method has been widely used in genome project research to determines the order of contigs made of pieces and connects the remaining sequence gap between the contigs. In this way, a draft sequence for 10 BAC clones was created. Finally, completed BAC clone sequences were checked by using the HiSeq sequence data for sequence error correction. To validate the genome assembly obtained by FALCON, we performed all-by-all alignment (-minIdentity=80-99, -minScore=100, -fastMap) of the 10 complete BACs and the assemblies using BLAT v3.2.4 (*14*).

#### 2. Linkage Map Analysis

**Genotype-by-sequencing (GBS) linkage analysis** – Genomic DNA was extracted from the two parents (*S. tora* cv. Myeongyun (voucher number: IT89788) and ST-9 (voucher number: IT104602)) and 153 F2 progeny using a Qiagen plant DNAeasy kit. Two GBS libraries were prepared using *Ape*KI restriction enzyme as described in Elshire *et al.* (*15*). The GBS libraries (74 F2 individuals + two parents; 79 F2 individuals + two parents) were sequenced on an Illumina HiSeq2500 system. Low quality bases and adapter sequences were trimmed using Trimmomatic v0.36 (*2*) and the trimmed reads from each sample were mapped to the *S. tora* draft assembly using BWA-MEM (*16*). HaplotypeCaller in GATK (*11*) was used to call single nucleotide polymorphisms (SNPs) and generate a raw vcf file. High-quality biallelic SNPs were selected using VCFtools (*17*) with the following conditions: 1) minimum read depth $\geq 5$; 2) minimum genotype quality $\geq 20$; and 3) missing genotype $\leq 30\%$. The SNP positions that showed polymorphic homozygous SNPs between the parents were retained for linkage analysis. Linkage analysis was conducted using QTL IciMapping v4.1 (*18*) with the Kosambi function.

A total of 721.8 million raw PE reads was generated from two *Apek*I GBS libraries, and 372 million trimmed PE reads were used for subsequent linkage analysis. Of those, about 89.8% reads were mapped to the *S. tora* reference assembly and 88.6% (329.5 million reads) were concordantly mapped, which was representing about 2.1 million properly mapped PE reads per sample. The GATK HaplotypeCaller called 289,768 and 4.78 million unfiltered variants from library 1 and 2, respectively. After low-quality SNPs were filtered, 7,584 and 15,604 high-quality SNPs were obtained from library 1 and 2, respectively, and 5,071 markers were commonly presented in both. Three genetics maps independently constructed with three sets of SNP markers (from library 1, 2 and common) were evaluated in terms of number of anchored contigs and genome representation, and the map used common markers between library 1 and 2 was selected for further analysis. This map contained 2,654 non-redundant markers representing 3,587 cM within 12 linkage groups (LG13 contained only one marker). With this linkage map, we tried to re-group markers by increasing group number parameters from 13 to 25, however the efforts were not successful to make 13th linkage group. Finally, the linkage map was compared to pseudochromosomes constructed by Hi-C and we were able to split LG5 into two groups; one with three contigs (164 markers covering 34.4 Mb) and the other with 8 contigs (263 markers covering about 44 Mb). The final *S. tora* genetic map with Hi-C information was resulted in 13 linkage groups with 4,455 markers spanning 2,780 cM of genetic distance. It enabled to anchor 111 contigs (contig3 split into two contigs; c31-1, c31-2) to 13 linkage groups, which represented about 401 Mb of *S. tora* sequence assembly (**Table S18**). Genetically, LG8 was the longest linkage group (347.5 cM) followed by LG13 (343.8 cM) and LG5 (313 cM). Whereas, 487 markers anchored about 45 Mb of sequences in LG5, which was the longest anchored chromosome. Physical distance per genetic distance was calculated as 144 kb/cM in average across whole genome (**Fig. S7**).

##### 3. Pseudochromosome Construction and Genome Annotation

**Hi-C library construction** – To generate pseudochromosomes, chromatin conformation capture (Hi-C) data was generated using a Phase Genomics (Seattle, WA) Proximo Hi-C Plant Kit, which is a commercially available version of the Hi-C protocol (*19*). Intact cells from two samples were crosslinked using a formaldehyde solution, digested using the *Sau*3AI restriction enzyme, and proximity ligated with biotinylated nucleotides to create chimeric molecules composed of fragments from different regions of the genome that were physically proximal *in vivo*, but not necessarily genomically proximal. Continuing with the manufacturer's protocol, molecules were pulled down with streptavidin beads and processed into an Illumina-compatible sequencing library. Sequencing was performed on an Illumina NextSeq 500 (Illumina, San Diego, USA), generating a total of 188,501,285 PE read pairs.

**Pseudochromosome construction by Hi-C** – Reads were aligned to the reference assembly following the manufacturer's recommendations (https://phasegenomics.hithub.io/2019/09/19/hic-alignment-and-qc.html). Briefly, reads were aligned using BWA-MEM with the -5SP and -t 8 options specified, and all other options as default. SAMBLASTER (*20*) was used to flag PCR duplicates, which were later excluded from further analysis. Alignments were then filtered with samtools (*21*) using the -F 2304 filtering flag to remove non-primary and secondary alignments.

Phase Genomics' Proximo Hi-C genome scaffolding platform was used to create chromosome-scale scaffolds from the corrected assembly (*22*). Similar to the LACHESIS method (*23*), this process computes a contact frequency matrix from the aligned Hi-C read pairs, normalized by the number of *Sau*3AI restriction sites (GATC) on each contig, and constructs scaffolds in such a way as to optimize expected contact frequency and other statistical patterns in Hi-C data. Approximately 20,000 separate Proximo runs were performed to optimize the number of scaffolds and scaffold construction in order to make the scaffolds as concordant with the observed Hi-C data as possible. The Hi-C sequences were aligned to the draft contig assemblies. Finally, Juicebox (*24*, *25*) was used to correct scaffolding errors as well as to introduce two new breaks into two putative mis-joined contigs (contigs 3 and 110). All contig sequences not anchored to chromosomes were constructed with 100 N's as linker following the order of contig sizes designated chromosome 00 (Chr 00). The length and number of contigs for each chromosome are shown in **Table S19**.

**Annotation of repetitive DNA** – Initially, repeat regions were predicted using the *de novo* method and classified into repeat subclasses. *De novo* repeat prediction for *S. tora* was conducted using RepeatModeler (http://www.repeatmasker.org/RepeatModeler/), which includes other methods such as RECON (*26*), RepeatScout (*27*), and TRF (*28*). Furthermore, the repeats were masked using RepeatMasker v4.0.5 (http://www.repeatmasker.org/) with

RMBlastn v2.2.27+ and classified into its subclasses with the reference of Repbase (*29*) v20.08 databases (https://www.girinst.org/repbase/).

Transposable elements are major components of plant genomes, but they have not been examined in *S. tora*. The *S. tora* genome masked 53.9% of the assembly as repeat sequences. Long terminal repeat (LTR) retrotransposons, mainly Gypsy-type LTRs, are the most abundant, occupying 15.6% of the genome (**Table S20**). The fraction of repeat sequences in the genome is very similar with other Leguminosae family plants such as pigeon pea (51.6%) (*30*), mung bean (50.1%) (*31*), and chickpea (49.4%) (*32*).

**Genome annotation** – The genes from the *S. tora* reference genome were predicted using an in-house gene prediction pipeline, which includes three modules: evidence-based gene modeler, *ab-initio* gene modeler, and consensus gene modeler. To improve the accuracy of gene prediction, we downloaded a total of 118,390 Iso-Seq reads in GenBank SRA database (SRP159435) (*33*). RNA-Seq from five tissues (leaf, root, stem, flower, and dry seed) and Iso-Seq data were aligned against the *S. tora* genome. The detail of the pipeline was described previously (*34, 35*). Initially, the sequenced transcriptomes were mapped to the *S. tora* repeat-masked reference genome using Tophat (*36*), and transcripts/gene structural boundaries were predicted using Cufflink (*36*) and PASA (*37*). To train the *ab-initio* gene modeler AUGUSTUS (*38*) and evidence-based gene modeler GENEID (*39*), we selected a few of genomes using Exonerate (*40*). Genomes we used are: *Abrus precatorius, Arachis hypogaea, Arachis duranensis, Arachis ipaensis, Cajanus cajan, Cicer arietinum, Faidherbia albida, Glycine max, Glycine soja, Lablab purpureus, Lupinus angustifolius, Medicago truncatula, Mucuna pruriens, Phaseolus vulgaris, Prosopis alba, Sclerocarya birrea, Trifolium medium, Trifolium subterraneum, Vigna angularis, Vigna radiata, Vigna subterranea, Vigna unguiculata, Arabidopsis thaliana*. Finally, the predicted *ab-initio* gene models, transcript models, and evidence-based gene models were subjected to build consensus gene models. The consensus genes were subjected to functional annotations from biological databases (NCBI - NR databases, Swiss-Prot, gene ontologies and KEGG pathways) by using the Blast2GO (*41*). The transcription factor genes were predicted through searching DNA binding domains using InterProScan v5.36-75.0 (*42*) and the family name assigned through the rules given in PlantTFDB (v5.0, http://planttfdb.cbi.pku.edu.cn/). The gene models were supported by 97.2% Iso-Seq data, which was comprised of 118,390 high-quality isoforms derived from leaf, root, and two different developmental stages of seeds, and 89.9% RNA-Seq data derived from seed, leaf, root, stem, flower, and seven different stages of seeds**,** suggesting that the assembly includes most of the *S. tora* gene space (**Table S17**).

**Identification of long non-coding RNA (lncRNA)** – A pipeline for lncRNA identification was designed according to a previous study (*43*). In brief, among total transcripts obtained from reference-guided assembly of transcriptome data, transcripts with open reading frame (ORF) for ≥ 100 amino acids and ≤ 200 nucleotides were removed. We also removed sequences with homology to protein sequences based on BLAST search against the SwissProt (*44*) and Pfam (*45*) protein databases. The coding potential of remaining sequences was calculated using Coding Potential Calculator (CPC) (*46*) and transcripts with CPC score ≥-1.0 were removed as authors of (*46*) suggest that CPC scores between -1 and 1 are 'weak noncoding' or 'weak coding'. From the remaining transcripts, housekeeping RNAs (tRNA, rRNA, snRNA, snoRNA and etc.) were removed by comparing with RNACentral database (*47*) sequences (cutoff E-value of 1e-10) and those completely matching with *S. tora* reference protein coding gene sequences were also removed. Finally, we only retained the longest isoform for each gene to obtain the final set of 3,278 lncRNAs (**Table S7**).

#### 4. Comparative Analysis and Genome Evolution

**Phylogenetic tree construction and evolution rate estimation** – To understand the evolutionary patterns of the *S. tora* genome and gene families, we performed comparative genome analysis. We used 14 legume species and one outlier (*Vitis vinifera*). The OrthoMCL v2.0.9 (*48*) method was used to find orthologus groups in the given genomes. The orthologus clusters that contain protiens from all 16 speceis were subjected to multiple sequence alignemnt with MAFFT v7.305b (*49*) and the alignments were corrected with Gblocks v0.91b (*50*). The phlylogenetic tree was re-constructed using IQ-Tree v1.5.0-beta (*51*), using a maximum likelihood method with 1,000 bootstrap iterations. Here, the longest protein in each genome was selected among the proteins in each orthologous cluster. From the trees, the gene pattern changes such as contraction and expansion were observed among the genomes using CAFE v3.1 method (*52*). Rapid expansion/contraction is indicated by statistically significant and non-random expansion/contraction at $p < 0.01$ as described in CAFE (*52*). The evolutionary divergence time scale of the species was obtained from the clock and Yule model with JTT substitution model (the gamma category count set to 4), which was implemented in BEAST2 method (*53*). The calibration priors were set as 58-70 MYA for the common ancestor of *S. tora, C. fasciculata, M. pudica, and M. truncatula* and 105-115 MYA for the root according to the TimeTree database (http://timetree.org).

**Ks analysis** – To calculate the synonymous substitution Ks values, we selected the orthologous gene pairs between species and the paralogous pairs within a species from the orthology analysis. The selected proteins were further subjected to multiple sequence alignment with MAFFT v7.305b (*49*) and corrected with Gblocks v0.91b (*50*). The corresponding genomic regions of conserved proteins, which were observed from the corrected multiple alignments, were subjected to Ks calculation using ParaAT v2.0 (*54*) with the Yang-Nielsen approach implemented in PAML (*55*). The Ks distribution plot (**Fig. S24**) was drawn using in-house Python and R scripts.

Ancient whole-genome duplication (WGD), also known as paleopolyploidization events, is shared throughout angiosperm history (*56*) and represents a powerful evolutionary force for diversification, neo-functionalization, and innovation (*57-59*). We did not detect the peak of the recent WGD found in soybean, suggesting that Caesalpinoideae including *S. tora* and *Mimosa pudica* do not have the soybean-specific WGD event (**Fig. S24**) (*60*). Homology analysis with 6,310 orthologous genes shared by *S. tora* and 15 other green plant species was used to construct a phylogenetic tree based on a concatenated sequence alignment using MAFFT v7.305b. In this phylogenetic tree, *S. tora*, as expected, clustered with other legume crops, although the evolutionary distance from *S. tora* to Papilionoideae such as soybean, *Medicago truncatula*, and chickpea was relatively large (**Fig. S11**). The phylogenetic tree confirmed the grouping of Caesalpinioideae species such as *S. tora* and *M. pudica*. The first divergence time between Caesalpinioideae and Papilionoideae was estimated at approximately 81.9-93.6 MYA (**Fig. S11**). Furthermore, *Senna* and *Chamaecrista* genera diverged from the Mimosoid clade (*Faidherbia albida* and *Mimosa pudica*) about 59.4-66.5 MYA (**Fig. S11**) (*61*).

#### 5. Metabolome and Transcriptome Analyses

**Primary metabolites profiling** – Metabolome analysis was performed with 21 samples of frozen seed powders (~50 mg each) collected from 7 seed developmental stages using Capillary Electrophoresis Time of Flight Mass Spectrometry (CE-TOF-MS). CE-TOF-MS was run in two modes for cationic and anionic metabolites at Human Metabolome Technologies (Yamagata, Japan). The samples were mixed with 500 μL of methanol containing internal standards (50 μM) and homogenized using a homogenizer (a cell breakage machine with beads (MS-100R, TOMY Digital Biology, Tokyo, Japan)). Then, chloroform (500 μL) and Milli-Q water (200 μL) were added to the homogenates, mixed thoroughly and centrifuged (2,300 x *g*, 4°C, 5min). The water layer (200 μL) was filtrated twice through 5-kDa cut-off filter (Ultra-free MC-PLHCC, Human Metabolome Technologies, Yamagata, Japan) to remove macromolecules. The filtrate was centrifuged, and re-suspended in 50 μL of ultra-pure water immediately before the measurement. Cationic metabolite levels were analyzed using a commercial fused silica capillary (H3305-1002, HMT; i.d. 50 μm x 80 cm) with a commercial cationic electrophoresis buffer (H3301-1001, HMT), or anionic electrophoresis buffer (H3301-1020, HMT) as the electrolyte. A commercial sheath liquid (H3301-1020, HMT) was delivered at a rate of 10 μl/min. Approximately 10 nL of sample solution was injected at a pressure of 50 mbar for 10 sec, and applied capillary voltages was set at 27 kV (cation mode) and 30 kV (anion mode), respectively. For both cationic and anionic modes, the spectrometer was scanned from *m/z* 50 to 1,000. Other conditions were followed as previously described (*62*).

**Data processing of primary metabolites** – Peaks detected in CE-TOF-MS were extracted using an automated integration software (MasterHands ver. 2.16.0.15 developed at Keio University) in order to obtain peak information including *m/z*, migration time (MT), and peak area. The peak detection limit was set at the signal-noise ratio (S/N) of 3. Signal peaks corresponding to isotopomers, adduct ions, and other product ions of known metabolites were excluded, and remaining peaks were annotated with putative metabolites from the MasterHands database based on their MTs and *m/z* values. The tolerance range for the peak annotation was configured at ±0.5 min for MT and ±10 ppm for *m/z*. For the 178 peaks detected (**Table S15)**, average relative area and standard deviations (S.D.) were calculated in the 7 developmental stages of *S. tora* seeds. Absolute quantification was performed for 110 metabolites including glycolytic and TCA cycle intermediates, amino acids, and nucleic acids. All the metabolite concentrations were calculated by normalizing the peak area of each metabolite with respect to the area of the internal standard and by using standard curves, which were obtained by single-point (100 μM) calibrations. Finally, we obtained absolute quantitative values for 69 out of 110 metabolites **(Table S16**). The ratio of average relative peak area and *p*-value from Welch's *t*-tests were calculated between the two stages (stage 1 *vs*. other stages).

**Anthraquinone extraction and Liquid Chromatography-Mass Spectrometry (LC-MS) analysis** – *S. tora* seeds were collected and sorted into seven different ripening stages (Stage1-Stage7) depending on their size, color, and hardness. Classified seeds were ground with a mortar and pestle using liquid nitrogen to a fine powder and freeze-dried. Powdered samples (20 mg) were extracted with 1 mL of methanol using sonication for 30 min at 60 °C. After extraction, samples were centrifuged at 500 x *g* for 3 min at 25 °C and the supernatant was filtered with 0.2 μm Acrodisc® MS syringe filters with PTFE membrane (Pall Corporation, Port Washington, NY, USA). The filtrate was completely dried by EvaT-0200 Total Concentration System equipped with EvaS-3600 N2 generator (Goojung engineering, Seoul, Korea), mixed with methanol, and filtered again with Acrodisc® 0.2 μm MS syringe filter for LC-MS analysis.

Quantitative analysis of anthraquinones was performed by a 3200 QTRAP mass spectrometer with a Turbo V ion source (AB Sciex, Ontario, CA, USA) coupled with a VANQUISH UHPLC system (ThermoFisher Scientific, CA, USA) equipped with binary solvent manager, sample manager, column heater, and photodiode array detector. UHPLC was performed on a ZORBAX Eclipse Plus column (1.8 μm, 2.1 mm x 100 mm, Agilent Technology, CA, USA) and mobile phases consisted of 5 mM ammonium acetate in water (eluent A) and 100% acetonitrile (eluent B). The gradient conditions were as follows: 0-1 min, 10% B; 1-4.5 min, 10-30% B; 4.5-8 min, 30-50% B; 8-11 min, 50-100% B; and 11-14 min, 100% B. The flow rate was 0.5 mL/min and two microliters of samples were injected. For detecting peaks from test samples, MS parameter in ESI-negative mode was used as follows: nebulizing gas, 50 psi; heating gas, 50 psi; curtain gas, 20 psi; desolvation temperature, 550 °C; and ion spray voltage floating, 4.5 kV. The data obtained from MRM mode was quantitated using MultiQuant 3.0.2 software (AB SCIEX).

**RNA sequencing and analysis** – Total RNA was isolated from seven developmental stages of seeds (Stage1-Stage7) (**Table S21**). RNA extraction and RNA-Seq library preparations were performed as described previously (*31*), and RNA-Seq libraries were sequenced on the Illumina NextSeq 500 (Illumina, San Diego, USA). First, low-quality bases (PHERD score (Q) < 20) and adaptor contamination were removed by Trimmomatic v0.36 using the parameters 'ILLUMINACLIP:TruSeq3-SE:2:30:10 LEADING:3 SLIDINGWINDOW:4:15 MINLEN:36' (*2*). After checking for quality scores and read lengths, RNA-Seq reads were mapped *to S. tora* genome using STAR-2.6.0a with default parameters (*63*). Expectation Maximization (RSEM-1.3.1) (*64*) method was used to obtain the expression value for each gene in the genome. The read counts estimated by RSEM were subjected to edgeR v3.22.5 (*65*) to obtain differential expression scores along with the statistical significance based on false discovery rate (FDR). Furthermore, we applied the standard filters, *i.e.*, genes per million (TPM) ≥ 0.3, read counts ≥ 5 and log2 fold changes ≥ 1 or ≤ -1 to derive the final list of differentially expressed genes (*66*). Finally, the expressed genes, (*i.e.*, TPM ≥ 0.3 and read count ≥ 5) were included to show the different expression patterns during seed development. An in-house R script was used to generate the heatmap.

#### 6. Biochemistry of Anthraquinones and Flavonoids

**Heterologous protein expression and enzyme assays** – STO07G228250 (1173 bp) encoding CHS-L and STO03G058250 (1173 bp) encoding CHS cDNAs were PCR amplified using a pair of oligonucleotide primers (**Table S22**). STO07G228250 and STO03G058250 were cloned in pET28a(+) vector. The *E. coli* BL21 (DE3) strain harboring correct pET28a(+)_STO07G228250 and pET28a(+)_STO03G058250 plasmids were used for protein production. Cultures were induced by 0.4 mM isopropyl-β-D-thiogalactopyranoside (GeneChem, Daejeon, Korea) to start the recombinant protein expression. After incubation at 20 °C for 20-24 h, the cells were harvested by centrifugation, washed twice with 100 mM phosphate buffer saline (pH 7.5) containing 10% glycerol and disrupted by sonication. The homogenates were centrifuged at 12,000 rpm (13475 x *g*) for 30 min at 4 °C to isolate soluble proteins from insoluble cell debris. The supernatants were applied to a separate column containing 1 ml of $His_6$ Ni-Superflow Resin (Takara, Japan) which was equilibrated with a buffer containing 100 mM phosphate buffer saline (pH 7.5), 500 mM NaCl, 5 mM imidazole, 1 mM dithiothreitol, and 10% glycerol. The $His_6$-tagged recombinant proteins were then eluted with eight column volumes of the aforementioned buffer containing 50 mM of imidazole. The elution was repeated with the same buffer containing 250 mM of imidazole. Purity and molecular mass of the recombinant proteins were verified by 12% SDS-PAGE. The fractions containing the pure protein were then pooled and concentrated using Amicon Ultra 15 (Millipore, 30 K NMWL centrifugal filters). Protein concentrations were measured by the Bradford method using the Bradford reagent (Protein Assay Dc, Bio-Rad, Hercules, CA, USA) using bovine serum albumin as standard.

Enzyme assays for anthraquinone biosynthesis were carried out in 1 ml volume in a microcentrifuge tube containing 5 mM of malonyl-CoA (Sigma-Aldrich, St. Louis, USA), 10 mM of $MgCl_2$, and 10 µg/ml of pure protein in 100 mM phosphate buffer saline (pH 7.5). An identical reaction mixture containing the same amount of heat-denatured protein served as a negative control. A separate reaction was carried out with the same reaction constituents with additional 1 mM of NADPH as a co-factor. Similarly, assays were carried out with identical reaction components except for malonyl-CoA, which was replaced with the same amount of $^{13}C_3$-malonyl-CoA (Sigma-Aldrich, St. Louis, USA). All reaction mixtures were incubated at 30 °C for 6 h, and stopped by heating the reaction mixture at 85 °C for 3 min.

For STO03G058250 (CHS) enzyme, separate sets of reactions were carried out in the presence of *p*-coumaroyl-CoA (PlantMetaChem, Giessen, Germany) as the starting substrate and malonyl-CoA and $^{13}C_3$-malonyl-CoA as extender substrates. Each reaction mixture contained 2 mM of *p*-coumaroyl-CoA, 5 mM of extender substrates, 10 mM of $MgCl_2$, and 10 µg/ml of pure protein in 100 mM phosphate buffer saline (pH 7.5). An identical reaction mixture without the starting substrate served as a negative control. Each reaction was performed in three biological replicates.

**Enzyme assay quantification** – The quenched reaction mixtures were centrifuged at 13,475 x *g* for 30 min to separate denatured protein, filtered through 0.2 µm syringe filter, and subjected to reverse phase ultra-pressure liquid chromatography (RPUPLC) coupled with photo-diode array (PDA) when necessary, followed by high-resolution time-of-flight electrospray ionization (HRTOF ESI-MS) analysis. All enzyme assays were performed in triplicates.

RPUPLC-PDA was performed with an RP-18 column (50 mm long, 2.1 mm internal diameter, 1.7 µm particle size) in Acquity (Waters) with UPLC LG 500nm PDA detector using water as aqueous solvent A and acetonitrile (Thermo Fisher Scientific Korea, Seoul, Korea) as organic solvent B at the flow rate of 0.3 mL/min for 12 min under the following conditions of solvent B (0-100%) for (0-7) min, 100% for (7-9.5) min, and 0% for (9.6-12) min. HRQTOF ESI-MS and ESI-$MS_2$ were performed in Acquity SYNAPT G2-S mass spectrometer (Waters, Milford, MA, USA). The selected precursor ions were further subjected to TOF ESI-$MS_2$ analysis in positive ionization mode.

The CHS enzyme STO03G058250 was investigated for its possible involvement in flavonoids biosynthesis. The reaction of STO03G058250 with *p*-coumaroyl-CoA and malonyl-CoA generated naringenin chalcone along with bisnoryangonin, and *p*-coumaroyltriacetic acid lactone (CTAL) demonstrating its participation in flavonoids biosynthesis in *S. tora* (**Figs. S25 and S26**). The pyrone ring containing metabolites, bisnoryangonin and CTAL, are the shunt products produced after two and three malonyl-CoAs condensation, respectively (*67*). None of the stilbene type derivatives were produced in the reactions.

#### Supplementary Figures:

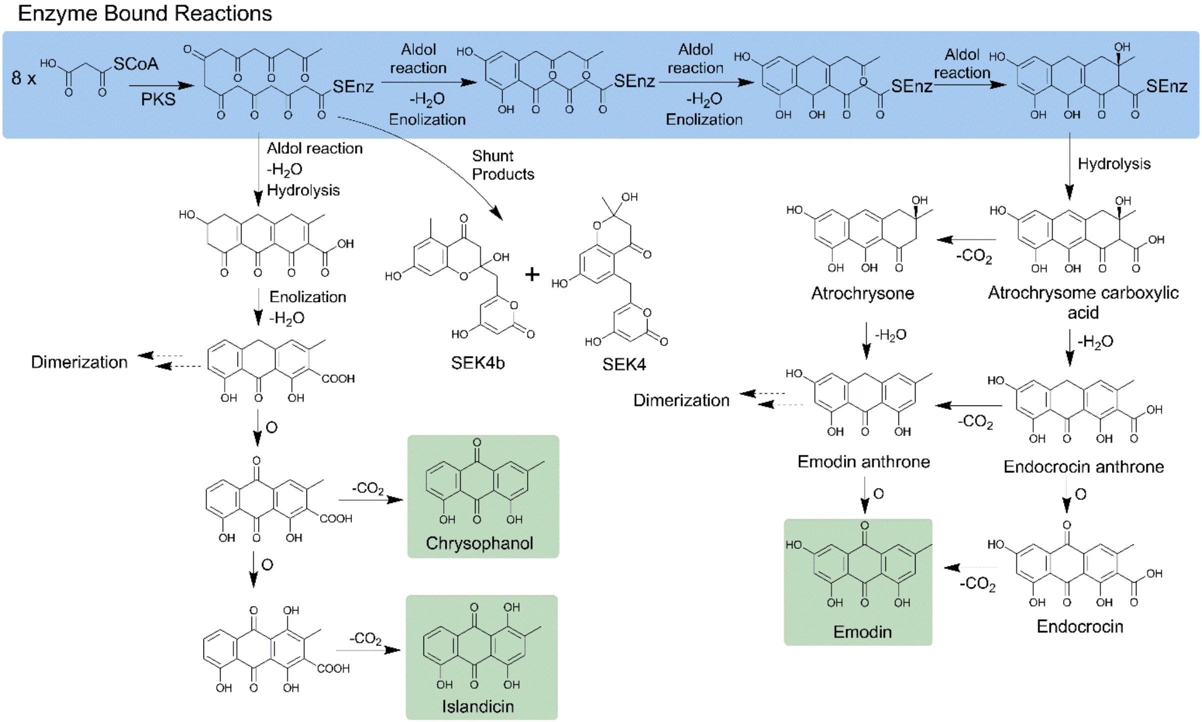


**Fig. S1.** Proposed PKS-mediated anthraquinone biosynthesis pathways. Eight molecules of malonyl-CoA are condensed to produce a linear octaketide non-reduced polyketide, which undergoes sequential cyclization and enolization (highlighted in blue shade), and released from PKS to produce atrochrysome carboxylic acid, the first PKS-produced anthranoid scaffold. Decarboxylation and oxidation results in final anthraquinones such as emodin. The intermediates might undergo dimerization reactions to produce anthraquinone dimers. Biosynthesis of anthraquinones such as chrysophanol and islandicin follows different dehydration and enolization steps. Final anthraquinones highlighted in green shade.

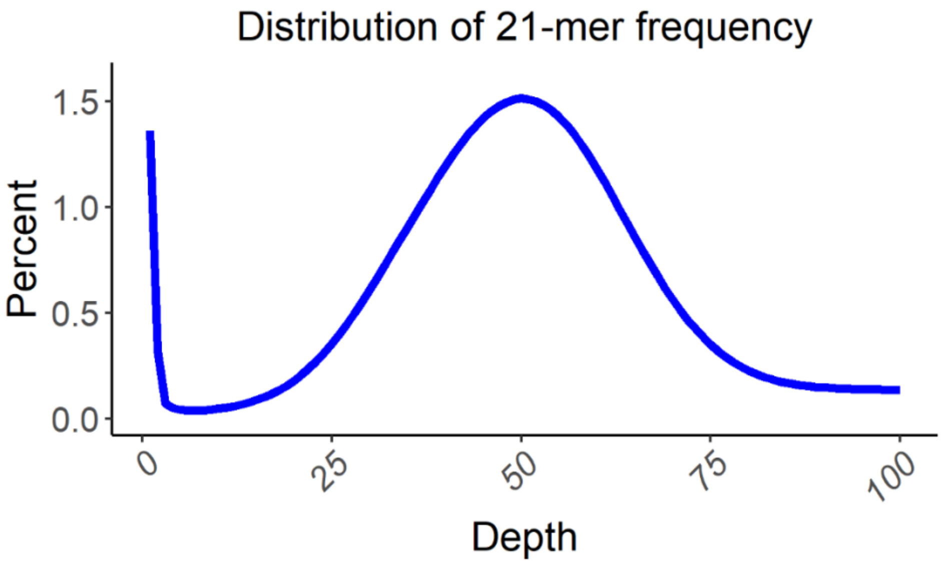


**Fig. S2.** Genome size estimation from distribution of 21-mer frequency in the sequencing reads. The reads used for *k*-mer distribution analysis were from the 200 bp paired-end library. A total of 27.5 Gb high-quality short-reads were used and only one peak was observed.

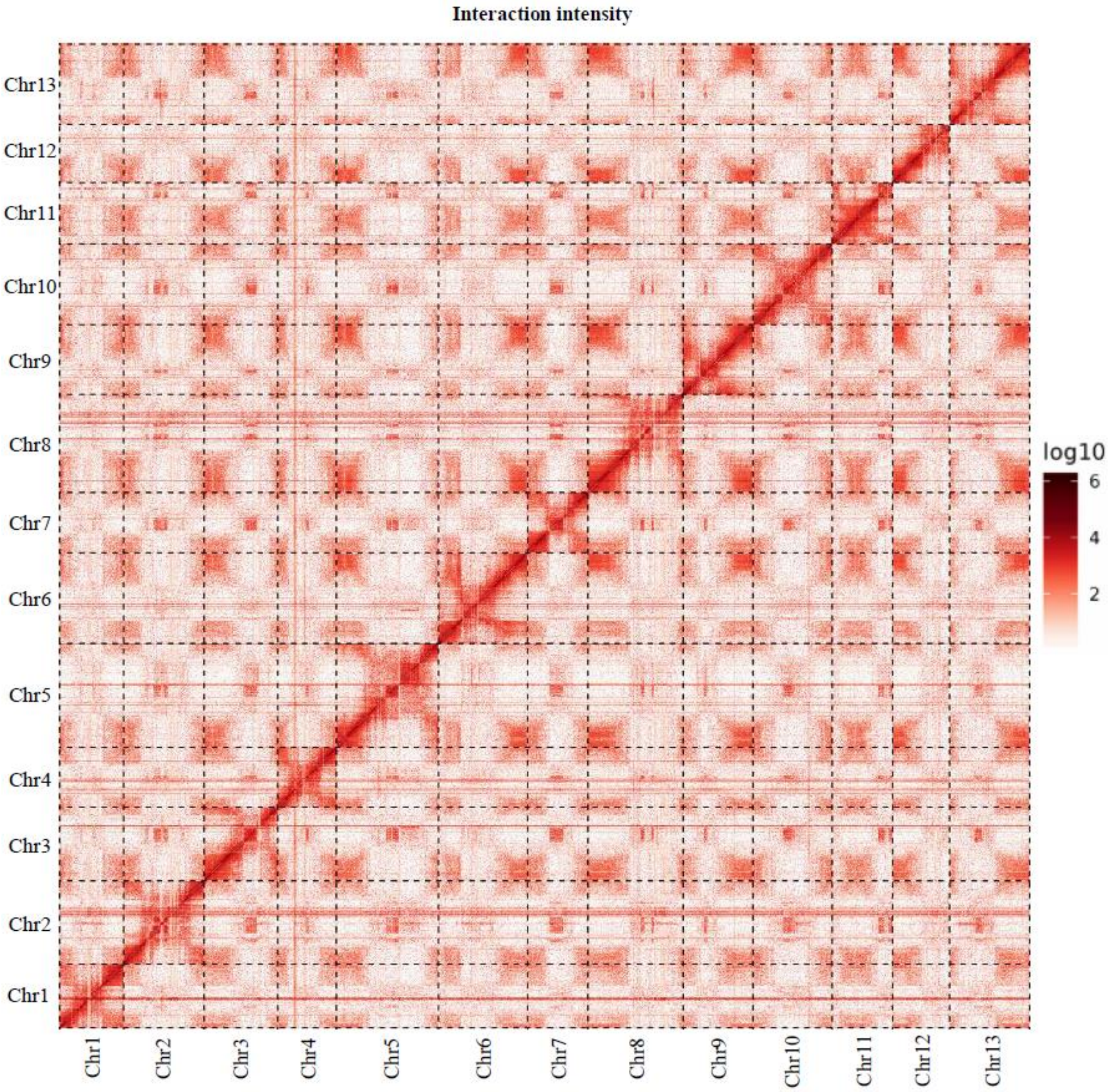


**Fig. S3.** Heatmap of chromosome conformation capture analysis. The intensity of interaction indicates the normalized count of Hi-C link on a logarithmic scale. The colored bar on the right represents the strength of interaction.

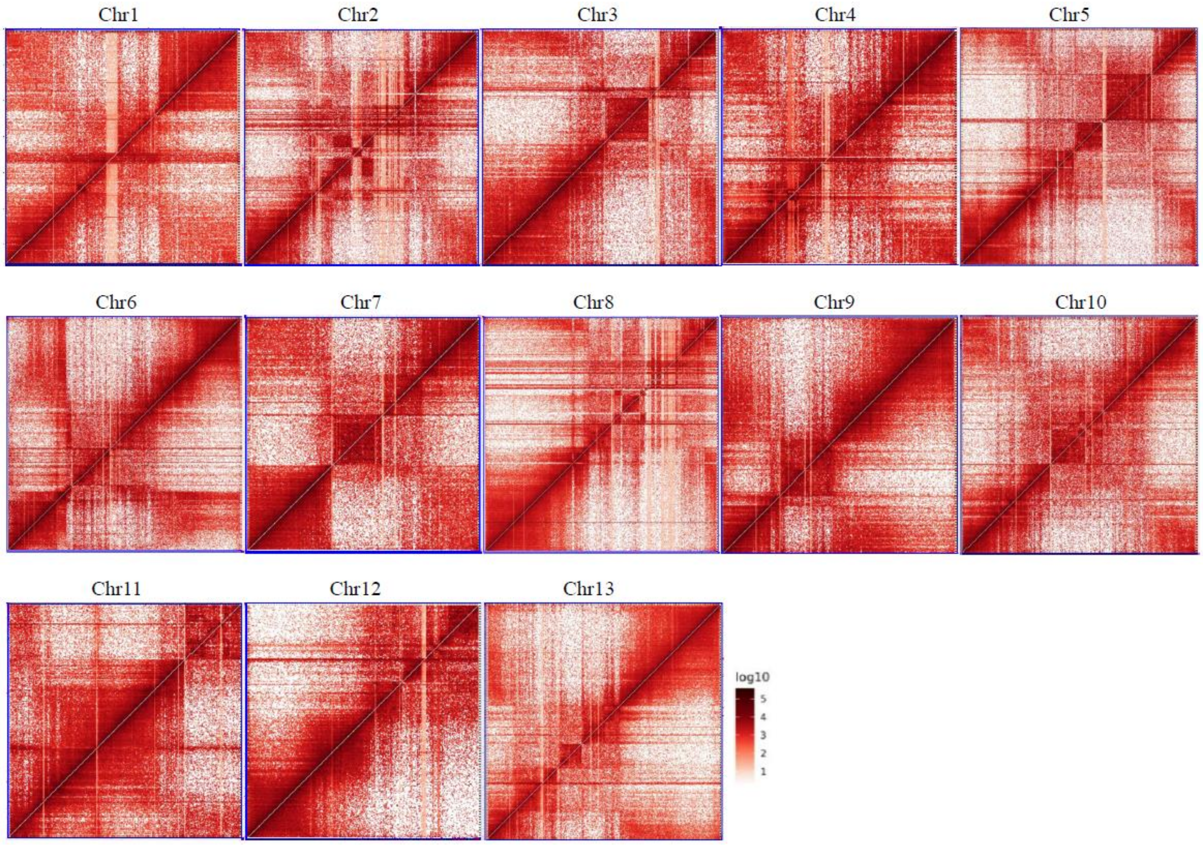


**Fig. S4.** Genome-wide analysis of chromatin interactions at 100-kb resolution in *S. tora* genome. The colored bar represents the strength of interaction.

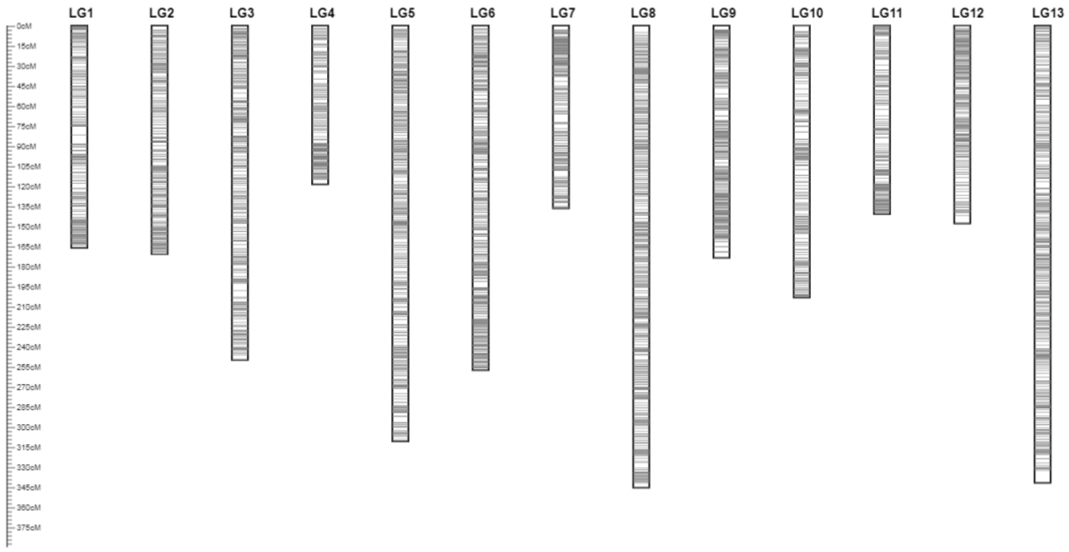


**Fig. S5.** Linkage map of an *S. tora* $F_2$ population (Myeongyun x ST-9). Linkage map was generated by integrating maps from two independent $F_2$ libraries. Gray bands in each linkage group indicate mapped markers. Numbers of each linkage group correspond to the numbering of chromosomes in this work.

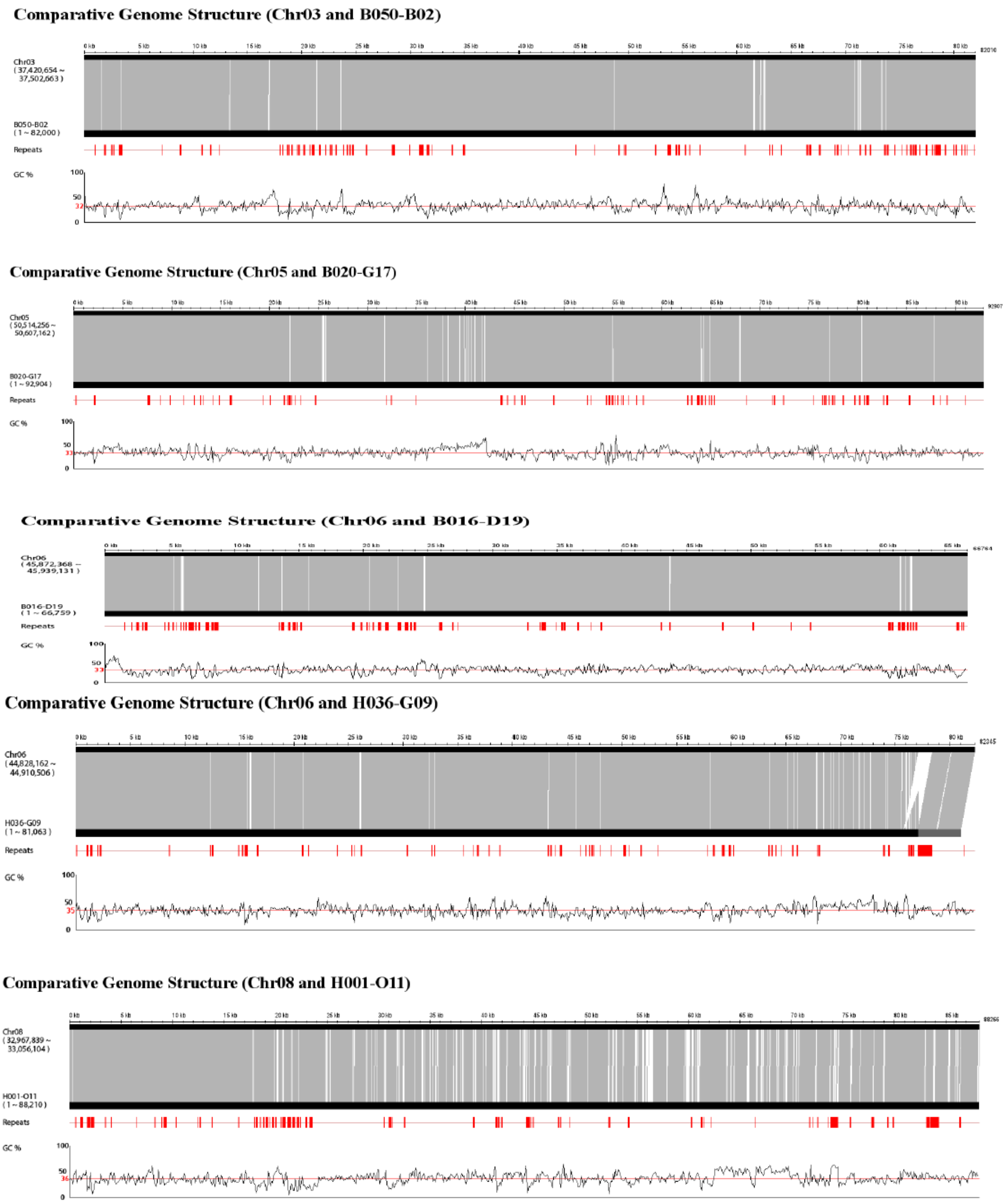


**Fig. S6.** Genome coverage evaluated by ten fully sequenced BAC clones sequenced by Sanger technology and 454 Life Sciences GS FLX System. Repeats (transposable elements) and GC contents (%) are also shown.

**Comparative Genome Structure (Chr08 and H019-A05)**

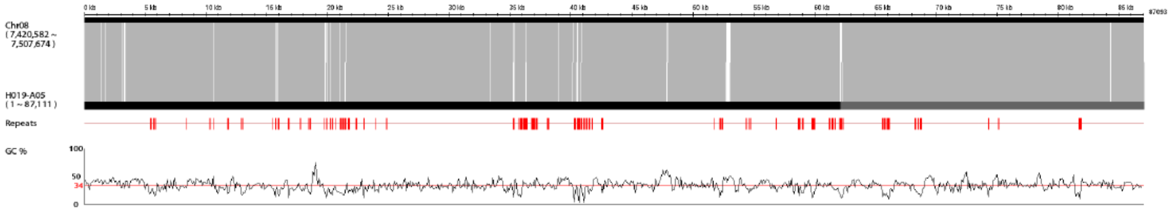


**Comparative Genome Structure (Chr08 and H024-N24)**

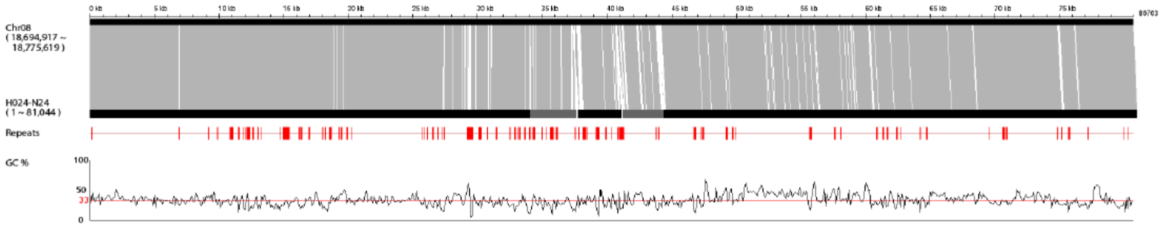


**Comparative Genome Structure (Chr09 and H002-L14)**

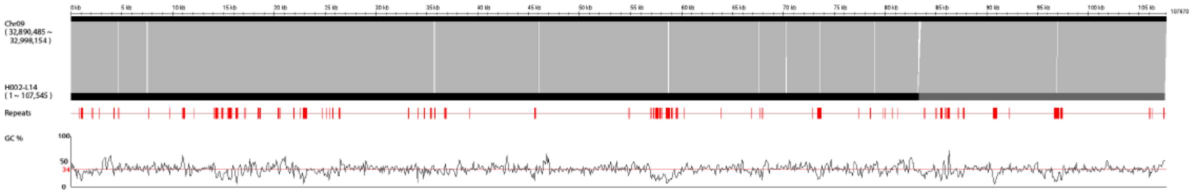


**Comparative Genome Structure (Chr10 and B011-O10)**

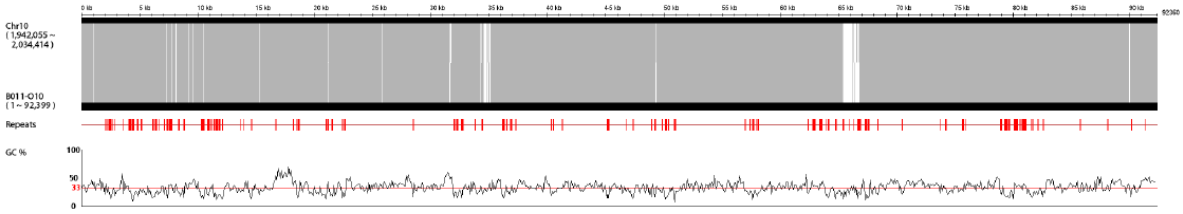


**Comparative Genome Structure (Chr13 and H017-L06)**

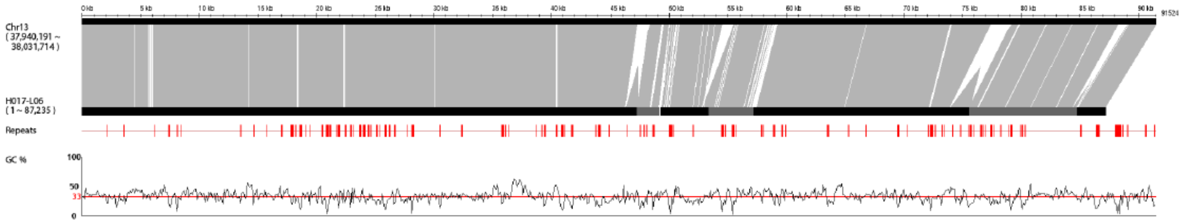


**Fig. S6** continued**.**

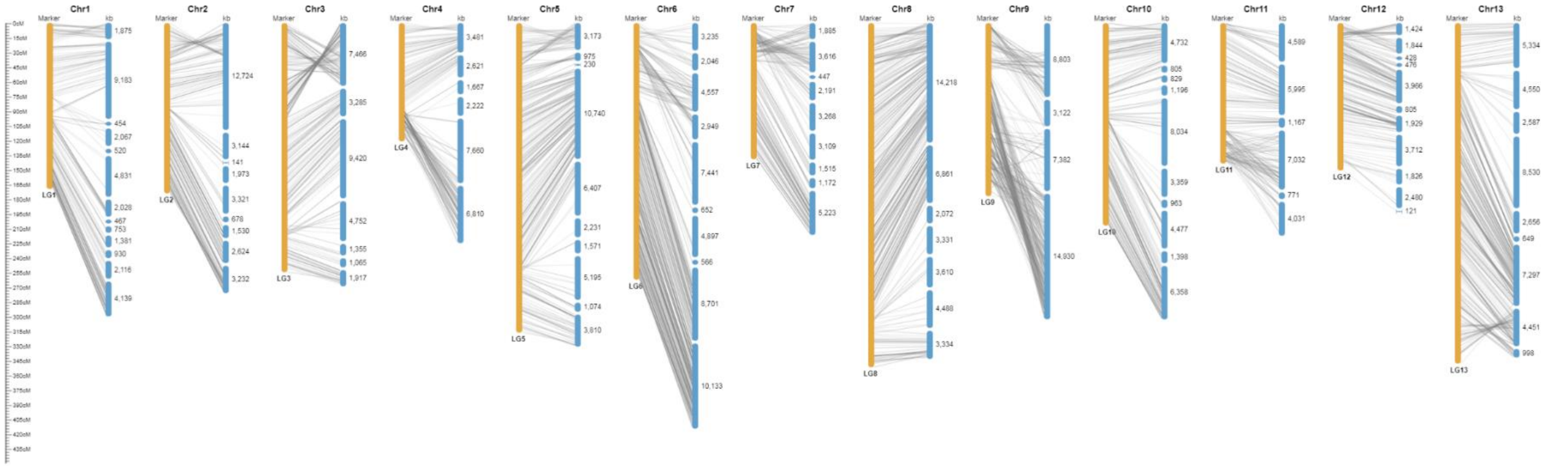


**Fig. S7.** Alignment of the genome sequence assembly with the genetic map of diploid *S. tora*. Assembled scaffolds (blue; 401.1 Mb, or 76.2% of the assembled genome sequence) were anchored to the thirteen linkage groups with 4,455 genetic markers (orange). Blue scaffolds were anchored and oriented using the Hi-C data.

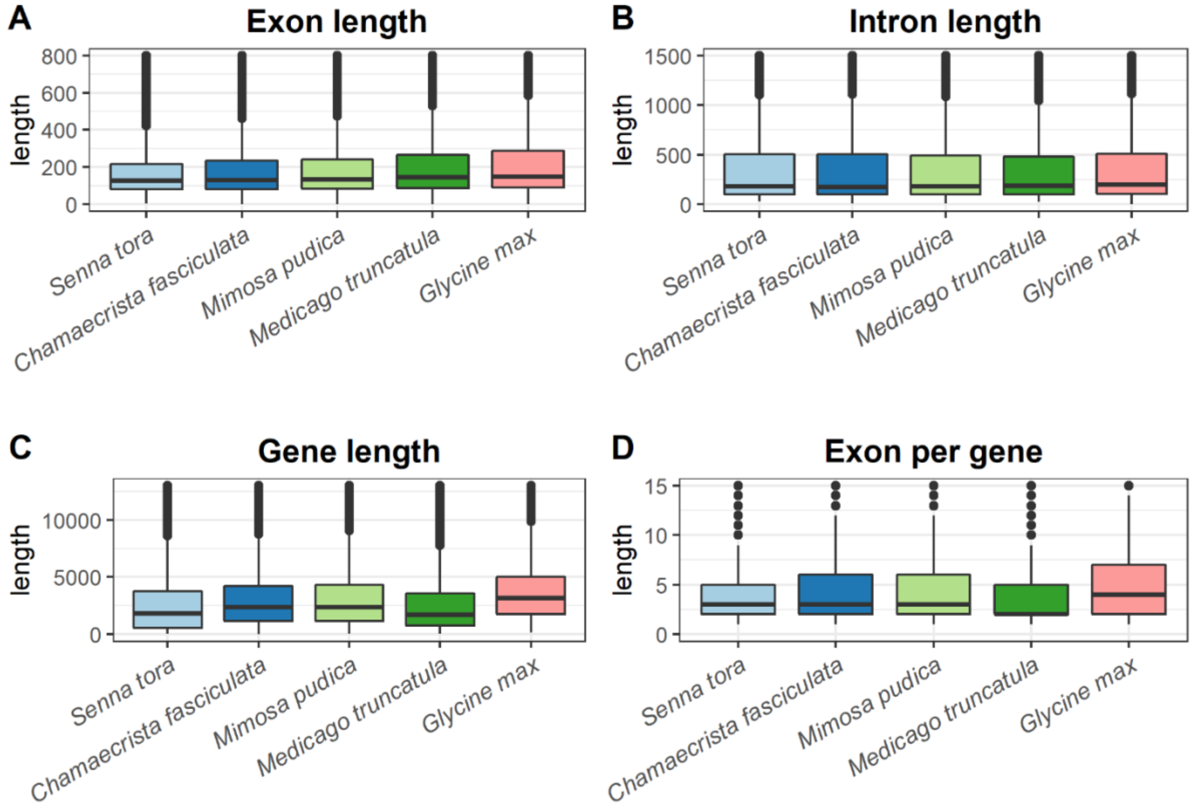


**Fig. S8.** Comparison of gene models of *S. tora*, *C. fasciculate*, *M. pudica*, *M. truncatula*, and *G. max*. The length distributions of (A) exons, (B) introns, (C) genes, and (D) exon per gene are shown.

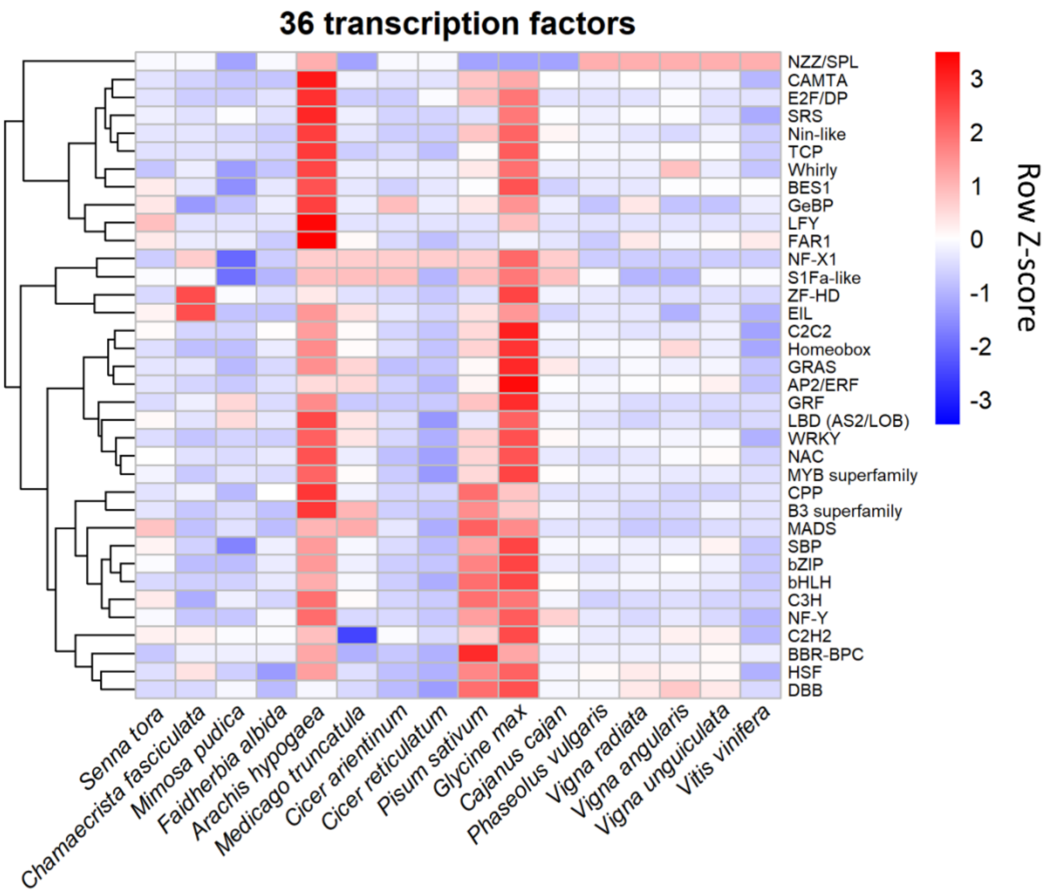


**Fig. S9.** Transcription factor genes in *S. tora* and 15 other species. Heatmap was drawn for the number of transcription factors for each genome using pheatmap R package v1.0.12 with clustering_method="ward.D2" and scale="row".

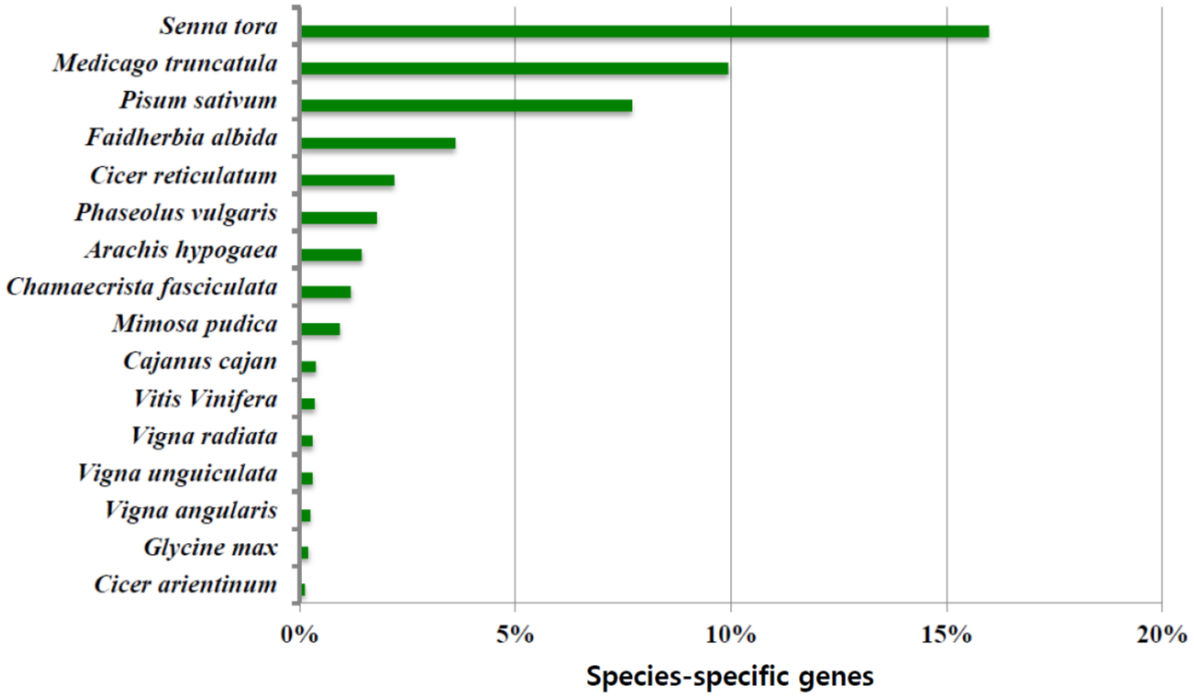


**Fig. S10.** Distribution of species-specific genes from 16 plant  species.

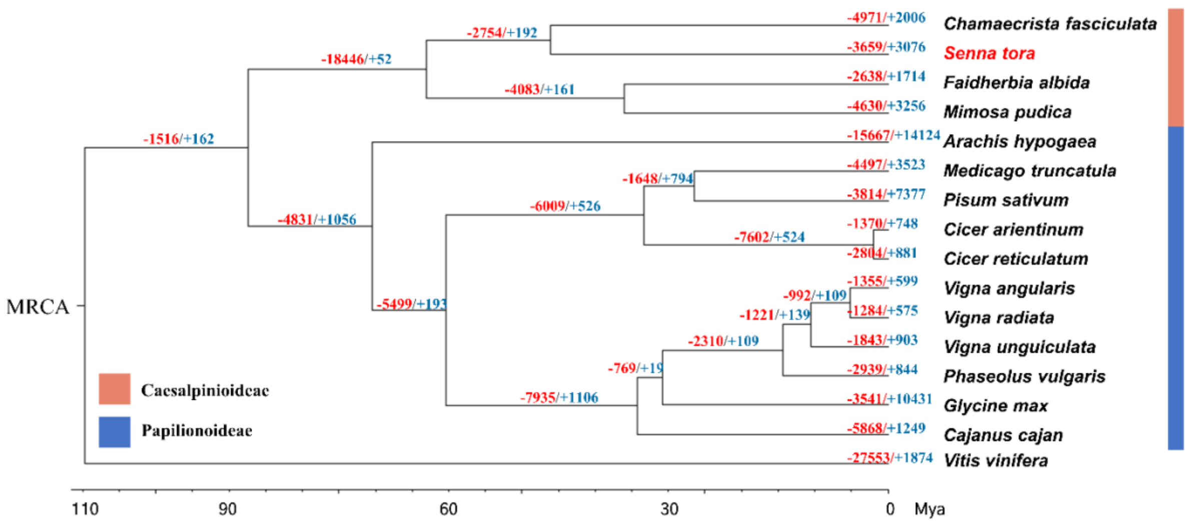


**Fig. S11.** Phylogenetic relationship and the expansion and contraction of gene families between Caesalpinioideae and Papilionoideae. Numbers on the branches show the number of gene gains (+, blue) and losses (-, red). The divergence times (MYA: Million Years Ago) are indicated by the scale bar at the bottom. *V. vinifera* was used as an outgroup. MRCA: Most Recent Common Ancestor.

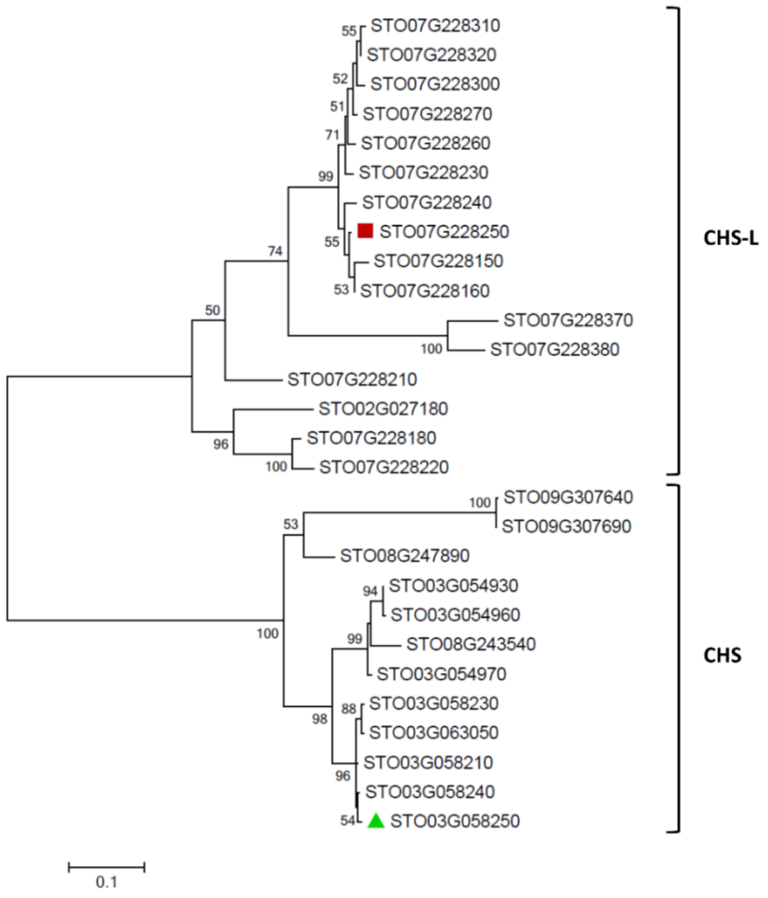


**Fig. S12.** The phylogenetic tree of CHS (12 genes) and CHS-L (16 genes) gene families in *S. tora*. This phylogenetic tree was generated using the maximum likelihood (ML) method with 1,000 bootstraps by MEGA v7.0 (https://www.megasoftware.net/), after alignment of predicted amino acid sequences by MUSCLE. Bootstrap support values (≥50%). Red square and green triangle indicate the CHS-L and CHS enzymes that were biochemically characterized in this study.

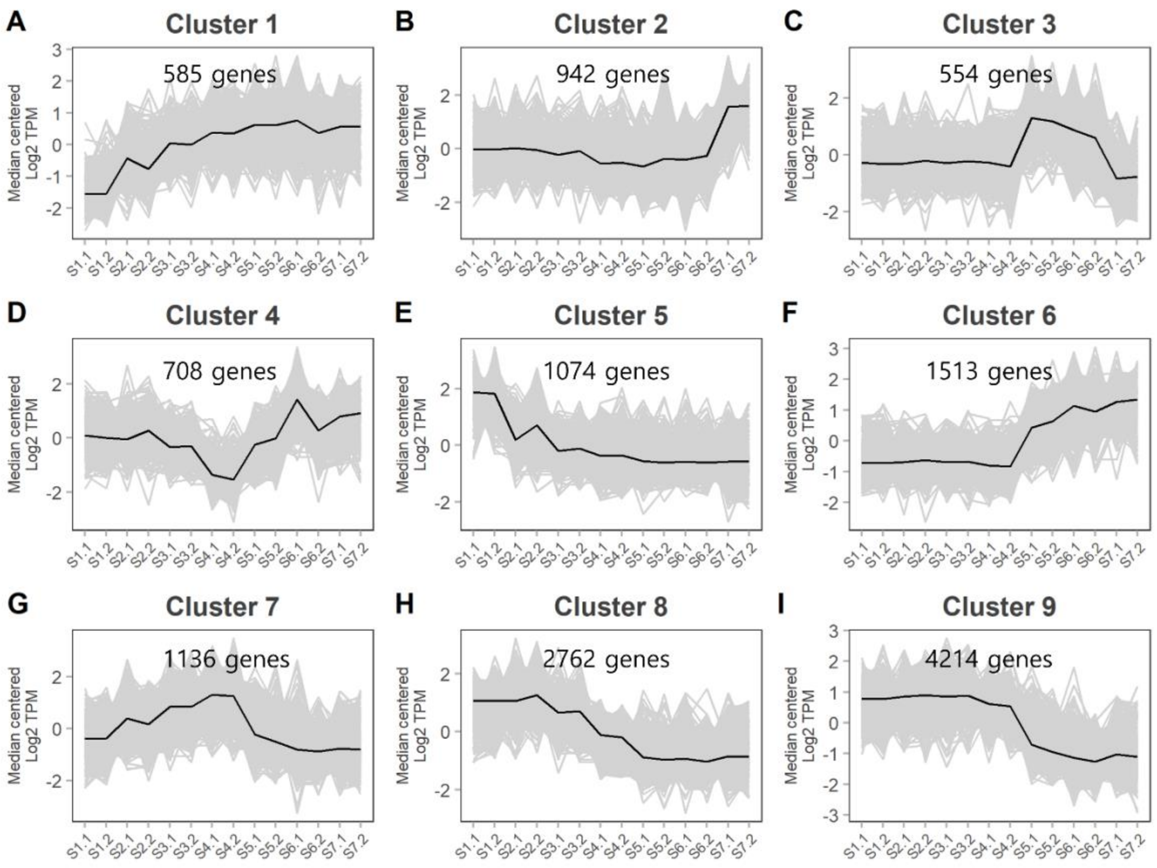


**Fig. S13.** Scaled transcript expression profiles (in transcripts per million, TPM) of representative gene co-expression clusters during seed development in *S. tora*. Numbers in x-axis indicate seed stages (S1 to S7) and two biological replicates (.1 and .2).

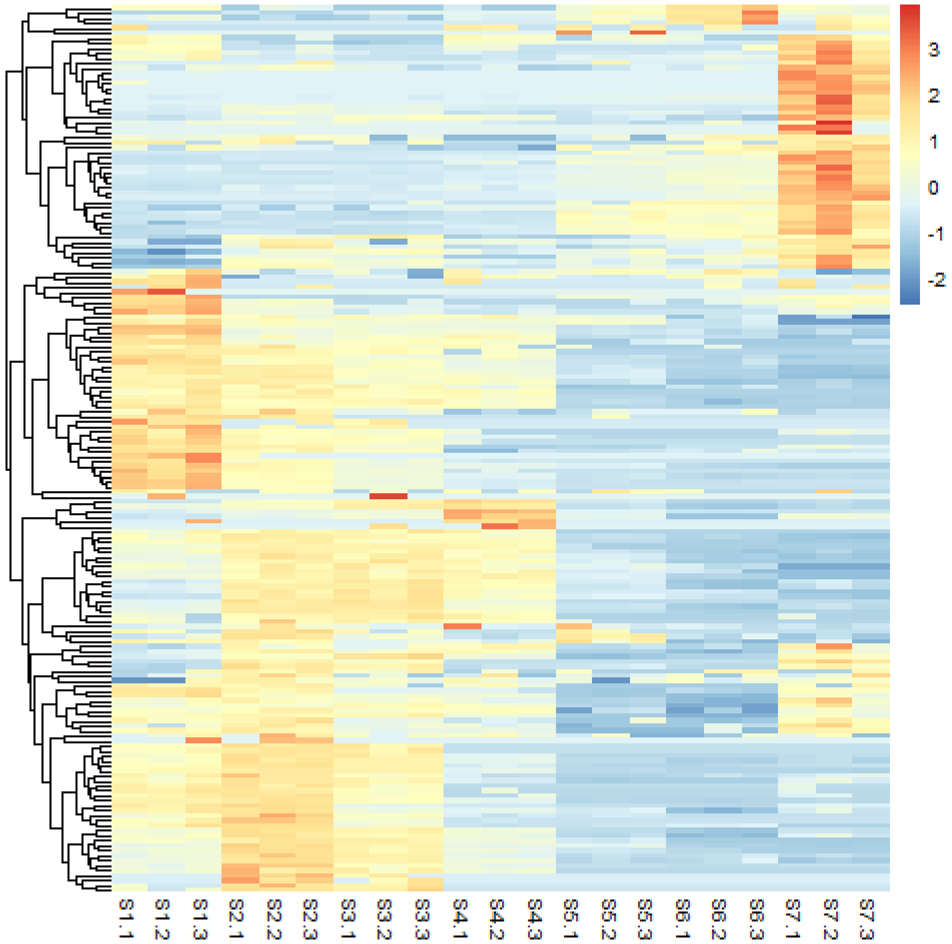


**Fig. S14.** Hierarchical clustering analysis (HCA) of primary metabolites from seven developmental stages of *S. tora* seeds. Heatmap was drawn for the relative area of 178 putative metabolites using pheatmap R package v1.0.12 with hierarchical clustering. X-axis labels indicate seed stages (S1 to S7) and three biological replicates (.1, .2, .3).

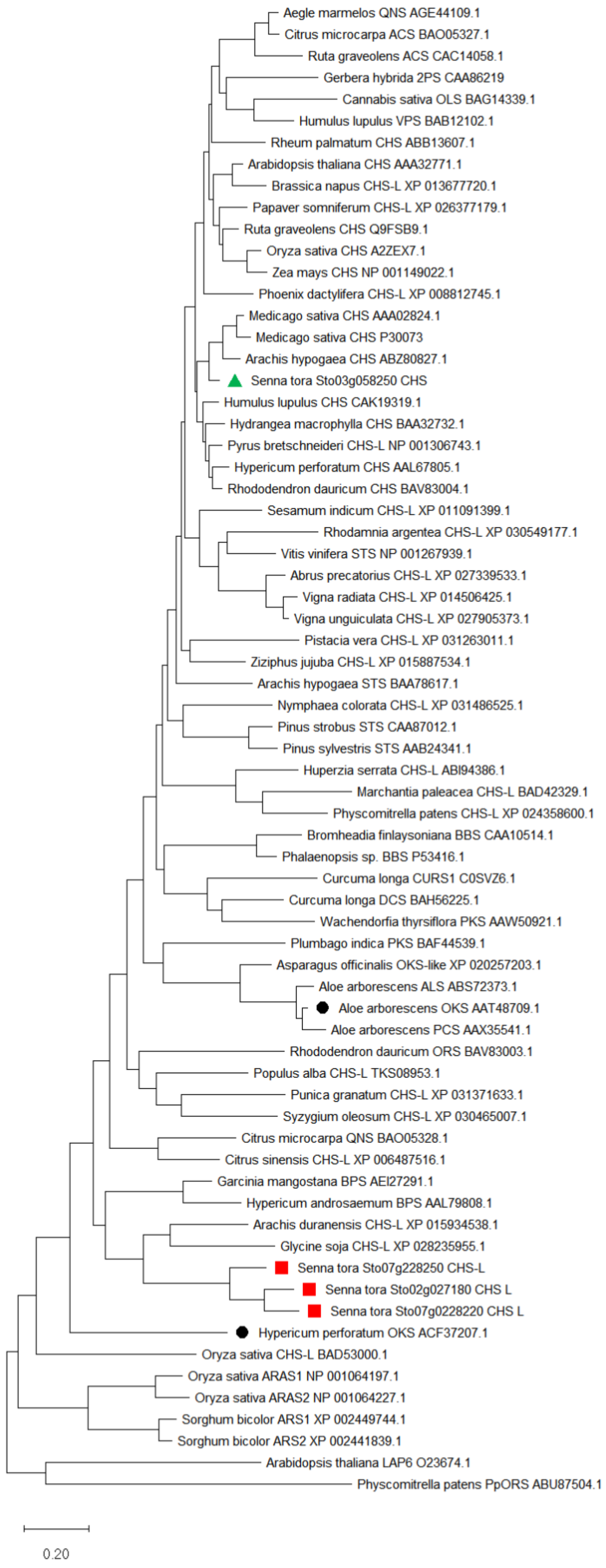
Aegle marmelos QNS AGE44109.1
Citrus microcarpa ACS BAO05327.1
Ruta graveolens ACS CAC14058.1
Gerbera hybrida 2PS CAA86219
Cannabis sativa OLS BAG14339.1
Humulus lupulus VPS BAB12102.1
Rheum palmatum CHS ABB13607.1
Arabidopsis thaliana CHS AAA32771.1
Brassica napus CHS-L XP 013677720.1
Papaver somniferum CHS-L XP 026377179.1
Ruta graveolens CHS Q9FSB9.1
Oryza sativa CHS A2ZEX7.1
Zea mays CHS NP 001149022.1
Phoenix dactylifera CHS-L XP 008812745.1
Medicago sativa CHS AAA02824.1
Medicago sativa CHS P30073
Arachis hypogaea CHS ABZ80827.1
Senna tora Sto03g058250 CHS
Humulus lupulus CHS CAK19319.1
Hydrangea macrophylla CHS BAA32732.1
Pyrus bretschneideri CHS-L NP 001306743.1
Hypericum perforatum CHS AAL67805.1
Rhododendron dauricum CHS BAV83004.1
Sesamum indicum CHS-L XP 011091399.1
Rhodamnia argentea CHS-L XP 030549177.1
Vitis vinifera STS NP 001267939.1
Abrus precatorius CHS-L XP 027339533.1
Vigna radiata CHS-L XP 014506425.1
Vigna unguiculata CHS-L XP 027905373.1
Pistacia vera CHS-L XP 031263011.1
Ziziphus jujuba CHS-L XP 015887534.1
Arachis hypogaea STS BAA78617.1
Nymphaea colorata CHS-L XP 031486525.1
Pinus strobus STS CAA87012.1
Pinus sylvestris STS AAB24341.1
Huperzia serrata CHS-L ABI94386.1
Marchantia paleacea CHS-L BAD42329.1
Physcomitrella patens CHS-L XP 024358600.1
Bromheadia finlaysoniana BBS CAA10514.1
Phalaenopsis sp. BBS P53416.1
Curcuma longa CURS1 C0SVZ6.1
Curcuma longa DCS BAH56225.1
Wachendorfia thyrsiflora PKS AAW50921.1
Plumbago indica PKS BAF44539.1
Asparagus officinalis OKS-like XP 020257203.1
Aloe arborescens ALS ABS72373.1
Aloe arborescens OKS AAT48709.1
Aloe arborescens PCS AAX35541.1
Rhododendron dauricum ORS BAV83003.1
Populus alba CHS-L TKS08953.1
Punica granatum CHS-L XP 031371633.1
Syzygium oleosum CHS-L XP 030465007.1
Citrus microcarpa QNS BAO05328.1
Citrus sinensis CHS-L XP 006487516.1
Garcinia mangostana BPS AEI27291.1
Hypericum androsaemum BPS AAL79808.1
Arachis duranensis CHS-L XP 015934538.1
Glycine soja CHS-L XP 028235955.1
Senna tora Sto07g228250 CHS-L
Senna tora Sto02g027180 CHS L
Senna tora Sto07g0228220 CHS L
Hypericum perforatum OKS ACF37207.1
Oryza sativa CHS-L BAD53000.1
Oryza sativa ARAS1 NP 001064197.1
Oryza sativa ARAS2 NP 001064227.1
Sorghum bicolor ARS1 XP 002449744.1
Sorghum bicolor ARS2 XP 002441839.1
Arabidopsis thaliana LAP6 O23674.1
Physcomitrella patens PpORS ABU87504.1
0.20

**Fig. S15.** The phylogenetic tree analysis of selected CHS and CHS-L genes in *S. tora* and other plant species. This phylogenetic tree was generated using the Maximum Likelihood method and JTT matrix-based model (*68*) after alignment of predicted amino acid sequences by MUSCLE. The tree with the highest log likelihood (-29513.52) is shown. Initial tree(s) for the heuristic search were obtained automatically by applying Neighbor-Join and BioNJ algorithms to a matrix of pairwise distances estimated using the JTT model, and then selecting the topology with superior log likelihood value. The tree is drawn to scale, with branch lengths measured in the number of substitutions per site. This analysis involved 69 amino acid sequences. There were a total of 880 positions in the final dataset. Evolutionary analyses were conducted in MEGA X (*69*). Green colored triangle and red colored rectangle represent the CHS and CHS-Ls from *S. tora* described in this study. Octaketide synthases from *Aloe arborescens* and *Hypericum perforatum* are labeled with black circle.

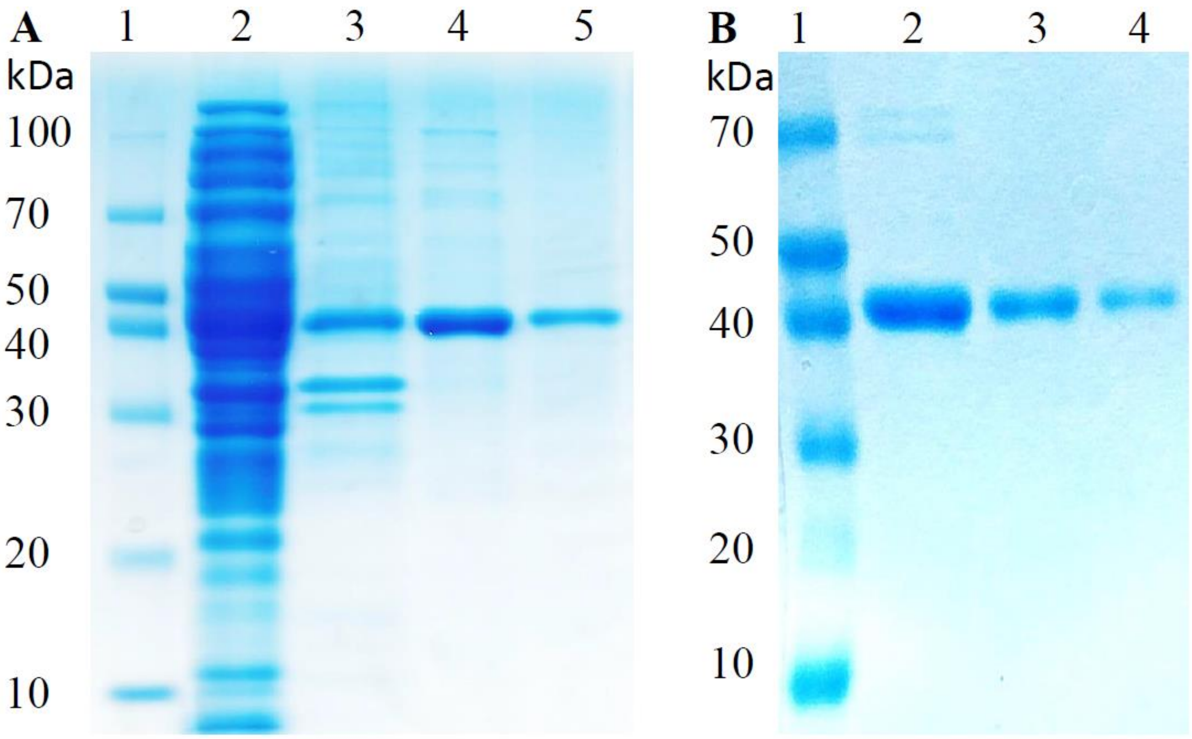


**Fig. S16.** SDS-PAGE analysis of (A) STO07G228250 (CHS-L). Lane 1: Standard protein ladder; Lane 2: total soluble fraction; Lane 3: total insoluble fraction; Lanes 4 and 5: pure protein fractions. The pure fractions of the protein were pooled and concentrated. (B) STO03G058250 (CHS). Lane 1: standard protein ladder; Lanes 2-4: pure soluble fractions. The pure fractions of the protein were pooled and concentrated.

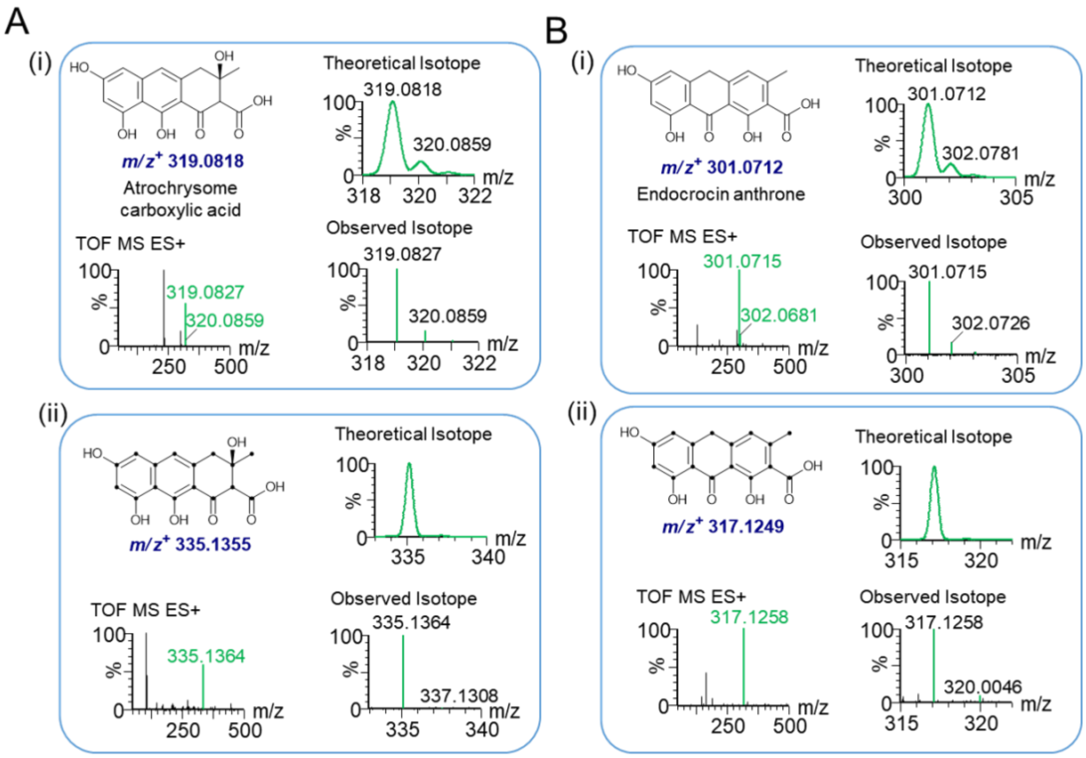


**Fig. S17. TOF ESI-MS analysis of the anthranoids generated in reaction assays.** (A) (i) TOF ESI-MS spectrum for the speculated product atrochrysome carboxylic acid with the molecular formula $C_{16}H_{14}O_7$ for which calculated theoretical exact mass was 319.0818 Da in proton adduct form. The theoretical mass isotope perfectly aligned to the observed mass isotope. (ii) ESI MS-spectrum for the speculated $^{13}C$-labelled atrochrysome carboxylic acid with molecular formula $^{13}C_{16}H_{14}O_7$ for which calculated theoretical exact mass was 335.1355 Da in proton adduct form. The theoretical mass isotope perfectly aligned to observed mass isotope. (B) (i) TOF ESI-MS spectrum for the speculated product endocrocin anthrone with molecular formula $C_{16}H_{12}O_6$ for which the calculated theoretical exact mass was 301.0712 Da in proton adduct form. The theoretical mass isotope perfectly aligned to the observed mass isotope. (ii) TOF ESI-MS spectrum for the speculated $^{13}C$-labelled endocrocin anthrone with molecular formula $^{13}C_{16}H_{12}O_6$ for which calculated theoretical exact mass was 317.1249 Da in proton adduct form. The theoretical mass isotope perfectly aligned to the observed mass isotope.

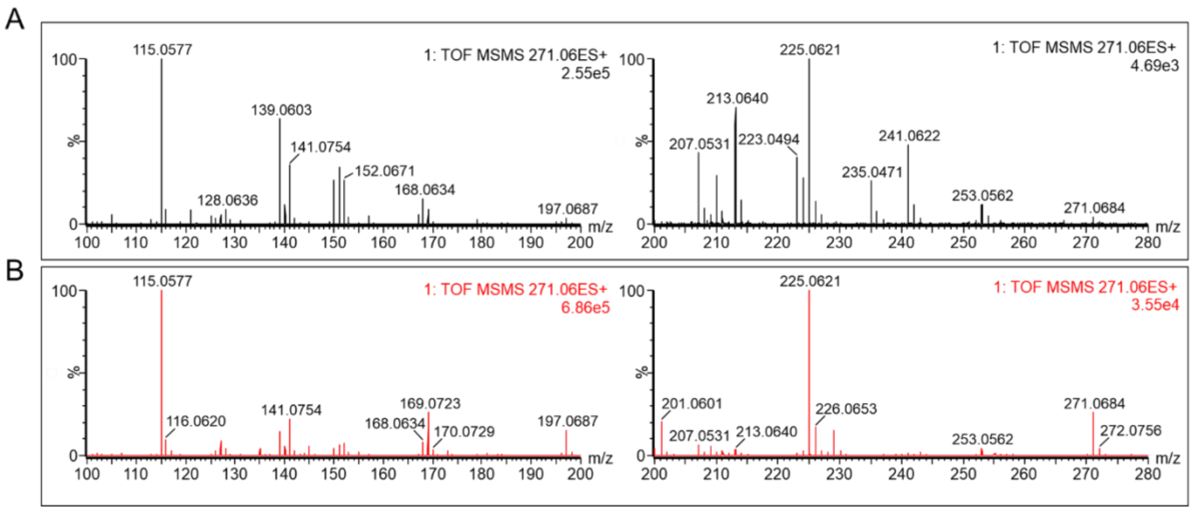


**Fig. S18.** ESI-MS-MS analysis of precursor ion 271 for (A) aloe-emodin and (B) emodin.

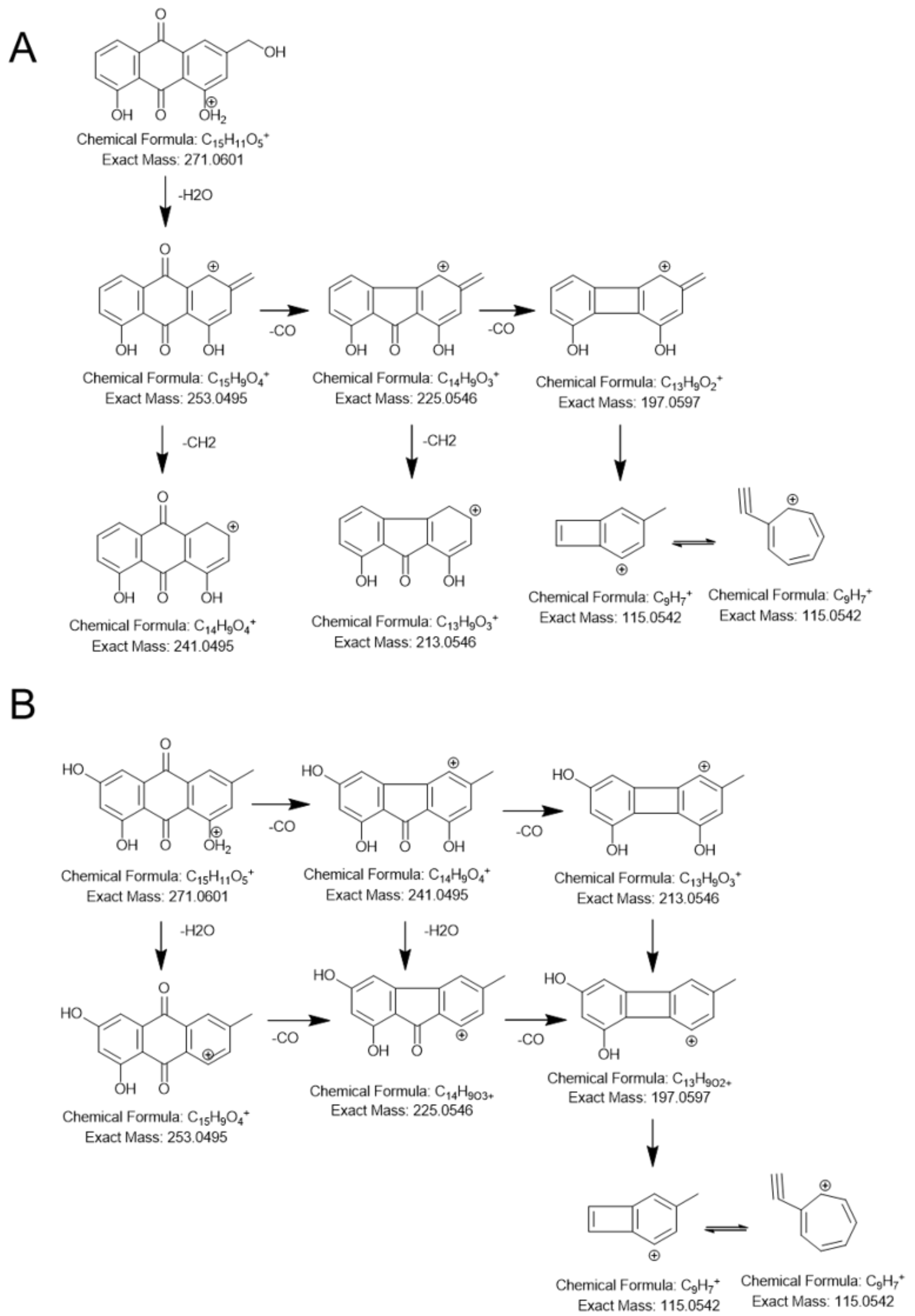


**Fig. S19.** Proposed MS-MS fragmentation of standard (A) emodin and (B) aloe-emodin.

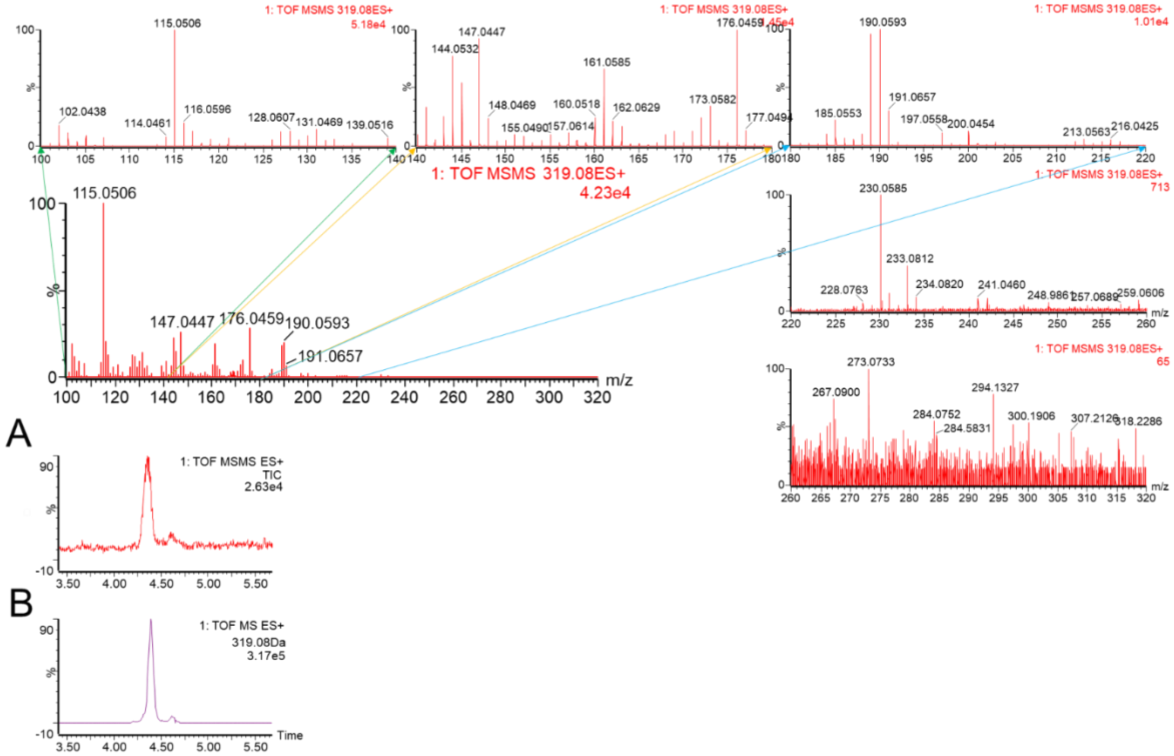


**Fig. S20.** ESI-MS$_2$ analysis of precursor ion 319.08. (A) Total ion chromatogram (TIC) of ESI-MS$_2$ for precursor ion 319 and (B) extracted ion chromatogram (EIC) of ESI-MS for mass 319.08 Da. Mass spectra are shown in expanded view.

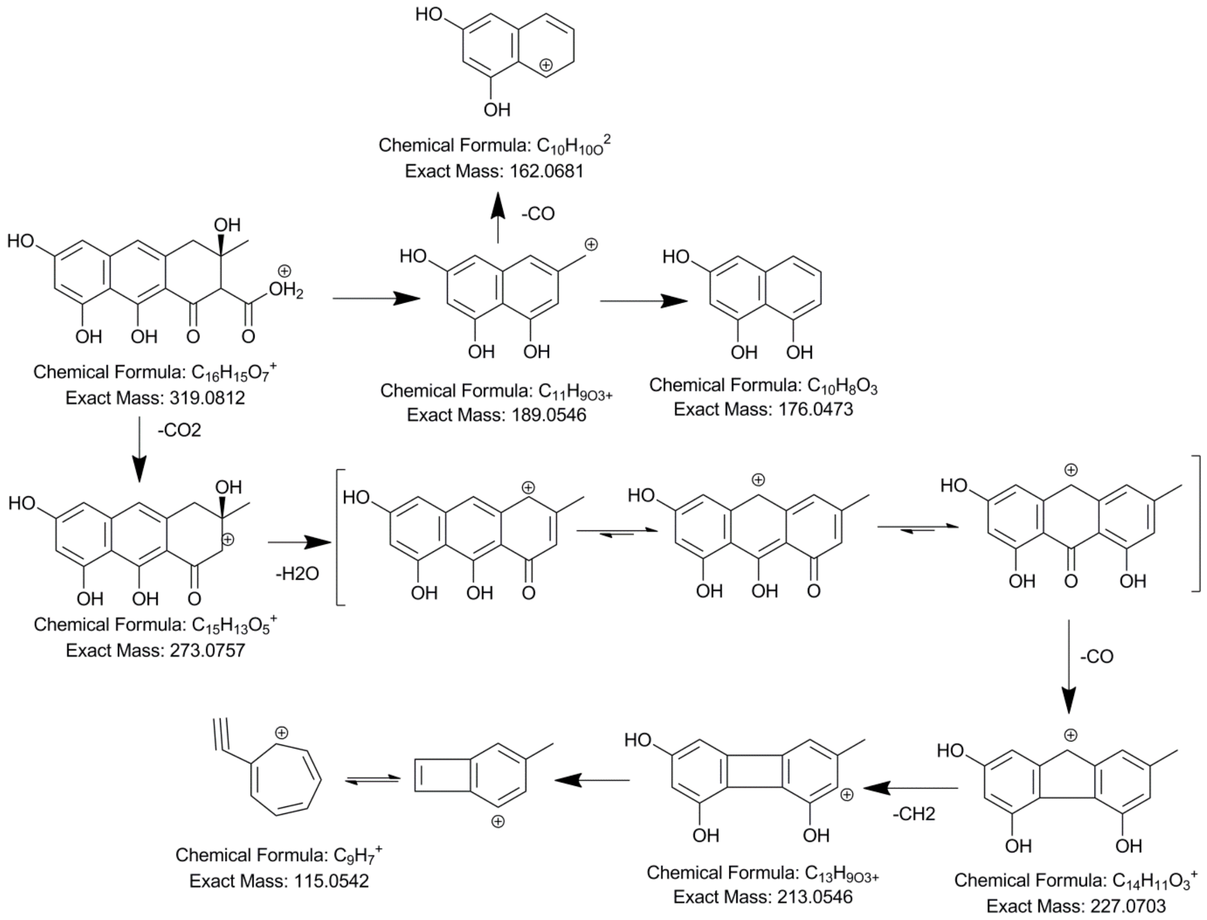


**Fig. S21.** Proposed MS-MS fragmentation of atrochrysome carboxylic acid based on MS-MS fragmentation pattern of emodin and aloe–emodin under identical MS conditions.

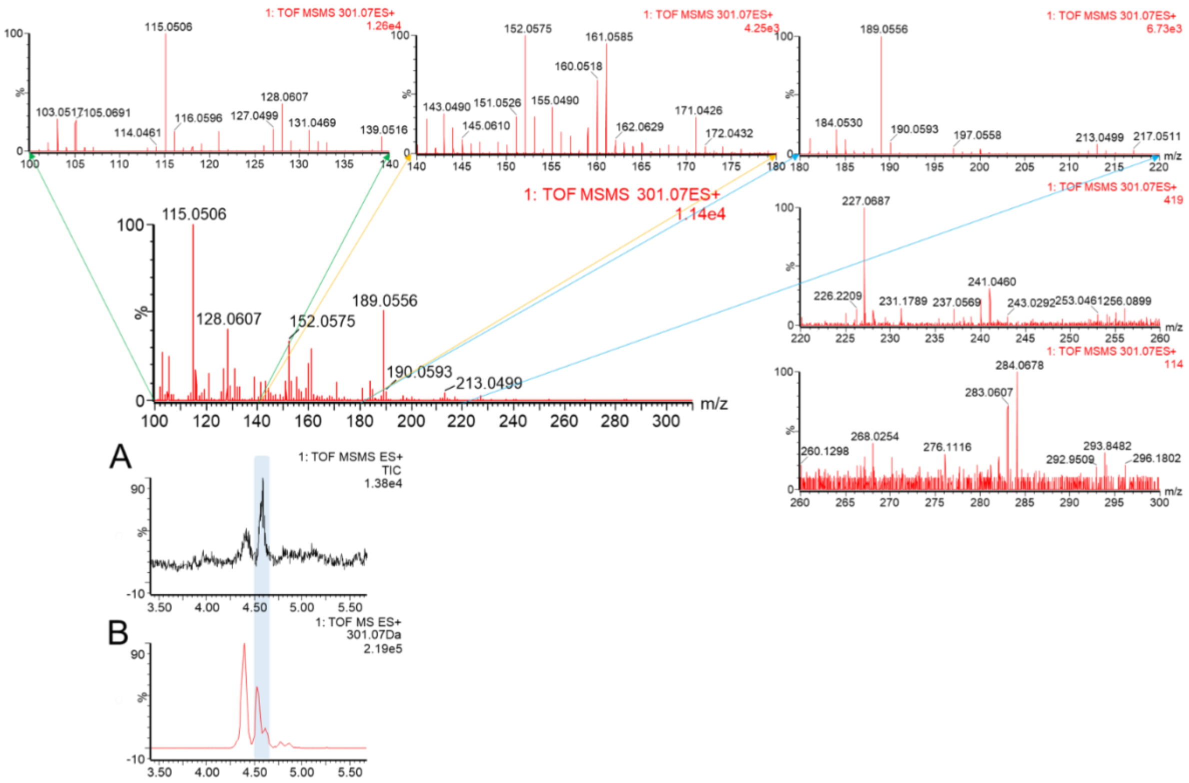


**Fig. S22.** ESI-MS2 analysis of precursor ion 301. (A) TIC of ESI-MS2 for precursor ion 301 and (B) EIC of ESI-MS for mass 301.08 Da. ESI- MS2 spectra are shown in expanded view.

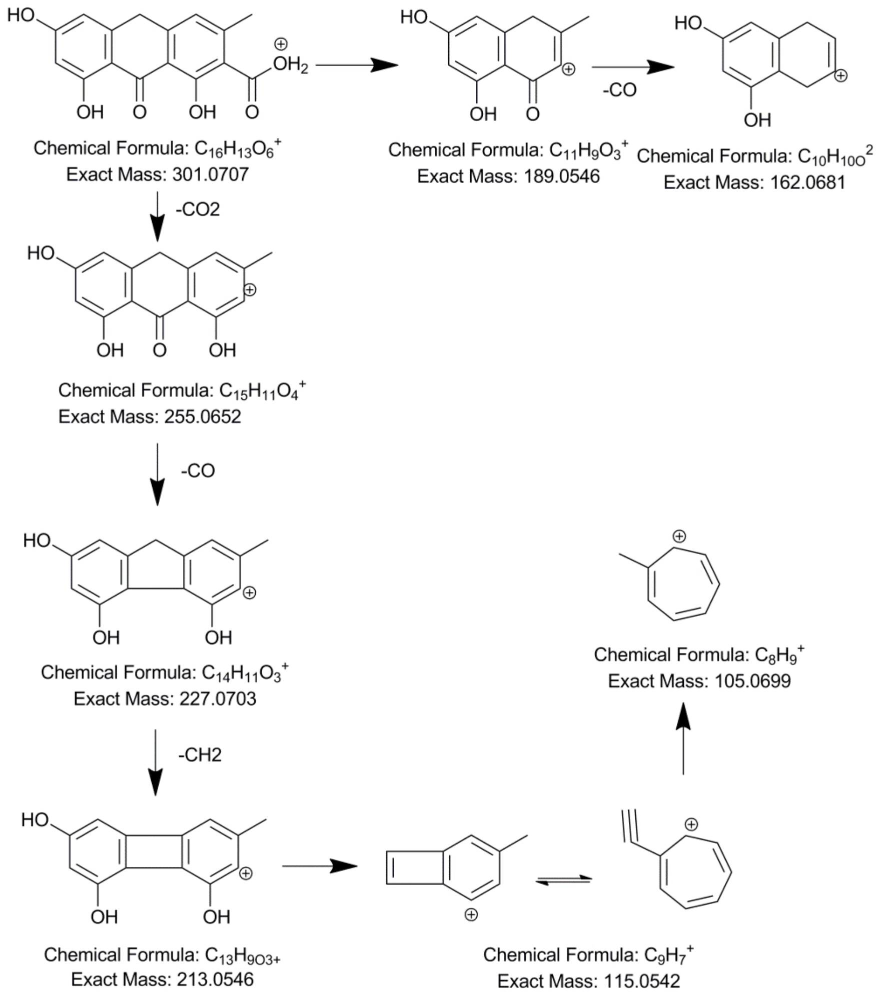


**Fig. S23.** Proposed MS-MS fragmentation of endocrocin anthrone based on MS-MS fragmentation pattern of emodin and aloe–emodin under identical MS conditions.

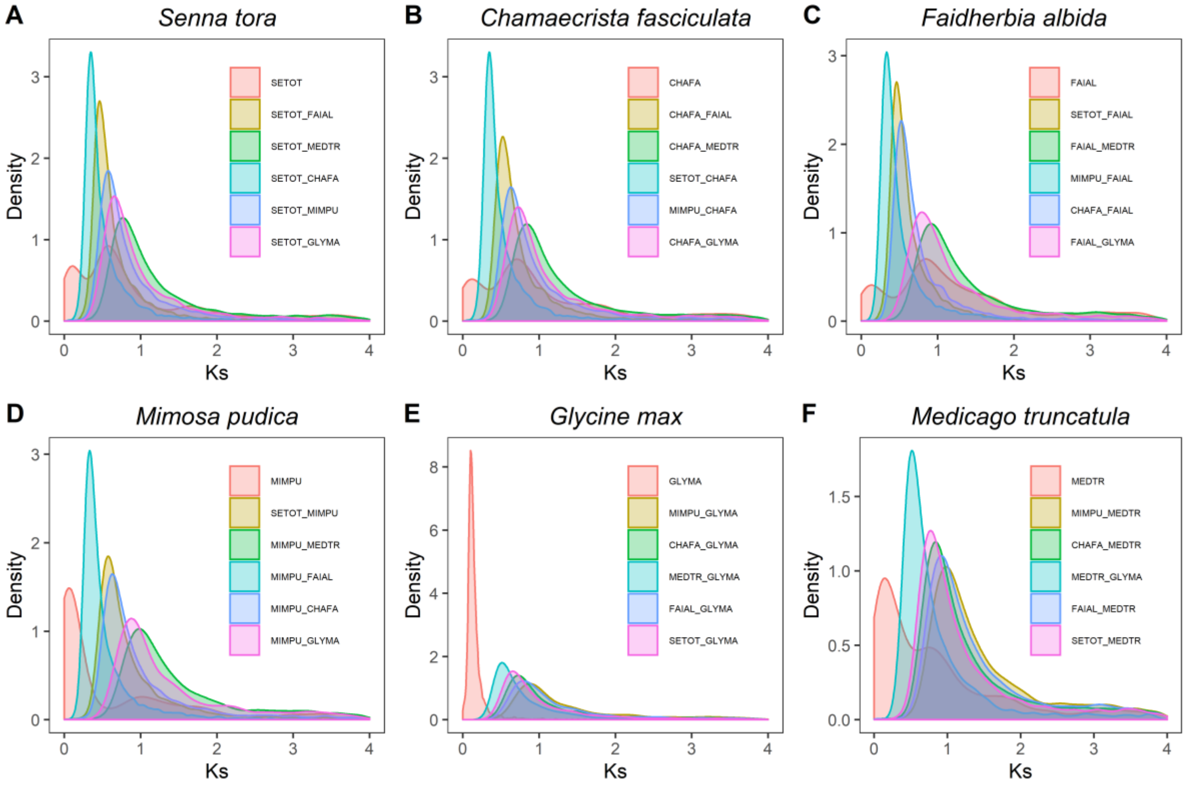


**Fig. S24.** Distribution of the synonymous substitution rate (Ks) among (A) *S. tora*, (B) *C. fasciculata*, (C) *F. albida*, (D) *M. pudica*, (E) *G. max*, and (F) *M. truncatula* in intra- and inter-genomic comparisons. Intra-genomic analysis indicates the WGD events and inter-genomic comparisons represent the divergence. *S. tora* (SETOT), *C. fasciculata* (CHAFA), *M. pudica* (MIMPU), *F. albida* (FAIAL), *M. truncatula* (MEDTR), *G. max* (GLYMA).

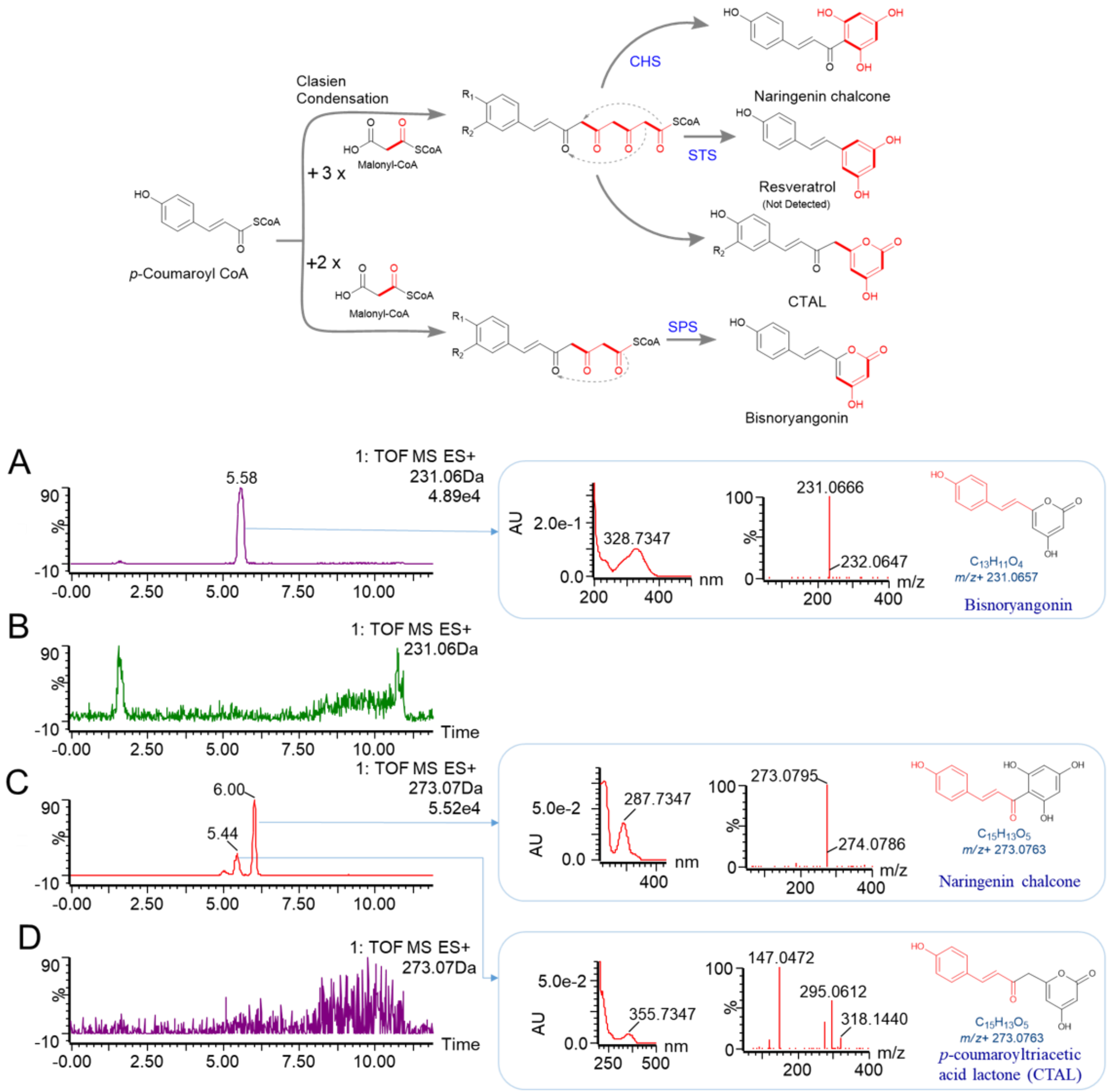


**Fig. S25.** High resolution TOF ESI-MS analysis of STO03G058250 (CHS) reaction mixture containing *p*-coumaroyl-CoA as a starter unit. Extracted ion chromatogram for mass 231.06 Da in (A) reaction mixture and (B) reaction mixture containing only malonyl-CoA. EIC for mass 273.07 Da in (C) reaction mixture and (D) reaction mixture containing only malonyl-CoA. UV-VIS, ESI-MS spectrum along with their possible structure of each peak is shown with arrow. Scheme in the top shows folding of the polyketide chain to produce different metabolites by different type III PKS enzymes.

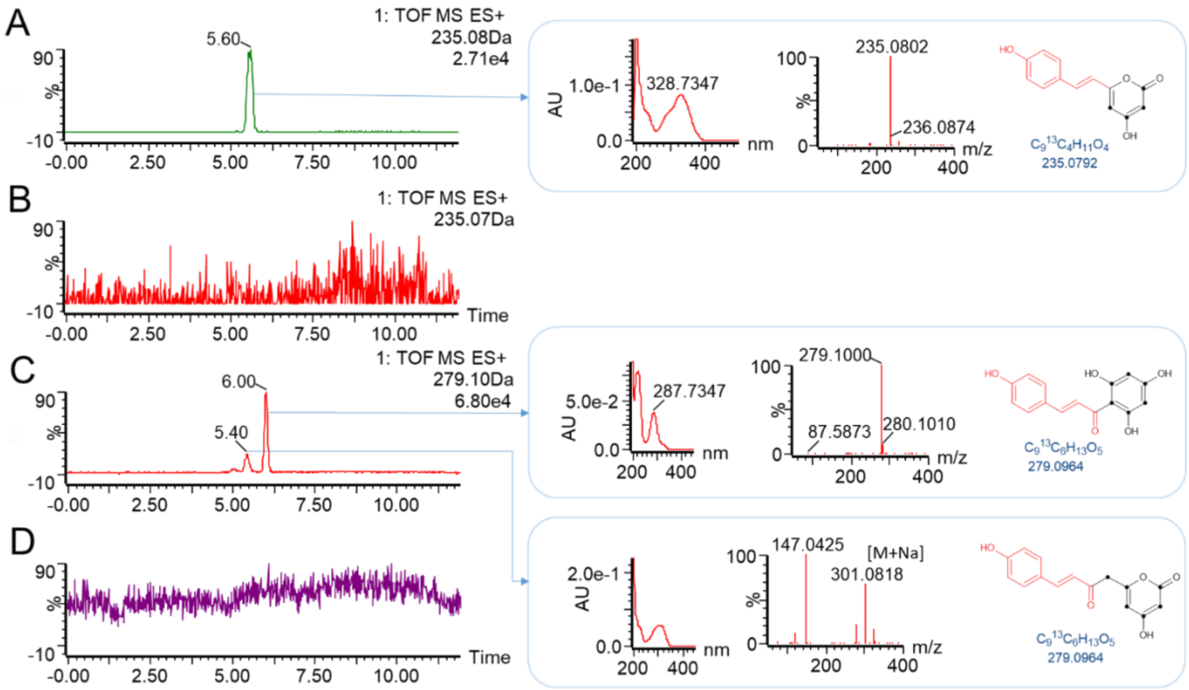


**Fig. S26.** High resolution TOF ESI-MS analysis of STO03G058250 (CHS) reaction mixture containing *p*-coumaroyl-CoA as a starter unit and $^{13}C_3$-malonyl-CoA as extender substrate. Extracted ion chromatogram for mass 235.07 Da in (A) reaction mixture and (B) reaction mixture containing only $^{13}C_3$-malonyl-CoA. EIC for mass 279.09 Da in (C) reaction mixture and (D) reaction mixture containing only $^{13}C_3$-malonyl-CoA. UV-VIS and ESI-MS spectrum, along with their possible structure of each peak, are shown with arrow.

**Supplementary Tables:**

**Table S1.** Sequence information of the *de novo* genome

| Platform | Library | Data (Gb) | Depth (x) | Q20 (%) | Q30 (%) |
|---|---|---|---|---|---|
| HiSeq | PE† _200bp-1 | 42.14 | 77.03 | 97.0 | 92.8 |
| | PE_200bp-2 | 44.51 | 81.36 | 96.9 | 92.4 |
| | PE_200bp-3 | 46.27 | 84.58 | 96.6 | 91.9 |
| | MP† _3kb-1 | 22.65 | 41.40 | 94.0 | 86.8 |
| | MP_3kb-2 | 24.38 | 44.56 | 93.2 | 85.5 |
| | MP_3kb-3 | 23.23 | 42.46 | 92.8 | 85.1 |
| | MP_5kb-1 | 25.46 | 46.54 | 93.4 | 86.0 |
| | MP_5kb-2 | 19.22 | 35.13 | 92.2 | 84.2 |
| | MP_5kb-3 | 20.40 | 37.29 | 92.1 | 84.0 |
| | MP_10kb-1 | 43.49 | 79.50 | 93.8 | 87.4 |
| | MP_10kb-2 | 43.17 | 78.91 | 94.4 | 88.6 |
| | MP_10kb-3 | 44.17 | 80.74 | 93.0 | 86.7 |
| | MP_20kb-1 | 45.56 | 83.28 | 92.3 | 85.8 |
| | MP_20kb-2 | 45.80 | 83.72 | 91.6 | 85.0 |
| | MP_20kb-3 | 44.49 | 81.33 | 92.5 | 86.7 |
| MiSeq | PE_500bp-1 | 13.90 | 25.41 | 89.7 | 80.4 |
| | PE_500bp-2 | 14.46 | 26.43 | 90.1 | 80.0 |
| | PE_500bp-3 | 14.63 | 26.74 | 91.0 | 80.9 |
| | Total | 577.93 | 1056.41 | -- | -- |

† PE and MP represent pair-end and mate-pair, respectively.

| Platform | Data (Gb) | Depth (x) | Long reads (bp) | Average length (bp) |
|---|---|---|---|---|
| PacBio RS II system | 40.88 | 74.73 | 3,330,429 | 12,275 |
| PacBio Sequel system | 39.13 | 71.53 | 3,487,455 | 11,221 |
| Total | 80.01 | 146.26 | 6,817,884 | -- |

**Table S2.** Comparison of assembly results of four assemblers

| | SOAPdenovo2 | | Allpaths-LG | | Platanus | | FALCON |
|---|---|---|---|---|---|---|---|
| | Contigs | Scaffolds (1k over) | Contigs | Scaffolds (1k over) | Contigs | Scaffolds (1k over) | Contigs |
| **No** | 47,840 | 1,270 | 28,597 | 5,323 | 16,941 | 4,550 | 957 |
| **Length (bp)** | 563,756,844 | 602,528,808 | 574,497,226 | 603,199,927 | 523,669,775 | 536,028,526 | 533,300,920 |
| **N50 (bp)** | 27,794 | 2,221,313 | 71,672 | 1,298,531 | 248,367 | 2,358,844 | 3,966,958 |
| **Largest (bp)** | 249,767 | 13,596,213 | 1,234,493 | 9,174,740 | 1,329,697 | 14,262,124 | 14,930,962 |
| **Average (bp)** | 11,784 | 474,432 | 20,089 | 113,319 | 30,911 | 117,808 | 557,263 |
| **N (bp)** | | 30,295,731 | | 28,700,086 | | 12,358,751 | -- |

**Table S3.** BUSCO evaluation of the assembly from Platanus and FALCON Assemblies

| Category | Platanus | | FALCON | |
|---|---|---|---|---|
| | Number | Percentage (%) | Number | Percentage (%) |
| Complete BUSCOs | 1,305 | 90.6 | 1,358 | 94.3 |
| Complete and single-copy BUSCOs | 1,156 | 80.3 | 1,244 | 86.4 |
| Complete and duplicated BUSCOs | 149 | 10.3 | 114 | 7.9 |
| Fragmented BUSCOs | 66 | 4.6 | 16 | 1.1 |
| Missing BUSCOs | 69 | 4.8 | 66 | 4.6 |
| Total BUSCO groups searched | 1,440 | 100 | 1,440 | 100 |

**Table S4.** Comparison of the assembled pseudochromosomes and 10 independently sequenced BACs

| No. | BAC ID | BAC length (bp) | Chr ID | Coverage (%) | Identities (%) |
|---|---|---|---|---|---|
| 1 | B050-B02 | 82,000 | Chr3 | 100 | 99.94 |
| 2 | B020-G17 | 92,904 | Chr5 | 100 | 99.93 |
| 3 | B016-D19 | 66,759 | Chr6 | 100 | 99.95 |
| 4 | H036-G09 | 81,063 | Chr6 | 98.4 | 99.88 |
| 5 | H001-O11 | 88,210 | Chr8 | 100 | 99.49 |
| 6 | H019-A05 | 87,111 | Chr8 | 100 | 99.90 |
| 7 | H024-N24 | 81,044 | Chr8 | 100 | 99.81 |
| 8 | H002-L14 | 107,545 | Chr9 | 100 | 99.97 |
| 9 | B011-O10 | 92,399 | Chr10 | 100 | 99.79 |
| 10 | H017-L06 | 87,235 | Chr13 | 96 | 99.83 |

**Table S5.** Statistical analysis of the functional annotations of protein-coding genes in *S. tora*

| | NO. | Percent (%) |
|---|---|---|
| **TOTAL** | 45,268 | |
| **NR** | 31,010 | 68.50 |
| **GO** | 25,453 | 56.23 |
| **KEGG** | 17,450 | 38.55 |
| **SWISSPROT** | 23,533 | 51.99 |
| **EGGNOG** | 17,786 | 39.29 |
| **NO HIT** | 13,708 | 30.28 |

**Table S6.** Annotation of non-coding RNA genes in *S. tora*

| Type | | Copy | Average length (bp) | Total length (bp) | % of genome |
|---|---|---|---|---|---|
| rRNA | 5S | 432 | 119.13 | 51463 | 0.009 |
| | 5.8S | 106 | 155.24 | 16455 | 0.003 |
| | 18S | 107 | 1839.17 | 196791 | 0.036 |
| | 28S | 107 | 3968.50 | 424629 | 0.078 |
| tRNA | | 839 | 74.33 | 62363 | 0.011 |

**Table S7.** Annotation of long non-coding RNA (lncRNA) genes in *S. tora* (See large tables file)

**Table S8.** Transcription factor genes in *S. tora* and 15 other plant species

| TF family | Species‡ | | | | | | | | | | | | | | | |
|---|---|---|---|---|---|---|---|---|---|---|---|---|---|---|---|---|
| | SETOT | CHAFA | MIMPU | FALAL | ARHYP | MEDTR | CICAR | CICRE | PISSA | GLYMA | CAJCA | PHAVU | VIGRA | VIGAN | VIGUN | VITVI |
| **AP2/ERF** | 169 | 159 | 151 | 167 | 209 | 210 | 157 | 140 | 192 | 341 | 184 | 179 | 184 | 187 | 195 | 148 |
| **B3** | 69 | 51 | 63 | 48 | 195 | 125 | 56 | 49 | 147 | 114 | 77 | 70 | 60 | 62 | 77 | 67 |
| **BBR-BPC** | 3 | 5 | 5 | 5 | 10 | 2 | 3 | 2 | 16 | 10 | 5 | 5 | 5 | 5 | 6 | 5 |
| **BES1** | 9 | 7 | 3 | 7 | 16 | 7 | 6 | 7 | 8 | 16 | 6 | 7 | 7 | 8 | 8 | 8 |
| **bHLH** | 137 | 130 | 131 | 149 | 225 | 159 | 128 | 104 | 271 | 300 | 166 | 154 | 158 | 154 | 149 | 125 |
| **bZIP** | 72 | 53 | 53 | 67 | 104 | 68 | 58 | 55 | 111 | 130 | 67 | 63 | 69 | 73 | 69 | 56 |
| **C2C2** | 102 | 86 | 86 | 101 | 135 | 102 | 86 | 80 | 114 | 180 | 98 | 94 | 91 | 92 | 95 | 68 |
| **C2H2** | 13 | 13 | 12 | 12 | 16 | 1 | 12 | 10 | 15 | 23 | 12 | 11 | 11 | 13 | 13 | 8 |
| **C3H** | 53 | 33 | 46 | 41 | 77 | 50 | 41 | 39 | 77 | 76 | 47 | 40 | 42 | 43 | 41 | 40 |
| **CAMTA** | 7 | 6 | 5 | 5 | 25 | 8 | 7 | 7 | 13 | 15 | 9 | 8 | 9 | 8 | 8 | 4 |
| **CPP** | 6 | 7 | 3 | 8 | 22 | 7 | 5 | 5 | 18 | 12 | 6 | 6 | 6 | 5 | 5 | 6 |
| **DBB** | 6 | 6 | 7 | 5 | 7 | 6 | 5 | 4 | 12 | 13 | 7 | 7 | 8 | 9 | 8 | 6 |
| **E2F/DP** | 7 | 6 | 6 | 7 | 17 | 6 | 6 | 8 | 11 | 14 | 7 | 7 | 7 | 8 | 7 | 7 |
| **EIL** | 9 | 18 | 5 | 5 | 14 | 10 | 7 | 6 | 10 | 14 | 6 | 7 | 7 | 4 | 7 | 4 |
| **FAR1** | 71 | 44 | 45 | 25 | 214 | 60 | 33 | 17 | 36 | 48 | 41 | 25 | 71 | 52 | 59 | 69 |
| **GeBP** | 7 | 4 | 5 | 6 | 11 | 6 | 8 | 6 | 7 | 9 | 6 | 5 | 7 | 5 | 5 | 6 |
| **GRAS** | 54 | 54 | 46 | 52 | 80 | 67 | 47 | 48 | 60 | 99 | 63 | 55 | 58 | 56 | 57 | 48 |
| **GRF** | 9 | 10 | 13 | 9 | 17 | 8 | 8 | 8 | 14 | 22 | 10 | 10 | 9 | 9 | 9 | 9 |
| **Homeobox** | 73 | 63 | 63 | 77 | 122 | 85 | 73 | 64 | 98 | 150 | 78 | 82 | 82 | 96 | 78 | 55 |
| **HSF** | 25 | 33 | 23 | 16 | 42 | 25 | 21 | 19 | 45 | 50 | 28 | 30 | 32 | 31 | 30 | 19 |
| **LBD(AS2/LOB)** | 57 | 50 | 63 | 51 | 92 | 61 | 49 | 35 | 51 | 87 | 54 | 50 | 47 | 50 | 47 | 48 |
| **LFY** | 2 | 1 | 1 | 1 | 4 | 1 | 1 | 1 | 1 | 2 | 1 | 1 | 1 | 1 | 1 | 1 |
| **MADS** | 127 | 63 | 80 | 60 | 134 | 140 | 83 | 52 | 179 | 157 | 81 | 79 | 67 | 69 | 77 | 78 |
| **MYB** | 270 | 219 | 253 | 241 | 467 | 288 | 222 | 162 | 330 | 506 | 287 | 270 | 265 | 258 | 260 | 245 |
| **NAC** | 103 | 91 | 88 | 94 | 174 | 96 | 77 | 64 | 120 | 173 | 94 | 90 | 93 | 101 | 105 | 85 |
| **NF-X1** | 2 | 3 | 1 | 2 | 3 | 3 | 3 | 3 | 3 | 4 | 3 | 2 | 2 | 2 | 2 | 2 |
| **NF-Y** | 10 | 7 | 7 | 10 | 19 | 8 | 8 | 7 | 16 | 20 | 13 | 9 | 8 | 9 | 8 | 6 |
| **Nin-like** | 11 | 11 | 10 | 9 | 29 | 11 | 9 | 9 | 18 | 26 | 14 | 12 | 11 | 10 | 12 | 9 |
| **NZZ/SPL** | 1 | 1 | 0 | 1 | 2 | 0 | 1 | 1 | 0 | 0 | 0 | 2 | 2 | 2 | 2 | 2 |
| **S1Fa-like** | 2 | 2 | 0 | 1 | 3 | 3 | 3 | 1 | 3 | 4 | 3 | 2 | 1 | 1 | 2 | 2 |
| **SBP** | 25 | 19 | 11 | 22 | 34 | 23 | 20 | 17 | 33 | 43 | 23 | 23 | 22 | 22 | 25 | 18 |
| **SRS** | 10 | 9 | 11 | 9 | 27 | 10 | 8 | 8 | 9 | 21 | 11 | 10 | 11 | 11 | 9 | 5 |
| **TCP** | 24 | 24 | 24 | 22 | 57 | 21 | 23 | 19 | 29 | 52 | 28 | 27 | 27 | 28 | 28 | 21 |
| **Whirly** | 2 | 3 | 1 | 2 | 8 | 3 | 3 | 3 | 4 | 7 | 3 | 3 | 3 | 5 | 3 | 2 |
| **WRKY** | 80 | 70 | 76 | 74 | 166 | 107 | 78 | 60 | 116 | 173 | 98 | 91 | 93 | 91 | 94 | 61 |
| **ZF-HD** | 17 | 42 | 21 | 18 | 24 | 18 | 17 | 15 | 18 | 43 | 20 | 19 | 18 | 18 | 18 | 17 |
| **Total** | **1,644** | **1,403** | **1,418** | **1,429** | **2,801** | **1,805** | **1,372** | **1,135** | **2,205** | **2,954** | **1,656** | **1,555** | **1,594** | **1,598** | **1,619** | **1,360** |

‡ represents *S. tora* (SETOT), *C. fasciculata* (CHAFA), *M. pudica* (MIMPU), *F. albida* (FAIAL), *A. hypogaea* (ARHYP), *M. truncatula* (MEDTR), *C. arientinum* (CICAR), *C. reticulatum* (CICRE), *P. sativum* (PISSA), *G. max* (GLYMA), *C. cajan* (CAJCA), *P. vulgaris* (PHAVU), *V. radiata* (VIGRA), *V. angularis* (VIGAN), *V. unguiculata* (VIGUN), *V. vinifera* (VITVI)

**Table S9.** Statistics of orthologs and paralogs in Fabaceae and *Vitis vinifera*

| Species | Genome size (Mbp) | No. coding genes | No. ortholog genes | No. paralog genes | No. species-specific genes | No. uncertain genes | No. non-species-specific genes | Species-specific genes (%) | P-value |
|---|---|---|---|---|---|---|---|---|---|
| ***Senna tora*** | **526.40** | **45,268** | **23,461** | **8,938** | **7,231** | **5,638** | **38,037** | **15.97** | **< 0.00001** |
| *Chamaecrista fasciculata* | 429.27 | 32,832 | 24,314 | 3,275 | 388 | 4,855 | 32,444 | 1.18 | < 0.00001 |
| *Mimosa pudica* | 557.21 | 33,108 | 24,883 | 3,264 | 304 | 4,657 | 32,804 | 0.92 | < 0.00001 |
| *Faidherbia albida* | 653.73 | 28,979 | 21,624 | 818 | 1,040 | 5,497 | 27,939 | 3.59 | < 0.00001 |
| *Arachis hypogaea* | 2539.16 | 83,709 | 44,585 | 27,369 | 1,202 | 10,553 | 82,507 | 1.44 | |
| *Medicago truncatula* | 412.92 | 41,939 | 29,619 | 7,200 | 4,165 | 9,460 | 46,279 | 9.93 | |
| *Cicer arientinum* | 530.89 | 35,754 | 23,044 | 400 | 36 | 1,482 | 24,926 | 0.10 | |
| *Cicer reticulatum* | 416.66 | 26,404 | 21,425 | 1,514 | 573 | 2,892 | 25,831 | 2.17 | |
| *Pisum sativum* | 3920.16 | 57,835 | 35,155 | 11,388 | 4,447 | 6,845 | 53,388 | 7.69 | |
| *Glycine max* | 979.05 | 71,219 | 41,070 | 1,110 | 137 | 4,477 | 46,657 | 0.19 | |
| *Cajanus cajan* | 592.97 | 41,387 | 25,788 | 899 | 148 | 2,284 | 28,971 | 0.36 | |
| *Phaseolus vulgaris* | 521.08 | 32,720 | 24,899 | 374 | 579 | 2,282 | 27,555 | 1.77 | |
| *Vigna radiata* | 463.64 | 42,284 | 24,438 | 439 | 123 | 1,961 | 26,838 | 0.29 | |
| *Vigna angularis* | 467.30 | 37,769 | 24,491 | 305 | 85 | 1,753 | 26,549 | 0.23 | |
| *Vigna unguiculata* | 519.07 | 41,173 | 26,063 | 749 | 117 | 1,301 | 28,113 | 0.28 | |
| *Vitis vinifera* | 486.20 | 41,208 | 21,064 | 2,002 | 140 | 2,470 | 28,971 | 0.36 | |

**Table S10.** All significantly enriched biological process GO and KEGG categories of expanding gene families in *S. tora* compared to other 15 species (*C. fasciculata*, *M. pudica*, *F. albida*, *A. hpogaea*, *M. truncatula*, *C. arientinum*, *C. reticulatum*, *P. Sativum*, *G. max*, *C.cajan*, *P. vulgaris*, *V. radiata*, *V. angularis*, *V. unguiculata*, and *V. vinifera*)

| GO ID | GO description | Number of genes | FDR-corrected p-value |
|---|---|---|---|
| GO:0006259 | DNA metabolic process | 71 | 2.24E-23 |
| GO:0006278 | RNA-dependent DNA replication | 40 | 1.95E-14 |
| GO:0032197 | transposition, RNA-mediated | 31 | 8.47E-13 |
| GO:0019076 | viral release from host cell | 27 | 1.69E-11 |
| GO:0006310 | DNA recombination | 63 | 9.02E-10 |
| GO:0071897 | DNA biosynthetic process | 38 | 3.25E-09 |
| GO:0044238 | primary metabolic process | 31 | 4.37E-08 |
| GO:0043170 | macromolecule metabolic process | 23 | 1.05E-07 |
| GO:0090501 | RNA phosphodiester bond hydrolysis | 27 | 4.12E-07 |
| GO:0051276 | chromosome organization | 23 | 1.20E-05 |
| GO:0006807 | nitrogen compound metabolic process | 27 | 1.41E-05 |
| GO:0010584 | pollen exine formation | 17 | 5.26E-05 |
| GO:0090304 | nucleic acid metabolic process | 20 | 8.05E-05 |
| GO:0001560 | regulation of cell growth by extracellular stimulus | 7 | 8.86E-05 |
| GO:0000723 | telomere maintenance | 31 | 0.000102 |
| GO:0009698 | phenylpropanoid metabolic process | 11 | 0.000352 |
| GO:0010930 | negative regulation of auxin mediated signaling pathway | 5 | 0.000388 |
| GO:0044237 | cellular metabolic process | 15 | 0.001238 |
| GO:0006974 | cellular response to DNA damage stimulus | 23 | 0.001586 |
| GO:0009987 | cellular process | 27 | 0.004064 |
| GO:0046777 | protein autophosphorylation | 41 | 0.005557 |
| GO:0050792 | regulation of viral process | 3 | 0.012344 |
| GO:0046246 | terpene biosynthetic process | 3 | 0.012344 |

| | | | |
|---|---|---|---|
| GO:0009793 | embryo development ending in seed dormancy | 37 | 0.019838 |
| GO:0019761 | glucosinolate biosynthetic process | 6 | 0.019838 |
| GO:0006725 | cellular aromatic compound metabolic process | 4 | 0.020891 |
| GO:0009791 | post-embryonic development | 16 | 0.021442 |
| GO:0006366 | transcription from RNA polymerase II promoter | 17 | 0.023933 |
| GO:0009615 | response to virus | 10 | 0.034372 |
| GO:1902290 | positive regulation of defense response to oomycetes | 4 | 0.034372 |
| **KEGG ID** | **KEGG description** | **Number of genes** | **FDR-corrected p-value** |
| PATH:ko00940 | phenylpropanoid biosynthesis | 25 | 1.03E-10 |
| PATH:ko00130 | ubiquinone and other terpenoid quinone biosynthesis | 15 | 2.08E-09 |
| PATH:ko03020 | RNA polymerase | 8 | 0.001468 |
| PATH:ko00943 | isoflavonoid biosynthesis | 5 | 0.009071 |

**Table S11.** List of statistically significant expanded/contracted CHS-L and CHS subfamilies in *S. tora* and 15 related species

| Subfamilies | | Species‡ | | | | | | | | | | | | | | | |
|---|---|---|---|---|---|---|---|---|---|---|---|---|---|---|---|---|---|
| | | SETOT | CHAFA | MIMPU | FAIAL | ARHYP | MEDTR | CICAR | CICRE | PISSA | GLYMA | CAJCA | PHAVU | VIGRA | VIGAN | VIGUN | VITVI |
| CHS-L | E/C* | Rapid_E† | E | - | - | E | - | - | - | C | - | C | - | - | - | - | C |
| | Gain/Loss gene count | 12 | 1 | - | - | 1 | - | - | - | -1 | - | -1 | - | - | - | - | -1 |
| | Orthologous gene | 16 | 5 | 0 | 0 | 2 | 1 | 0 | 0 | 0 | 1 | 0 | 0 | 0 | 0 | 0 | 0 |
| CHS | E/C* | C | E | C | C | Rapid_E† | Rapid_E† | - | - | Rapid_C† | E | C | C | - | C | Rapid_E† | Rapid_E† |
| | Gain/Loss gene count | -3 | 7 | -2 | -6 | 32 | 11 | - | - | -10 | 4 | -2 | -2 | - | -1 | 7 | 17 |
| | Orthologous gene | 12 | 22 | 11 | 7 | 48 | 21 | 6 | 6 | 0 | 15 | 9 | 8 | 8 | 7 | 17 | 39 |

* E/C represents expansion/contraction. † Rapid_E and Rapid_C indicate rapid expansion and rapid contraction (see Methods). ‡ represents *S. tora* (SETOT), *C. fasciculata* (CHAFA), *M. pudica* (MIMPU), *F. albida* (FAIAL), *A. hypogaea* (ARHYP), *M. truncatula* (MEDTR), *C. arientinum* (CICAR), *C. reticulatum* (CICRE), *P. sativum* (PISSA), *G. max* (GLYMA), *C. cajan* (CAJCA), *P. vulgaris* (PHAVU), *V. radiata* (VIGRA), *V. angularis* (VIGAN), *V. unguiculata* (VIGUN), *V. vinifera* (VITVI).

**Table S12.** List of anthraquinone standards used in this study

| No. | Name | Formula | Average MS(Da) | Monoisotopic MS(Da) | $[M+H]^+$ | $[M-H]^-$ | Production for MRM |
|---|---|---|---|---|---|---|---|
| 1 | Glucoaurantio-obtusin | $C_{23}H_{24}O_{12}$ | 492.436 | 492.127 | 493.1341 | 491.1195 | 242.1 |
| 2 | Obtusin | $C_{18}H_{16}O_7$ | 344.321 | 344.09 | 345.0969 | 343.08233 | 313 |
| 3 | Chryso-obtusin | $C_{19}H_{18}O_7$ | 358.348 | 358.105 | 359.1125 | 357.09798 | 342.1 |
| 4 | Chrysophanol | $C_{15}H_{10}O_4$ | 254.242 | 254.058 | 255.0652 | 253.05063 | 225.1 |
| 5 | Emodin | $C_{15}H_{10}O_5$ | 270.241 | 270.053 | 271.0601 | 269.04555 | 225.1 |
| 6 | Gluco-obtusifolin | $C_{22}H_{22}O_{10}$ | 446.41 | 446.121 | 447.1286 | 445.11402 | 268.2 |
| 7 | Aurantio-obtusin | $C_{17}H_{14}O_7$ | 330.294 | 330.074 | 331.0812 | 329.06668 | 298.9 |
| 8 | Aloe-emodin | $C_{15}H_{10}O_5$ | 270.241 | 270.053 | 271.0601 | 269.04555 | 240.1 |
| 9 | Physcion | $C_{16}H_{12}O_5$ | 284.268 | 284.068 | 285.0758 | 283.0612 | 239.9 |
| 10 | Obtusinfolin | $C_{16}H_{12}O_5$ | 284.268 | 284.068 | 285.0758 | 283.0612 | 92.1 |

**Table S13.** Contents of ten anthraquinone compounds at different stages of *S. tora* seed development (Stage1-Stage7)

| Compounds | Amount of anthraquinones (ug/g)* | | | | | | |
|---|---|---|---|---|---|---|---|
| | **Stage1** | **Stage2** | **Stage3** | **Stage4** | **Stage5** | **Stage6** | **Stage7** |
| Glucoaurantio-obtusin | N.D | 4.03±0.30 | 35.45±1.48 | 224. 80±20.80 | 1009.73±66.67 | 1144.47±24.91 | 296.00±11.01 |
| Obtusin | N.D | N.D | N.D | N.D | N.D | N.D | 68.22 ±8.00 |
| Chryso-obtusin | N.D | N.D | N.D | 0.82±0.18 | 1.76±0.58 | 1.77±0.49 | 54.66±8.03 |
| Chrysophanol | 3.24±0.05 | 13.98±0.65 | 18.23±1.67 | 19.69±1.89 | 46.23±2.25 | 30.21±1.63 | 12.89±0.98 |
| Emodin | 215.5±12.51 | 74.39±9.64 | 47.34±4.32 | 13.68±0.49 | 13.27±1.19 | 4.13±1.24 | 5.52±1.21 |
| Gluco-obtusifolin | N.D | N.D | 3.07±1.14 | 31.16±1.21 | 193.04±12.45 | 224.13±4.92 | 56.78±3.79 |
| Aurantio-obtusin | N.D | N.D | N.D | 0.57±0.14 | 3.57±0.41 | 4.94±0.76 | 312.20±73.63 |
| Aloe-emodin | N.D | N.D | N.D | N.D | 1.47±0.19 | 1.79±0.40 | 0.93±0.19 |
| Physcion | 6.84±0.18 | 10.19±0.72 | 8.21±0.71 | 4.16±0.32 | 5.97±1.14 | 3.55±0.34 | 1.81±0.25 |
| Obtusifolin | N.D | 0.13±0.02 | 0.20±0.06 | 1.10±0.13 | 1.04±0.08 | 1.22±0.04 | 84.45±3.27 |
| **Total** | **225.58±12.74** | **102.72±11.33** | **112.5±9.38** | **295.98±25.16** | **1,276.08±84.96** | **1,416.21±34.73** | **893.46±110.36** |

* indicates mean of three biological replicate experiments.

**Table S14.** Significantly enriched molecular function GO categories of the gene expression cluster 6 during seed development

| GO ID | GO description | Number of genes | FDR-corrected P-value |
|---|---|---|---|
| GO:0016758 | transferase activity, transferring hexosyl groups | 39 | 0.00001 |
| GO:0010427 | abscisic acid binding | 10 | 0.00142 |
| GO:0008194 | UDP-glycosyltransferase activity | 20 | 0.00514 |
| GO:0051536 | iron-sulfur cluster binding | 15 | 0.01000 |
| GO:0004864 | protein phosphatase inhibitor activity | 9 | 0.01000 |
| GO:0038023 | signaling receptor activity | 9 | 0.01000 |
| GO:0000978 | RNA polymerase II proximal promoter sequence-specific DNA binding | 8 | 0.01362 |
| GO:0004842 | ubiquitin-protein transferase activity | 39 | 0.03018 |
| GO:0010295 | (+)-abscisic acid 8'-hydroxylase activity | 4 | 0.03018 |
| GO:0036402 | proteasome-activating ATPase activity | 4 | 0.03018 |
| GO:0047216 | inositol 3-alpha-galactosyltransferase activity | 3 | 0.03472 |
| GO:0016705 | oxidoreductase activity, acting on paired donors, with incorporation or reduction of molecular oxygen | 37 | 0.04860 |

**Table S15.** 178 putative metabolites from seven different seed development in *S. tora*. (See large tables file.)

**Table S16.** Quantitative estimation of 69 primary metabolites from seven different seed development in *S. tora*. (See large tables file.)

**Table S17.** Mapping statistics of Illumina, RNA-Seq, and Iso-Seq data in this study

| | Library | No. sequencing reads | No. high quality reads | Mapping rate (%) |
|---|---|---|---|---|
| Genome-Seq | PE_200-1 | 279,095,332 | 185,792,369 | 99.75 |
| | PE_200-2 | 294,819,090 | 205,902,564 | 99.75 |
| | PE_200-3 | 306,485,872 | 213,085,841 | 99.74 |
| RNA-Seq | Seed | 22,986,190 | 22,122,628 | 84.92 |
| | Flower | 69,614,064 | 54,230,644 | 85.15 |
| | Leaf | 45,662,632 | 39,833,132 | 73.21 |
| | Stem | 66,562,030 | 52,162,636 | 84.45 |
| | root | 46,386,464 | 40,201,556 | 89.62 |
| | Stage1-1 (S†) | 44,230,592 | 39,568,687 | 94.05 |
| | Stage1-2 (S) | 41,572,424 | 37,710,345 | 94.83 |
| | Stage2-1 (S) | 40,494,100 | 34,912,700 | 93.32 |
| | Stage2-2 (S) | 38,637,340 | 34,932,019 | 89.88 |
| | Stage3-1 (S) | 44,752,688 | 40,510,133 | 95.49 |
| | Stage3-2 (S) | 39,833,596 | 36,001,604 | 92.36 |
| | Stage4-1 (S) | 44,470,644 | 40,165,885 | 95.50 |
| | Stage4-2 (S) | 37,584,824 | 34,093,193 | 93.71 |
| | Stage5-1 (S) | 38,394,282 | 34,885,044 | 95.16 |
| | Stage5-2 (S) | 42,445,844 | 38,388,021 | 83.86 |
| | Stage6-1 (S) | 38,629,298 | 34,615,713 | 95.46 |
| | Stage6-2 (S) | 41,408,918 | 37,541,325 | 94.69 |
| | Stage7-1 (S) | 38,492,940 | 35,313,423 | 87.78 |
| | Stage7-2 (S) | 38,251,900 | 35,065,516 | 85.39 |
| Iso-Seq | Consensus seq. | 768,745 | 118,390 | 97.18 |

† indicates seed.

**Table S18.** Summary of *S. tora* genetic map and anchored contigs

| LG | No. of markers | Genetics length (cM) | No. of contig | Physical length (bp) | Anchored contig list |
|---|---|---|---|---|---|
| 1 | 235 | 167.2 | 13 | 30,974,193 | c102, c129, c14, c150, c152, c174, c175, c23, c60, c61, c64, c76, c78 |
| 2 | 280 | 171.89 | 9 | 29,371,314 | c154, c256, c36, c37, c41, c44, c48, c68, c97 |
| 3 | 358 | 251.46 | 7 | 29,263,208 | c103, c119, c15, c34, c40, c73, c84 |
| 4 | 226 | 119.36 | 6 | 24,463,614 | c30, c52, c56, c6, c82, c87 |
| 5 | 417 | 312.65 | 10 | 35,412,120 | c117, c12, c125, c2, c217, c26, c43, c59, c7, c95 |
| 6 | 487 | 259.11 | 10 | 45,180,853 | c1, c124, c13, c161, c17, c38, c46, c62, c79, c83 |
| 7 | 257 | 137.4 | 9 | 22,430,049 | c109, c11, c177, c29, c35, c45, c58, c69, c98 |
| 8 | 508 | 347.47 | 7 | 37,916,848 | c21, c28, c32, c33, c42, c5, c63 |
| 9 | 426 | 174.66 | 4 | 34,240,060 | c39, c3-1, c3-2, c85 |
| 10 | 279 | 204.49 | 10 | 32,156,945 | c100, c107, c130, c137, c143, c16, c20, c31, c8, c81 |
| 11 | 263 | 141.72 | 6 | 23,588,853 | c108, c149, c18, c24, c4, c9 |
| 12 | 281 | 148.82 | 11 | 19,016,816 | c145, c173, c184, c25, c27, c273, c54, c70, c74, c75, c99 |
| 13 | 438 | 343.8 | 9 | 37,054,565 | c10, c123, c163, c19, c22, c49, c53, c80, c86 |
| Total | 4,455 | 2,780.03 | 111 | 401,069,438 | |

**Table S19.** Statistics of the assembled 13 chromosomes of *S. tora*

| Chromosome | Anchored contig number | Length of chromosome (bp) |
|---|---|---|
| Chr1 | 21 | 32,816,166 |
| Chr2 | 35 | 42,009,719 |
| Chr3 | 17 | 37,860,065 |
| Chr4 | 39 | 30,689,712 |
| Chr5 | 26 | 52,777,034 |
| Chr6 | 14 | 46,512,068 |
| Chr7 | 17 | 30,975,534 |
| Chr8 | 30 | 49,705,205 |
| Chr9 | 7 | 35,860,388 |
| Chr10 | 21 | 41,499,577 |
| Chr11 | 22 | 30,871,617 |
| Chr12 | 28 | 29,799,933 |
| Chr13 | 13 | 41,270,226 |
| **Total** | 290 | 502,647,244 |

**Table S20.** Statistics of repeat elements in the *S. tora* genome

| Types | Counts | Masked length | Masked (%) |
|---|---|---|---|
| **Retroelements** | **146,719** | **126,200,327** | **23.97** |
| SINEs: | 3,648 | 320,822 | 0.06 |
| ALUs | 258 | 32,943 | 0.01 |
| LINEs: | 36,028 | 17,305,497 | 3.29 |
| CR1 | 33 | 1,657 | 0.00 |
| L1 | 10,974 | 4,629,733 | 0.88 |
| L2 | 8,913 | 9,617,265 | 1.83 |
| RTE-BovB | 16,036 | 3,043,940 | 0.58 |
| Penelope | 61 | 11,083 | 0.00 |
| LTR elements: | 106,503 | 108,574,008 | 20.63 |
| Ty1/Copia | 36,468 | 24,658,248 | 4.68 |
| Ty3/Gypsy | 67,989 | 81,902,798 | 15.56 |
| BEL/Pao | 594 | 732,594 | 0.14 |
| Caulimoviruses | 554 | 1,034,136 | 0.20 |
| **DNA transposons** | **126,655** | **34,116,808** | **6.48** |
| MULE-MuDR | 37,089 | 8,340,270 | 1.58 |
| CMC-EnSpm | 17,286 | 8,174,915 | 1.55 |
| hAT-Ac | 26,300 | 5,529,442 | 1.05 |
| hAT-Tag1 | 11,009 | 2,970,951 | 0.56 |
| hAT-Charlie | 3,450 | 2,231,696 | 0.42 |
| Unclassified: | 325,176 | 111,279,209 | 21.14 |
| **Total interspersed repeats:** | | **261,871,205** | **49.75** |
| Small RNA: | 2,030 | 377,643 | 0.07 |
| Satellites: | 72 | 21,056 | 0.00 |
| Simple repeats: | 191,675 | 13,232,686 | 2.51 |
| **Total** | | **283,551,945** | **53.87** |

**Table S21.** Statistics of the RNA-Seq reads produced by Illumina sequencing platform

| Stage | Sample | Read bases | Reads | GC (%) | Q20 (%) | Q30 (%) |
|---|---|---|---|---|---|---|
| 1 | Stage1-1 | 5,716,321,575 | 44,230,592 | 44.33 | 89.46 | 77.52 |
| | Stage1-2 | 5,416,922,426 | 41,572,424 | 44.31 | 90.71 | 79.20 |
| 2 | Stage2-1 | 5,222,538,241 | 40,494,100 | 45.15 | 90.36 | 79.00 |
| | Stage2-2 | 4,957,068,902 | 38,637,340 | 43.89 | 90.41 | 78.57 |
| 3 | Stage3-1 | 5,849,017,608 | 44,752,688 | 45.23 | 90.52 | 78.95 |
| | Stage3-2 | 5,182,252,082 | 39,833,596 | 45.52 | 90.38 | 78.70 |
| 4 | Stage4-1 | 5,794,527,131 | 44,470,644 | 46.98 | 90.32 | 79.02 |
| | Stage4-2 | 4,900,442,069 | 37,584,824 | 47.75 | 90.71 | 79.78 |
| 5 | Stage5-1 | 5,022,226,735 | 38,394,282 | 47.26 | 90.86 | 79.96 |
| | Stage5-2 | 5,555,427,803 | 42,445,844 | 48.23 | 90.44 | 79.33 |
| 6 | Stage6-1 | 5,171,309,937 | 38,629,298 | 45.82 | 89.61 | 78.35 |
| | Stage6-2 | 5,547,031,298 | 41,408,918 | 46.55 | 90.66 | 78.80 |
| 7 | Stage7-1 | 5,193,556,843 | 38,492,940 | 47.35 | 91.74 | 82.09 |
| | Stage7-2 | 5,148,850,942 | 38,251,900 | 47.13 | 91.67 | 82.09 |

**Table S22.** Primers used for the study of CHS-L (STO07G228250) and CHS (STO03G058250) genes

| Names | Used for | Sequences* (5′-3′) |
|---|---|---|
| Sto07g228250F | Cloning | AAGGATCCATGGAGAGTGCTGGAG |
| Sto07g228250R | Cloning | AACTCGAGCTAGTCTCTCAGAGGG |
| Sto03g058250F | Cloning | GAATTCATGGTGAGTGTGAGTGAGATC |
| Sto03g058250R | Cloning | AAGCTTTTAGTTAACTCCCACACTGCG |

*Underlined sequences represent restriction enzyme sites.

**Supplementary References:**


1. G. C. Allen, M. A. Flores-Vergara, S. Krasynanski, S. Kumar, W. F. Thompson, A modified protocol for rapid DNA isolation from plant tissues using cetyltrimethylammonium bromide. *Nat. Protoc.* **1**, 2320-2325 (2006).
2. A. M. Bolger, M. Lohse, B. Usadel, Trimmomatic: a flexible trimmer for Illumina sequence data. *Bioinformatics* **30**, 2114-2120 (2014).
3. S.-H. Kang, S. Y. Won, C.-K. Kim, The complete mitochondrial genome sequences of *Senna tora* (Fabales: Fabaceae). *Mitochondrial DNA B* **4**, 1283-1284 (2019).
4. G.-H. Shin, Y. Shin, M. Jung, J.-m. Hong, S. Lee, S. Subramaniyam, E.-S. Noh, E.-H. Shin, E.-H. Park, J. Y. Park, Y.-O. Kim, K.-M. Choi, B.-H. Nam, C.-I. Park, First Draft Genome for Red Sea Bream of Family Sparidae. *Front. Genet.* **9**, (2018).
5. G. Marçais, C. Kingsford, A fast, lock-free approach for efficient parallel counting of occurrences of k-mers. *Bioinformatics* **27**, 764-770 (2011).
6. R. Luo, B. Liu, Y. Xie, Z. Li, W. Huang, J. Yuan, G. He, Y. Chen, Q. Pan, Y. Liu, J. Tang, G. Wu, H. Zhang, Y. Shi, Y. Liu, C. Yu, B. Wang, Y. Lu, C. Han, D. W. Cheung, S.-M. Yiu, S. Peng, Z. Xiaoqian, G. Liu, X. Liao, Y. Li, H. Yang, J. Wang, T.-W. Lam, J. Wang, SOAPdenovo2: an empirically improved memory-efficient short-read *de novo* assembler. *Gigascience* **1**, (2012).
7. J. Butler, I. MacCallum, M. Kleber, I. A. Shlyakhter, M. K. Belmonte, E. S. Lander, C. Nusbaum, D. B. Jaffe, ALLPATHS: *de novo* assembly of whole-genome shotgun microreads. *Genome Res.* **18**, 810-820 (2008).
8. I. MacCallum, D. Przybylski, S. Gnerre, J. Burton, I. Shlyakhter, A. Gnirke, J. Malek, K. McKernan, S. Ranade, T. P. Shea, L. Williams, S. Young, C. Nusbaum, D. B. Jaffe, ALLPATHS 2: small genomes assembled accurately and with high continuity from short paired reads. *Genome Biol.* **10**, R103 (2009).
9. R. Kajitani, K. Toshimoto, H. Noguchi, A. Toyoda, Y. Ogura, M. Okuno, M. Yabana, M. Harada, E. Nagayasu, H. Maruyama, Y. Kohara, A. Fujiyama, T. Hayashi, T. Itoh, Efficient de novo assembly of highly heterozygous genomes from whole-genome shotgun short reads. *Genome Res.* **24**, 1384-1395 (2014).
10. C.-S. Chin, P. Peluso, F. J. Sedlazeck, M. Nattestad, G. T. Concepcion, A. Clum, C. Dunn, R. O'Malley, R. Figueroa-Balderas, A. Morales-Cruz, G. R. Cramer, M. Delledonne, C. Luo, J. R. Ecker, D. Cantu, D. R. Rank, M. C. Schatz, Phased diploid genome assembly with single-molecule real-time sequencing. *Nat. Methods* **13**, 1050-1054 (2016).
11. M. A. DePristo, E. Banks, R. Poplin, K. V. Garimella, J. R. Maguire, C. Hartl, A. A. Philippakis, G. del Angel, M. A. Rivas, M. Hanna, A. McKenna, T. J. Fennell, A. M. Kernytsky, A. Y. Sivachenko, K. Cibulskis, S. B. Gabriel, D. Altshuler, M. J. Daly, A framework for variation discovery and genotyping using next-generation DNA sequencing data. *Nat. Genet.* **43**, 491-498 (2011).
12. F. A. Simão, R. M. Waterhouse, P. Ioannidis, E. V. Kriventseva, E. M. Zdobnov, BUSCO: assessing genome assembly and annotation completeness with single-copy orthologs. *Bioinformatics* **31**, 3210-3212 (2015).
13. T. Kaczorowski, W. Szybalski, Genomic DNA sequencing by SPEL-6 primer walking using hexamer ligation1Published in conjunction with A Wisconsin Gathering Honoring Waclaw Szybalski on the occasion of his 75th year and 20years of Editorship-in-Chief of Gene, 10–11 August 1997, University of Wisconsin, Madison, WI, USA.1. *Gene* **223**, 83-91 (1998).
14. W. J. Kent, BLAT--the BLAST-like alignment tool. *Genome Res.* **12**, 656-664

(2002).
15. R. J. Elshire, J. C. Glaubitz, Q. Sun, J. A. Poland, K. Kawamoto, E. S. Buckler, S. E. Mitchell, A robust, simple genotyping-by-sequencing (GBS) approach for high diversity species. *PLoS One* **6**, e19379-e19379 (2011).
16. H. Li, R. Durbin, Fast and accurate long-read alignment with Burrows-Wheeler transform. *Bioinformatics* **26**, 589-595 (2010).
17. P. Danecek, A. Auton, G. Abecasis, C. A. Albers, E. Banks, M. A. DePristo, R. E. Handsaker, G. Lunter, G. T. Marth, S. T. Sherry, G. McVean, R. Durbin, G. Genomes Project Analysis, The variant call format and VCFtools. *Bioinformatics* **27**, 2156-2158 (2011).
18. L. Meng, H. Li, L. Zhang, J. Wang, QTL IciMapping: Integrated software for genetic linkage map construction and quantitative trait locus mapping in biparental populations. *Crop J.* **3**, 269-283 (2015).
19. E. Lieberman-Aiden, N. L. van Berkum, L. Williams, M. Imakaev, T. Ragoczy, A. Telling, I. Amit, B. R. Lajoie, P. J. Sabo, M. O. Dorschner, R. Sandstrom, B. Bernstein, M. A. Bender, M. Groudine, A. Gnirke, J. Stamatoyannopoulos, L. A. Mirny, E. S. Lander, J. Dekker, Comprehensive Mapping of Long-Range Interactions Reveals Folding Principles of the Human Genome. *Science* **326**, 289-293 (2009).
20. G. G. Faust, I. M. Hall, SAMBLASTER: fast duplicate marking and structural variant read extraction. *Bioinformatics* **30**, 2503-2505 (2014).
21. H. Li, B. Handsaker, A. Wysoker, T. Fennell, J. Ruan, N. Homer, G. Marth, G. Abecasis, R. Durbin, S. Genome Project Data Processing, The Sequence Alignment/Map format and SAMtools. *Bioinformatics* **25**, 2078-2079 (2009).
22. D. M. Bickhart, B. D. Rosen, S. Koren, B. L. Sayre, A. R. Hastie, S. Chan, J. Lee, E. T. Lam, I. Liachko, S. T. Sullivan, J. N. Burton, H. J. Huson, J. C. Nystrom, C. M. Kelley, J. L. Hutchison, Y. Zhou, J. Sun, A. Crisà, F. A. Ponce de León, J. C. Schwartz, J. A. Hammond, G. C. Waldbieser, S. G. Schroeder, G. E. Liu, M. J. Dunham, J. Shendure, T. S. Sonstegard, A. M. Phillippy, C. P. Van Tassell, T. P. L. Smith, Single-molecule sequencing and chromatin conformation capture enable *de novo* reference assembly of the domestic goat genome. *Nat. Genet.* **49**, 643-650 (2017).
23. J. N. Burton, A. Adey, R. P. Patwardhan, R. Qiu, J. O. Kitzman, J. Shendure, Chromosome-scale scaffolding of *de novo* genome assemblies based on chromatin interactions. *Nat. Biotechnol.* **31**, 1119-1125 (2013).
24. N. C. Durand, J. T. Robinson, M. S. Shamim, I. Machol, J. P. Mesirov, E. S. Lander, E. L. Aiden, Juicebox Provides a Visualization System for Hi-C Contact Maps with Unlimited Zoom. *Cell Syst.* **3**, 99-101 (2016).
25. S. S. P. Rao, M. H. Huntley, N. C. Durand, E. K. Stamenova, I. D. Bochkov, J. T. Robinson, A. L. Sanborn, I. Machol, A. D. Omer, E. S. Lander, E. L. Aiden, A 3D map of the human genome at kilobase resolution reveals principles of chromatin looping. *Cell* **159**, 1665-1680 (2014).
26. Z. Bao, S. R. Eddy, Automated *de novo* identification of repeat sequence families in sequenced genomes. *Genome Res.* **12**, 1269-1276 (2002).
27. A. L. Price, N. C. Jones, P. A. Pevzner, *De novo* identification of repeat families in large genomes. *Bioinformatics* **21 Suppl 1**, i351-i358 (2005).
28. G. Benson, Tandem repeats finder: a program to analyze DNA sequences. *Nucleic Acids Res.* **27**, 573-580 (1999).
29. W. Bao, K. K. Kojima, O. Kohany, Repbase Update, a database of repetitive elements in eukaryotic genomes. *Mob. DNA* **6**, 11-11 (2015).

30. R. K. Varshney, W. Chen, Y. Li, A. K. Bharti, R. K. Saxena, J. A. Schlueter, M. T. A. Donoghue, S. Azam, G. Fan, A. M. Whaley, A. D. Farmer, J. Sheridan, A. Iwata, R. Tuteja, R. V. Penmetsa, W. Wu, H. D. Upadhyaya, S.-P. Yang, T. Shah, K. B. Saxena, T. Michael, W. R. McCombie, B. Yang, G. Zhang, H. Yang, J. Wang, C. Spillane, D. R. Cook, G. D. May, X. Xu, S. A. Jackson, Draft genome sequence of pigeonpea (*Cajanus cajan*), an orphan legume crop of resource-poor farmers. *Nat. Biotechnol.* **30**, 83-89 (2012).
31. S.-H. Kang, J.-Y. Lee, T.-H. Lee, S.-Y. Park, C.-K. Kim, *De novo* transcriptome assembly of the Chinese pearl barley, adlay, by full-length isoform and short-read RNA sequencing. *PLoS One* **13**, e0208344-e0208344 (2018).
32. R. K. Varshney, C. Song, R. K. Saxena, S. Azam, S. Yu, A. G. Sharpe, S. Cannon, J. Baek, B. D. Rosen, B. Tar'an, T. Millan, X. Zhang, L. D. Ramsay, A. Iwata, Y. Wang, W. Nelson, A. D. Farmer, P. M. Gaur, C. Soderlund, R. V. Penmetsa, C. Xu, A. K. Bharti, W. He, P. Winter, S. Zhao, J. K. Hane, N. Carrasquilla-Garcia, J. A. Condie, H. D. Upadhyaya, M.-C. Luo, M. Thudi, C. L. L. Gowda, N. P. Singh, J. Lichtenzveig, K. K. Gali, J. Rubio, N. Nadarajan, J. Dolezel, K. C. Bansal, X. Xu, D. Edwards, G. Zhang, G. Kahl, J. Gil, K. B. Singh, S. K. Datta, S. A. Jackson, J. Wang, D. R. Cook, Draft genome sequence of chickpea (*Cicer arietinum*) provides a resource for trait improvement. *Nat. Biotechnol.* **31**, 240-246 (2013).
33. S.-H. Kang, W.-H. Lee, C.-M. Lee, J.-S. Sim, S. Y. Won, S.-R. Han, S.-J. Kwon, J. S. Kim, C.-K. Kim, T.-J. Oh, *De novo* transcriptome sequence of *Senna tora* provides insights into anthraquinone biosynthesis. *bioRxiv*, 837385 (2019).
34. S. Kim, M. Park, S.-I. Yeom, Y.-M. Kim, J. M. Lee, H.-A. Lee, E. Seo, J. Choi, K. Cheong, K.-T. Kim, K. Jung, G.-W. Lee, S.-K. Oh, C. Bae, S.-B. Kim, H.-Y. Lee, S.-Y. Kim, M.-S. Kim, B.-C. Kang, Y. D. Jo, H.-B. Yang, H.-J. Jeong, W.-H. Kang, J.-K. Kwon, C. Shin, J. Y. Lim, J. H. Park, J. H. Huh, J.-S. Kim, B.-D. Kim, O. Cohen, I. Paran, M. C. Suh, S. B. Lee, Y.-K. Kim, Y. Shin, S.-J. Noh, J. Park, Y. S. Seo, S.-Y. Kwon, H. A. Kim, J. M. Park, H.-J. Kim, S.-B. Choi, P. W. Bosland, G. Reeves, S.-H. Jo, B.-W. Lee, H.-T. Cho, H.-S. Choi, M.-S. Lee, Y. Yu, Y. Do Choi, B.-S. Park, A. van Deynze, H. Ashrafi, T. Hill, W. T. Kim, H.-S. Pai, H. K. Ahn, I. Yeam, J. J. Giovannoni, J. K. C. Rose, I. Sørensen, S.-J. Lee, R. W. Kim, I.-Y. Choi, B.-S. Choi, J.-S. Lim, Y.-H. Lee, D. Choi, Genome sequence of the hot pepper provides insights into the evolution of pungency in Capsicum species. *Nat. Genet.* **46**, 270-278 (2014).
35. B.-H. Nam, W. Kwak, Y.-O. Kim, D.-G. Kim, H. J. Kong, W.-J. Kim, J.-H. Kang, J. Y. Park, C. M. An, J.-Y. Moon, C. J. Park, J. W. Yu, J. Yoon, M. Seo, K. Kim, D. K. Kim, S. Lee, S. Sung, C. Lee, Y. Shin, M. Jung, B.-C. Kang, G.-H. Shin, S. Ka, K. Caetano-Anolles, S. Cho, H. Kim, Genome sequence of pacific abalone (*Haliotis discus hannai*): the first draft genome in family Haliotidae. *Gigascience* **6**, 1-8 (2017).
36. C. Trapnell, A. Roberts, L. Goff, G. Pertea, D. Kim, D. R. Kelley, H. Pimentel, S. L. Salzberg, J. L. Rinn, L. Pachter, Differential gene and transcript expression analysis of RNA-seq experiments with TopHat and Cufflinks. *Nat. Protocols* **7**, 562-578 (2012).
37. B. J. Haas, A. L. Delcher, S. M. Mount, J. R. Wortman, R. K. Smith, Jr., L. I. Hannick, R. Maiti, C. M. Ronning, D. B. Rusch, C. D. Town, S. L. Salzberg, O. White, Improving the Arabidopsis genome annotation using maximal transcript alignment assemblies. *Nucleic Acids Res.* **31**, 5654-5666 (2003).
38. M. Stanke, O. Schöffmann, B. Morgenstern, S. Waack, Gene prediction in

eukaryotes with a generalized hidden Markov model that uses hints from external sources. *BMC Bioinformatics* **7**, 62-62 (2006).

39. E. Blanco, G. Parra, R. Guigó, Using geneid to identify genes. *Curr. Protoc. Bioinformatics* **Chapter 4**, 4.3 (2007).
40. G. S. C. Slater, E. Birney, Automated generation of heuristics for biological sequence comparison. *BMC Bioinformatics* **6**, 31-31 (2005).
41. S. Götz, J. M. García-Gómez, J. Terol, T. D. Williams, S. H. Nagaraj, M. J. Nueda, M. Robles, M. Talón, J. Dopazo, A. Conesa, High-throughput functional annotation and data mining with the Blast2GO suite. *Nucleic Acids Res.* **36**, 3420-3435 (2008).
42. P. Jones, D. Binns, H.-Y. Chang, M. Fraser, W. Li, C. McAnulla, H. McWilliam, J. Maslen, A. Mitchell, G. Nuka, S. Pesseat, A. F. Quinn, A. Sangrador-Vegas, M. Scheremetjew, S.-Y. Yong, R. Lopez, S. Hunter, InterProScan 5: genome-scale protein function classification. *Bioinformatics* **30**, 1236-1240 (2014).
43. M. Jayakodi, J. W. Jung, D. Park, Y.-J. Ahn, S.-C. Lee, S.-Y. Shin, C. Shin, T.-J. Yang, H. W. Kwon, Genome-wide characterization of long intergenic non-coding RNAs (lincRNAs) provides new insight into viral diseases in honey bees *Apis cerana* and *Apis mellifera*. *BMC Genomics* **16**, 680 (2015).
44. T. U. Consortium, UniProt: a worldwide hub of protein knowledge. *Nucleic Acids Res.* **47**, D506-D515 (2018).
45. S. El-Gebali, J. Mistry, A. Bateman, S. R. Eddy, A. Luciani, S. C. Potter, M. Qureshi, L. J. Richardson, G. A. Salazar, A. Smart, E. L L. Sonnhammer, L. Hirsh, L. Paladin, D. Piovesan, S. C E. Tosatto, R. D. Finn, The Pfam protein families database in 2019. *Nucleic Acids Res.* **47**, D427-D432 (2018).
46. L. Kong, Y. Zhang, Z.-Q. Ye, X.-Q. Liu, S.-Q. Zhao, L. Wei, G. Gao, CPC: assess the protein-coding potential of transcripts using sequence features and support vector machine. *Nucleic Acids Res.* **35**, W345-W349 (2007).
47. The Rnacentral Consortium, RNAcentral: a hub of information for non-coding RNA sequences. *Nucleic Acids Res.* **47**, D1250-D1251 (2019).
48. L. Li, C. J. Stoeckert, Jr., D. S. Roos, OrthoMCL: identification of ortholog groups for eukaryotic genomes. *Genome Res.* **13**, 2178-2189 (2003).
49. K. Katoh, D. M. Standley, MAFFT multiple sequence alignment software version 7: improvements in performance and usability. *Mol. Biol. Evol.* **30**, 772-780 (2013).
50. J. Castresana, Selection of conserved blocks from multiple alignments for their use in phylogenetic analysis. *Mol. Biol. Evol.* **17**, 540-552 (2000).
51. L.-T. Nguyen, H. A. Schmidt, A. von Haeseler, B. Q. Minh, IQ-TREE: a fast and effective stochastic algorithm for estimating maximum-likelihood phylogenies. *Mol. Biol. Evol.* **32**, 268-274 (2015).
52. M. V. Han, G. W. C. Thomas, J. Lugo-Martinez, M. W. Hahn, Estimating gene gain and loss rates in the presence of error in genome assembly and annotation using CAFE 3. *Mol. Biol. Evol.* **30**, 1987-1997 (2013).
53. R. Bouckaert, T. G. Vaughan, J. Barido-Sottani, S. Duchêne, M. Fourment, A. Gavryushkina, J. Heled, G. Jones, D. Kühnert, N. De Maio, M. Matschiner, F. K. Mendes, N. F. Müller, H. A. Ogilvie, L. du Plessis, A. Popinga, A. Rambaut, D. Rasmussen, I. Siveroni, M. A. Suchard, C.-H. Wu, D. Xie, C. Zhang, T. Stadler, A. J. Drummond, BEAST 2.5: An advanced software platform for Bayesian evolutionary analysis. *PLoS Comput. Biol.* **15**, e1006650-e1006650 (2019).
54. Z. Zhang, J. Xiao, J. Wu, H. Zhang, G. Liu, X. Wang, L. Dai, ParaAT: a parallel

tool for constructing multiple protein-coding DNA alignments. *Biochem. Biophys. Res. Commun.* **419**, 779-781 (2012).

55. Z. Yang, PAML 4: Phylogenetic Analysis by Maximum Likelihood. *Mol. Biol. Evol.* **24**, 1586-1591 (2007).
56. Y. Van de Peer, A mystery unveiled. *Genome Biol.* **12**, 113 (2011).
57. M. Freeling, B. C. Thomas, Gene-balanced duplications, like tetraploidy, provide predictable drive to increase morphological complexity. *Genome Res.* **16**, 805-814 (2006).
58. S. P. Otto, The evolutionary consequences of polyploidy. *Cell* **131**, 452-462 (2007).
59. M. E. Schranz, S. Mohammadin, P. P. Edger, Ancient whole genome duplications, novelty and diversification: the WGD Radiation Lag-Time Model. *Curr. Opin. Plant Biol.* **15**, 147-153 (2012).
60. J. Schmutz, S. B. Cannon, J. Schlueter, J. Ma, T. Mitros, W. Nelson, D. L. Hyten, Q. Song, J. J. Thelen, J. Cheng, D. Xu, U. Hellsten, G. D. May, Y. Yu, T. Sakurai, T. Umezawa, M. K. Bhattacharyya, D. Sandhu, B. Valliyodan, E. Lindquist, M. Peto, D. Grant, S. Shu, D. Goodstein, K. Barry, M. Futrell-Griggs, B. Abernathy, J. Du, Z. Tian, L. Zhu, N. Gill, T. Joshi, M. Libault, A. Sethuraman, X.-C. Zhang, K. Shinozaki, H. T. Nguyen, R. A. Wing, P. Cregan, J. Specht, J. Grimwood, D. Rokhsar, G. Stacey, R. C. Shoemaker, S. A. Jackson, Genome sequence of the palaeopolyploid soybean. *Nature* **463**, 178-183 (2010).
61. A. Bruneau, M. Mercure, G. P. Lewis, P. S. Herendeen, Phylogenetic patterns and diversification in the caesalpinioid legumes This paper is one of a selection of papers published in the Special Issue on Systematics Research. *Botany* **86**, 697-718 (2008).
62. T. Soga, Y. Ohashi, Y. Ueno, H. Naraoka, M. Tomita, T. Nishioka, Quantitative Metabolome Analysis Using Capillary Electrophoresis Mass Spectrometry. *J. Proteome Res.* **2**, 488-494 (2003).
63. A. Dobin, C. A. Davis, F. Schlesinger, J. Drenkow, C. Zaleski, S. Jha, P. Batut, M. Chaisson, T. R. Gingeras, STAR: ultrafast universal RNA-seq aligner. *Bioinformatics* **29**, 15-21 (2013).
64. B. Li, C. N. Dewey, RSEM: accurate transcript quantification from RNA-Seq data with or without a reference genome. *BMC Bioinformatics* **12**, 323-323 (2011).
65. M. D. Robinson, D. J. McCarthy, G. K. Smyth, edgeR: a Bioconductor package for differential expression analysis of digital gene expression data. *Bioinformatics* **26**, 139-140 (2010).
66. D. Kim, M. Jung, I. J. Ha, M. Y. Lee, S.-G. Lee, Y. Shin, S. Subramaniyam, J. Oh, Transcriptional profiles of secondary metabolite biosynthesis genes and cytochromes in the leaves of four papaver species. *Data* **3**, 55 (2018).
67. T. Pluskal, M. P. Torrens-Spence, T. R. Fallon, A. De Abreu, C. H. Shi, J.-K. Weng, The biosynthetic origin of psychoactive kavalactones in kava. *Nat. Plants* **5**, 867-878 (2019).
68. D. T. Jones, W. R. Taylor, J. M. Thornton, The rapid generation of mutation data matrices from protein sequences. *Bioinformatics* **8**, 275-282 (1992).
69. S. Kumar, G. Stecher, M. Li, C. Knyaz, K. Tamura, MEGA X: Molecular Evolutionary Genetics Analysis across Computing Platforms. *Mol. Biol. Evol.* **35**, 1547-1549 (2018).
